# A tribenzamide small molecule with retinoic acid-like activity promotes axon growth and motor recovery after spinal cord injury

**DOI:** 10.1101/2025.11.05.686894

**Authors:** Katha Sanyal, Dhanuush Balakannan, Sk Rajibul Haque, Rutuja Arun Pendharkar, Prakash Chermakani, Manojkumar Kumaran, Pratikhya Acharya, Debabrata Maity, Ishwariya Venkatesh

## Abstract

Injury to the adult central nervous system triggers minimal axonal regeneration, partly due to the limited activation of intrinsic growth programs. Retinoic acid (RA) signaling has been shown to promote modest regenerative responses, but its clinical utility is restricted by poor biochemical stability and short-lived receptor engagement. Here, we report a tribenzamide-based small molecule DM04, that recapitulates RA-like transcriptional and phenotypic effects and enhances regenerative outcomes — independent of canonical RARE-dependent transactivation. DM04 promoted neurite outgrowth in primary neurons, upregulated canonical RA-responsive genes, and supported neural induction from human iPSCs. In a murine spinal injury model, DM04 treatment improved motor recovery. Transcriptomic analysis revealed shared target gene activation between RA and DM04, along with unique enrichment of extracellular matrix remodeling pathways. These findings establish DM04 as a small molecule based approach with dual efficacy in injury and developmental contexts, acting through a mechanism distinct from canonical RAR transactivation, and highlight its promise as a candidate for further preclinical investigation in neural repair.

## Introduction

The adult mammalian central nervous system (CNS) exhibits a limited capacity for repair following injury, largely due to the failure of mature neurons to initiate intrinsic growth programs^1–8^ and the presence of a non-permissive extracellular environment^9,10^. Although a number of strategies have been tested to stimulate regeneration, including the use of transcription factors^11^, growth factors, and physical scaffolds, very few interventions have led to robust and reproducible axon regrowth, synaptic reformation, or complete behavioral recovery *in vivo*. Retinoic acid (RA), a small lipophilic molecule involved in development and neuronal plasticity^12–16^, has shown promise in models of regeneration due to its ability to modulate gene expression through nuclear receptors and trigger outgrowth programs^15^. However, RA’s use is limited by its poor chemical stability, that is, its intrinsic susceptibility to isomerisation and oxidation, and its rapid metabolic clearance via autoinductive catabolism under physiological conditions, leading to short-lived bioactivity and challenging dose control. Throughout this study we distinguish three properties that are often conflated: chemical stability (resistance of the free compound to degradation such as oxidation or photoisomerisation), metabolic stability (resistance to enzymatic clearance in a biological system), and receptor-complex stability (the persistence of a ligand-receptor complex once formed). These are independent properties, and evidence for one does not imply the others^17^. RA mediates its biological effects primarily through retinoic acid receptors (RARs), among which RARβ has been implicated in axon growth, neuronal differentiation, and regenerative signaling^18–19,56–58^. Nevertheless, transcription factor overexpression and combinatorial screening studies indicate that RARβ activity alone is insufficient to elicit robust regenerative growth or functional recovery in the adult CNS, underscoring the need for alternative approaches to modulate retinoid signaling more effectively^1^.

Together, the inherent instability of RA and the limited efficacy of direct RARβ modulation motivated the development of alternate small molecules, designed to resist the chemical degradation that limits RA, capable of engaging retinoid-responsive/similar regenerative pathways. The chemical structure of RA is amphiphilic in nature. The hydrophobic part consists of a trimethyl-substituted cyclohexene ring attached to a long conjugated tetraene side chain. This part is connected to a polar acidic end group. Chemically stable hydrophobic oligo-arylamides have been successfully shown to modulate different biomolecular interactions^20–22^. We rationalised that a hydrophobic oligo-arylamide with a terminal polar acidic group will be interesting to study in place of RA. In this study, we demonstrated a tribenzamide based small molecule, DM04, that exhibits RA-like activity in both injury and differentiation contexts.

Transcriptomic analysis revealed that DM04 treatment upregulated a core set of genes known to be downstream of RA signaling, many of which are associated with neurite outgrowth, ECM remodeling, and axonal regeneration. When tested in a mouse spinal cord injury model, DM04 treatment increased axonal re-growth at the lesion border and improved behavioral coordination. Parallel experiments in human induced pluripotent stem cells (hiPSCs) demonstrated that DM04 could substitute for RA in neural induction protocols, promoting the formation of Pax6-positive neural progenitor cells. These convergent findings from two model systems injury and development suggest that DM04 has robust neurogenic and regenerative potential.

Small-molecule modulation of endogenous signaling pathways offers a promising approach to promote neural regeneration, with practical advantages over viral gene delivery strategies, including controllable dosing and reversibility, which may facilitate future preclinical development. Retinoic acid (RA) signaling plays a well-established role in neuronal differentiation and neurite outgrowth, yet pharmacological strategies to harness this pathway for CNS repair remain limited. In this study, we investigated the tribenzamide compound DM04 as a potential modulator of RA signaling to assess whether a degradation-resistant RA-mimetic can enhance neurite growth in neuronal models.

## Results

### Rational Design and Computational Characterization of DM04

RARβ has previously been implicated in promoting modest axonal sprouting following spinal cord injury, largely through partial reactivation of growth-associated transcriptional programs^1,38^. However, the regenerative effects observed in these contexts are typically limited and short-lived, raising the possibility that intrinsic features of the receptor–ligand interaction may constrain its therapeutic potential. One plausible contributing factor is the biochemical instability of retinoic acid (RA), the endogenous ligand for RARβ. RA is well known to degrade rapidly through isomerization and oxidation under physiological conditions, which could limit its availability and reduce the duration of receptor activation^17,39^. This in turn may restrict the ability of RARβ to sustain the transcriptional programs required for cytoskeletal remodeling and axon extension after injury.

We hypothesized that an alternate stable small-molecule based ligand capable of engaging the receptor more effectively than RA could prolong receptor occupancy, enhance transcriptional output, and thereby amplify regenerative signaling. To explore this possibility, we designed DM04, a tribenzamide based small molecule having a terminal acidic group, which is expected to resist chemical degradation and, as predicted by modeling, potentially engage the ligand-binding pocket of RARβ in a more stable configuration (a receptor-complex property distinct from the compound’s own chemical stability). The tribenzamide part mimics the hydrophobic part of RA, which consists of a lengthy conjugated tetraene side chain connected to a trimethyl-substituted cyclohexene ring.

Docking simulations supported our hypothesis. Compared to RA, which appeared to adopt a relatively shallow and flexible pose within the RARβ pocket, DM04 engaged deeper residues and formed additional stabilizing interactions, including hydrogen bonds and hydrophobic contacts (Figure 1B, Supplementary Figure S1D). These interactions may help to reinforce the receptor’s active conformation and reduce conformational fluctuations that would otherwise lead to dissociation. Structural overlays suggested that DM04 may reduce solvent exposure and promote tighter packing within the ligand-binding domain, consistent with a more stable receptor–ligand complex in silico. We note this is a computational prediction of receptor-complex stability and was not confirmed by direct binding measurements.

**Figure 1.**
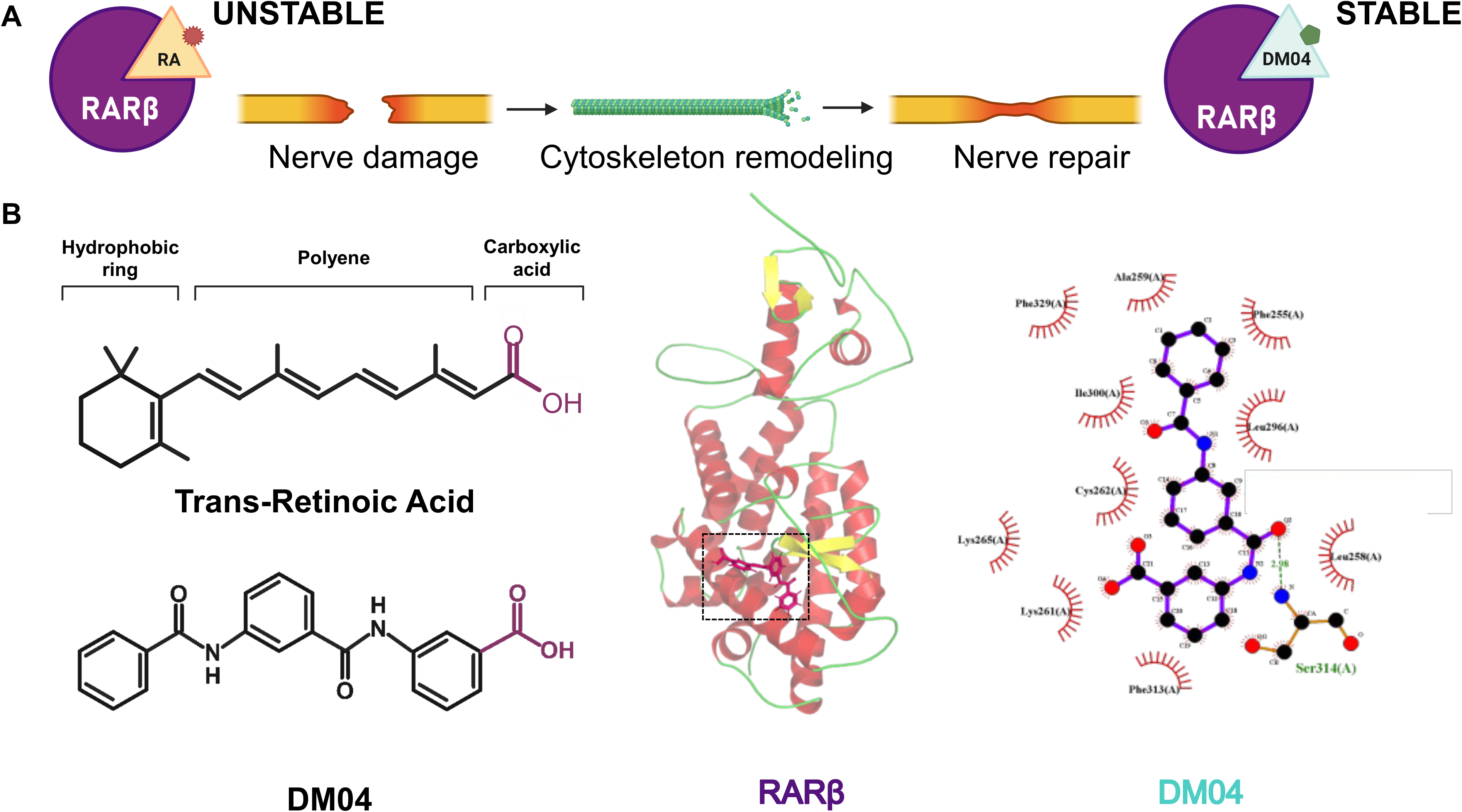
Rational Design and Computational Characterization of DM04. (A) Schematic representation of the proposed relationship between RARβ-associated signaling and neuronal repair. (B) Chemical structures of trans-retinoic acid and DM04. Structural representation of the modeled RARβ–DM04 complex, highlighting predicted interactions within the ligand-binding pocket. The docking model represents a structural hypothesis for potential RARβ engagement and does not establish receptor binding or receptor-complex stabilization.

Homology modeling of mouse RARβ yielded a highly refined tertiary structure characterized by the canonical alpha-helical sandwich architecture typical of nuclear receptor LBDs. Stereochemical evaluation confirmed excellent geometry, with 96.2% of residues in the core Ramachandran favored region and zero disallowed steric clashes.

Curvature-based cavity mapping identified five potential binding pockets on the RARβ receptor surface. Cavity 1 corresponded to the primary canonical ligand-binding pocket, possessing a voluminous cavity of 1595 Å³. Across all docked compounds, Cavity 1 yielded the highest binding affinities. Remarkably, compound DM04 exhibited the strongest binding affinity to RARβ with a Vina score of -12.2 kcal/mol, outperforming the endogenous agonist all-trans retinoic acid (9CR_RET, -11.9 kcal/mol). The remaining DM analogs are bound with descending affinities: DM03 (-11.5 kcal/mol), DM01 (-10.7 kcal/mol), and DM02 (-10.1 kcal/mol). DM04’s superior affinity arises from its extended multi-ring aromatic core, which establishes extensive hydrophobic contacts and aromatic pi-stacking within the lipophilic pocket of RARβ, supplemented by polar anchoring to conserved residues (Table 1).

**Table 1.** Molecular Docking Binding Affinities of Tested Compounds Against RARβ.

| Ligand ID | Chemical / Common Name | Binding Affinity (kcal/mol) |
| --- | --- | --- |
| **DM04** | 3-(3-benzamidobenzamido)benzoic acid | **-12.2** |
| **9CR_RET** | All-trans Retinoic Acid (Endogenous Agonist) | -11.9 |
| **DM03** | Methyl 3-(3-benzamidobenzamido)benzoate | -11.5 |
| **DM01** | 3-benzamidobenzoic acid | -10.7 |
| **DM02** | 3-(3-nitrobenzamido)benzoic acid | -10.1 |

**Table 2.**
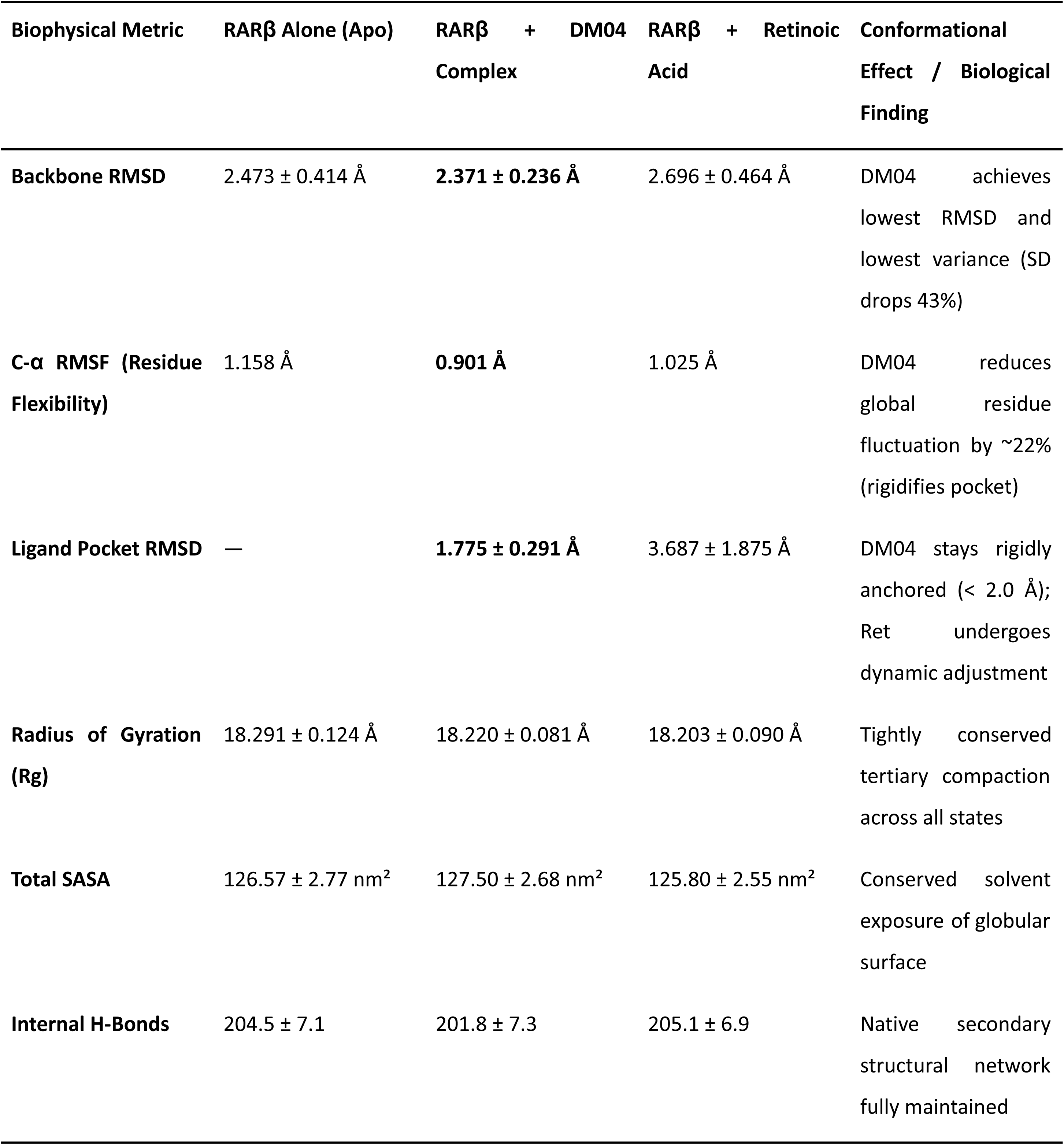
Quantitative Biophysical Parameters from 100 ns Molecular Dynamics Trajectories.

This was supported by 100 ns molecular dynamics simulations, in which the RARβ–DM04 complex exhibited a lower and less variable backbone RMSD (2.37 ± 0.24 Å) than the apo receptor (2.47 ± 0.41 Å) or the RARβ–RA complex (2.70 ± 0.46 Å), a ∼22% reduction in Cα RMSF relative to apo RARβ, and a stable ligand pose throughout the trajectory (RMSD 1.78 ± 0.29 Å), in contrast to RA, which sampled a broader range of poses within the pocket (RMSD 3.69 ± 1.88 Å; Supplementary Figure S2). Docking replicates additionally showed DM04 binding RARβ with higher affinity (−12.2 kcal/mol) than the endogenous agonist RA (−11.9 kcal/mol) (Table 1)

We synthesized a benzamide library in a straightforward synthetic way starting from commercially available 3-nitrobenzoic acid (Scheme 1). 3-Nitrobenzoic acid is esterified and reduced to provide methyl 3-aminobenzoate. Subsequently, the chain elongation of the benzamides was achieved using iterative amide coupling between methyl 3-aminobenzoate and different benzoic acids using Mukaiyama’s reagent, followed by reduction of the nitro groups (whenever necessary). Finally, the ester group was saponified to provide the final acid compound. DM04 showed resistance to degradation under selected stress conditions, including UV irradiation and oxidative treatment, conditions that are known to affect retinoic acid stability^81,82^. Following UV irradiation (360 nm) for up to 60 min, DM04 purity remained unchanged by HPLC analysis (Supplementary Figure S3). Similarly, incubation with H₂O₂ (1 mM) for 3 h resulted in >97% recovery of the compound, consistent with resistance to oxidative degradation under the conditions tested (Supplementary Figure S4). These experiments provide evidence of DM04 persistence under the specific stress conditions examined but do not constitute a direct matched comparison of the chemical stability of DM04 and all-trans retinoic acid. Accordingly, we interpret these findings as evidence of resistance to selected chemical stressors rather than as a quantitative demonstration of superior chemical stability relative to RA. Given that this hydrophobic–polar balance could in principle promote self-association, we next assessed whether DM04 aggregates at the concentrations used in our biological assays. DM04 was fully soluble up to 20 mM in DMSO with no visible precipitation or turbidity. To assess aggregation propensity at working concentrations, we recorded UV-Vis absorbance and dynamic light scattering (DLS) across a 5–25 μM concentration range in water. The absorbance band at ∼256 nm increased proportionally with concentration with no shift in λmax (Supplementary Figure S5), and DLS showed no evidence of large aggregate formation across this range (Supplementary Figure S6), indicating that DM04 does not aggregate at the concentrations used in this study.

To directly test whether DM04 drives RARE-dependent transcriptional activation as predicted by the docking model, we performed a RARE-luciferase reporter assay (pGL3-RARE-Luc) across a 48-hour time course (Supplementary Figure S7, Table S2). RA induced robust, time-dependent reporter activation, reaching an approximately 24-fold increase over vehicles by 24–48 hours, consistent with canonical RAR-mediated transactivation. In contrast, DM04 failed to activate the RARE reporter at any timepoint tested (log2FC ≈ 0.3–0.8), despite the favorable docking pose predicted above. This result indicates that, it does not drive canonical RARE-dependent transcriptional output in the way RA does. We therefore consider the docking-predicted stabilization model in Figure 1 to represent a structural hypothesis rather than a confirmed mechanism of transactivation, and revisit alternative explanations for DM04’s RA - like phenotypic and transcriptional effects.

**Scheme 1.**
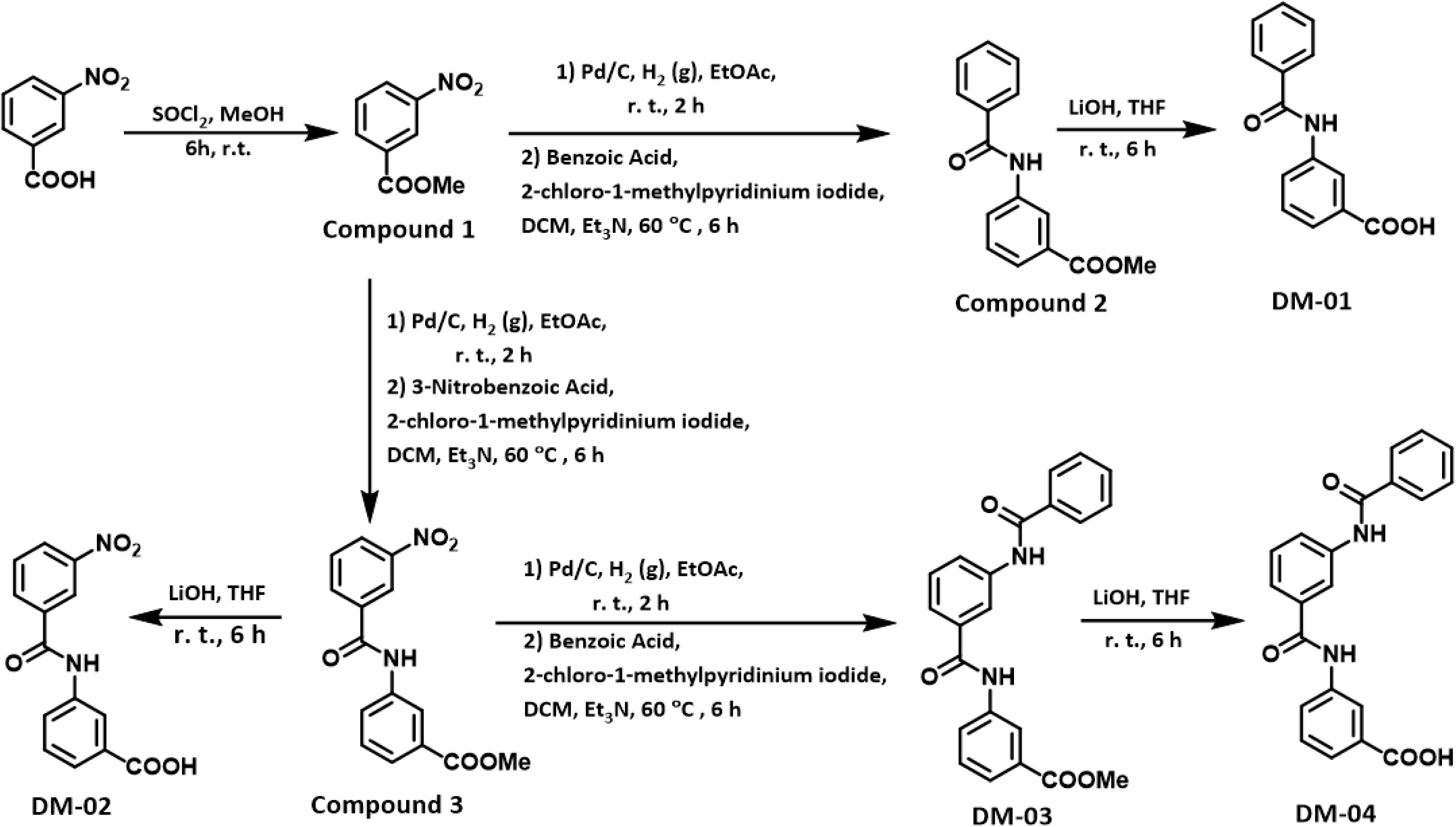
Synthesis of DM01-DM04.

### DM04 is well tolerated by neuronal cells and does not compromise viability across a broad concentration range

To determine the cellular tolerability of DM04, we first assessed its cytotoxicity in Neuro2a cells treated across a concentration gradient (10–100 μM). The experimental workflow is outlined in Figure 2A: cells were expanded for three days, plated into 96-well plates, and treated on Day 4 with DMSO (vehicle), DM04, RA (10 μM), or a combination of RA and DM04. Viability was assessed 48 hours later using acridine orange/propidium iodide (AO/PI) staining.

**Figure 2.**
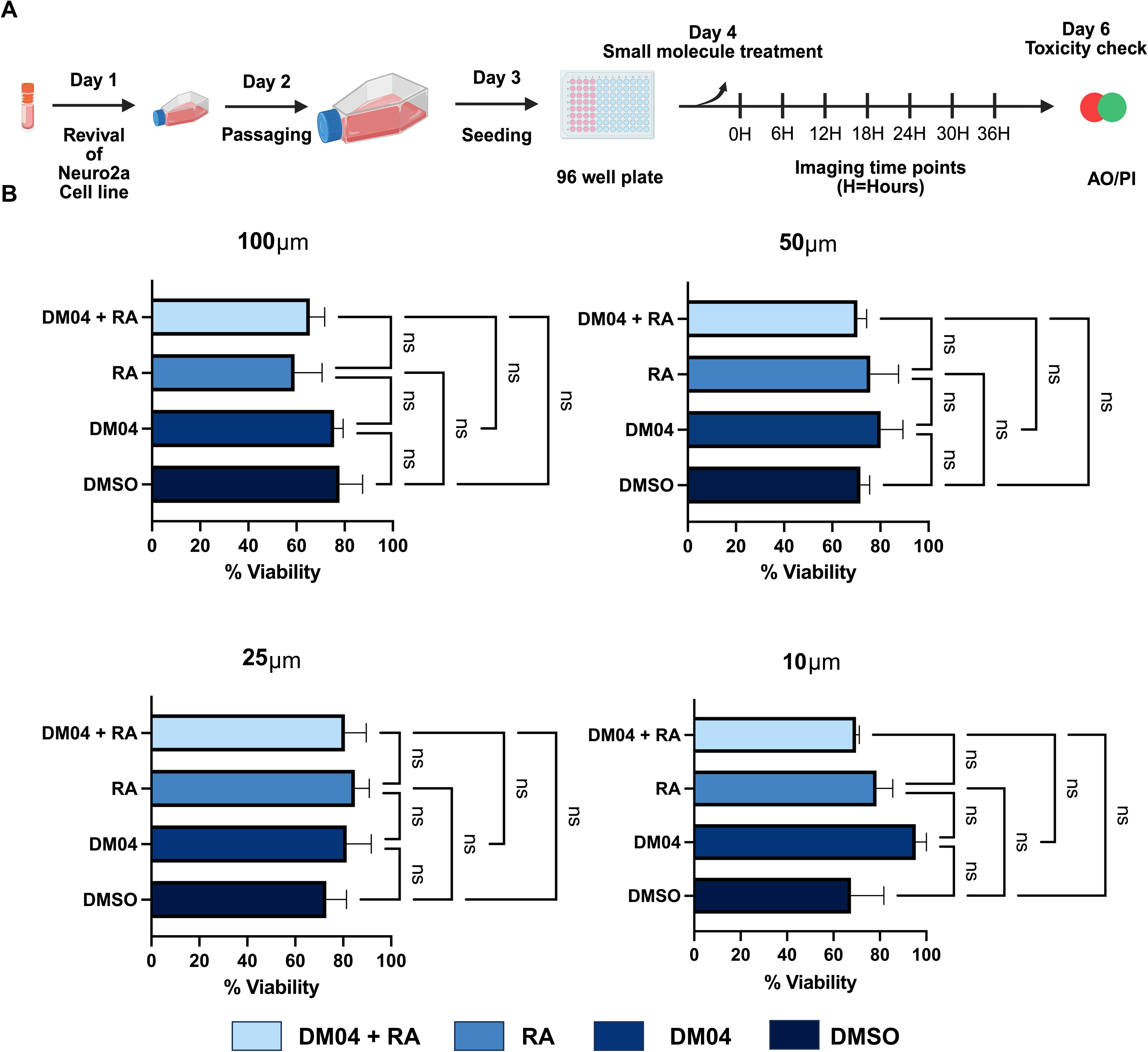
DM04 treatment across a concentration gradient shows no significant toxicity in Neuro2a cells. **(A)** Schematic of the experimental workflow : Neuro2a cells were revived and expanded over three days, followed by seeding in 96-well plates. On Day 4, cells were treated with DMSO, DM04, Retinoic Acid or a combination of DM04 and Retinoic Acid. Imaging was performed at regular intervals post-treatment, and cell viability was assessed on Day 6 using AO/PI staining. **(B)** Quantification of percentage viability of cells under DMSO, DM04, RA, DM04+RA conditions at different concentrations (100μM, 50μM, 25μM, 10μM). Approximately 60,000 cells were seeded onto each 96 well plate. The toxicity analysis was performed using data collected from three independent biological replicates and every replicate had three technical replicates to arrive at the mean percentage. Statistical analysis was performed using one-way ANOVA followed by Tukey’s multiple comparison test. ns: not significant.

The cells had no signs of cytoplasmic swelling, membrane blebbing, or nuclear fragmentation across all treatment groups, even at the highest DM04 concentration tested (100 μM). Cell morphology appeared comparable to DMSO treated controls, with preserved confluence and uniform soma size. No morphological perturbations were observed in the combination group (RA + DM04), suggesting that co-treatment does not induce additive stress.

AO/PI staining confirmed these observations at the molecular level, with uniformly green fluorescence in the majority of cells indicative of intact membranes and live-cell status. Quantitative analysis of viability (Figure 2B) across concentrations showed no significant differences between groups. Viability consistently remained above 80%, with no trend toward dose-dependent toxicity.

Together, these data establish that DM04 is well tolerated by neuronal cells across a tenfold concentration range and does not elicit measurable cytotoxicity or structural stress. The absence of adverse morphological or viability effects supports its suitability for use in neuronal repair paradigms, including combinatorial treatments with retinoic acid.

### DM04 promotes neurite outgrowth, with maximal extension observed at 10 μM

We next investigated whether DM04 influences neurite outgrowth in neuronal cells under differentiation-promoting conditions. The experimental setup is outlined in Figure 3A: Neuro2a cells were expanded over three days, treated on Day 4 with DMSO, RA, DM04, or a combination of RA and DM04, and then cultured in serum-deprived media to promote neurite initiation and elongation. Neurite morphology was evaluated 48 hours post-treatment by immunofluorescence staining of βIII-tubulin.

**Figure 3.**
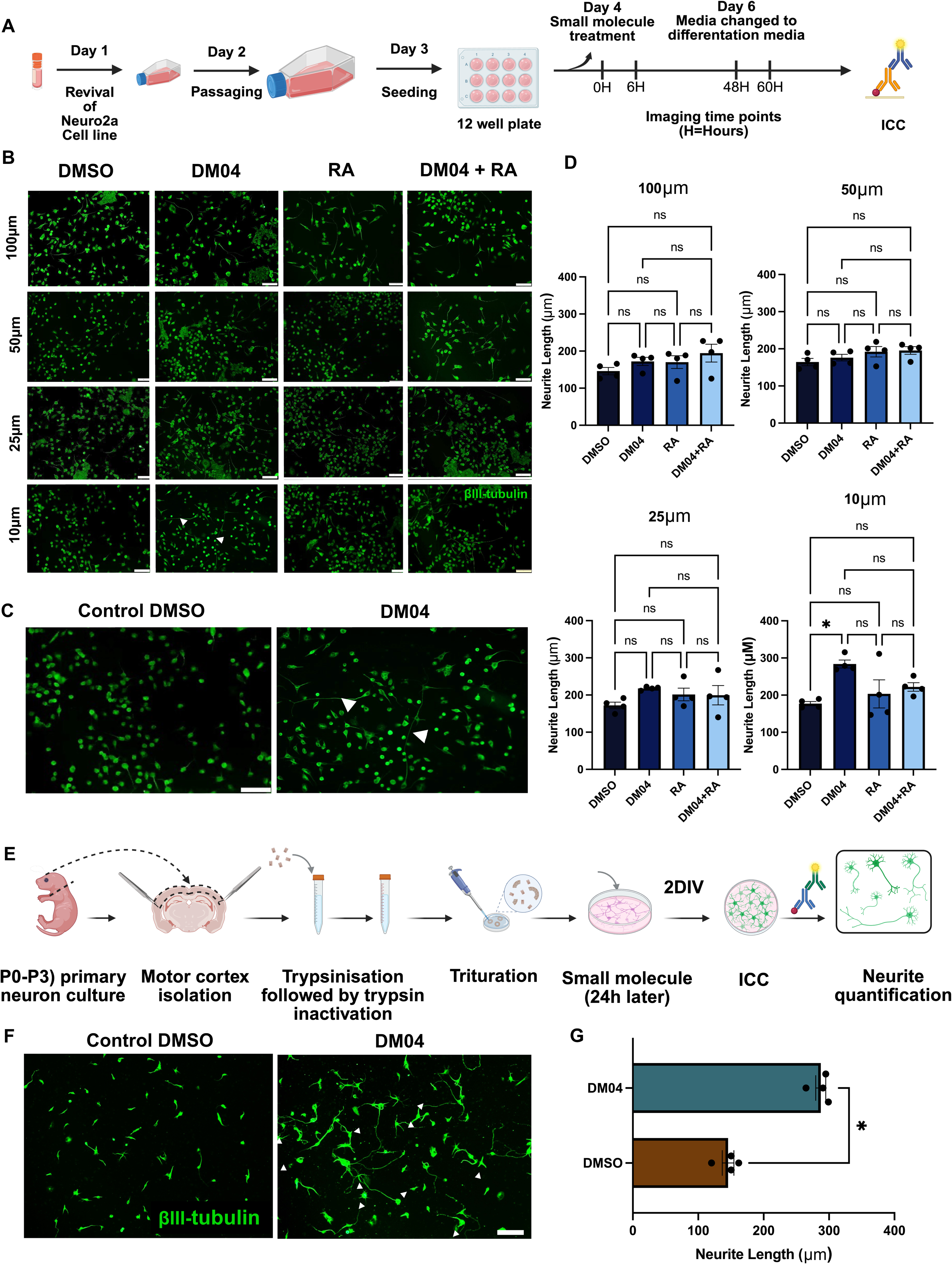
DM04 at 10 μM promotes the maximum neurite length in Neuro2a cells and primary neurons. Experimental workflow: Neuro2a cells were revived and expanded over three days, followed by seeding in 96-well plates. On Day 4, cells were treated with DMSO, DM04, RA, or a combination of DM04 and RA. The media was changed to serum-deprived media to allow cells to extend their neurites. (B) Representative fluorescent images of βIII-tubulin staining showing the neurite extension across a range of concentrations (100 μM to 10 μM) are shown. (C) Magnified image of DMSO and DM04-treated cells (Scale 50μm) at 10μM. (D) Quantification showing average neurite length in different concentrations. Approximately 60,000 cells were seeded onto each well of a 12-well plate. Neurite outgrowth was quantified from four (n=4) independent biological replicates. Statistical analysis was performed using Kruskal-Wallis test followed by Dunn’s multiple-comparisons test. At 10 μM, the overall treatment effect was statistically significant (Kruskal-Wallis, p = 0.0208), with DM04 showing a significant increase compared with DMSO (Dunn’s adjusted p = 0.0360). No significant differences were observed between DM04 and RA or between DM04 and DM04 + RA at 10 μM. No significant overall treatment effects were observed at 25 μM (p = 0.1003), 50 μM (p = 0.0990), or 100 μM (p = 0.2056).(E) Experimental workflow: Primary neurons from P0-P3 pups were isolated and cultured for 24 hours. Cells received DMSO and DM04 at 5μM concentration on Day 2. Post 48h treatment, cells were fixed and stained to detect βIII-tubulin. (F) Representative images (scale 50 μm) show enhanced neurite length indicated by white arrows. (G) Quantification showing comparison of neurite length between DMSO and DM04 treated primary neurons. Data are presented from four biological replicates per condition (n = 4). Neurite length was compared using a two-tailed Mann–Whitney U test (p = 0.0286)

Fluorescence microscopy revealed clear differences in neurite outgrowth across treatment conditions and concentrations (Figure 3B). At 100 μM and 50 μM, both RA and DM04 induced moderate neurite extension relative to DMSO, while the combination treatment produced a modest additive effect. At 25 μM, the response was similar. However, at 10 μM, DM04 alone induced the most pronounced neurite elongation, with several cells exhibiting long, unbranched processes (arrowheads, Figure 3B - bottom panel indicated by arrows). This enhancement was not further improved by RA co-treatment, suggesting that DM04 alone may be sufficient to trigger a maximal outgrowth response at this concentration.

A higher magnification comparison of DMSO and DM04 treated cells at 10 μM (Figure 3C) confirmed increased neurite number and length in the DM04 group, with greater uniformity across the field. Quantitative analysis of average neurite length (Figure 3D) showed a statistically significant increase in the DM04 treated group at 10 μM compared to DMSO (Dunn’s multiple-comparisons test, adjusted p = 0.0360). The overall difference between treatment groups at 10 μM was also statistically significant (Kruskal-Wallis test, p = 0.0208). DM04 did not show a statistically significant difference compared to RA or DM04 + RA at this concentration, indicating that RA co-treatment did not provide an additional significant increase in neurite outgrowth.

At 25 μM, 50 μM, and 100 μM, the overall differences between treatment groups were not statistically significant (Kruskal-Wallis test, p = 0.1003, p = 0.0990, and p = 0.2056, respectively). Similarly, the difference between DMSO and DM04 was not statistically significant at 25 μM (adjusted p = 0.0856), 50 μM (adjusted p > 0.9999), or 100 μM (adjusted p > 0.9999).

These results suggest that DM04 promotes neurite outgrowth in Neuro2a cells in a concentration dependent manner, with a clear optimum at 10 μM. Importantly, this effect is robust even in the absence of RA, indicating that DM04 can act as a functional retinoid analog with sufficient potency to drive neurite elongation on its own. This non-monotonic relationship with the strongest effect at the lowest concentration tested occurred despite comparable viability across all doses (Figure 2B), indicating that the diminished response at 25–100 µM reflects reduced efficacy rather than toxicity. Whether DM04 acts through any of these specific mechanisms remains untested in the present study. One plausible candidate is metabolic feedback, since our RNA-seq data show DM04 itself upregulates the retinoid-catabolizing enzyme Cyp26a1 (Figure 4), which could blunt effective ligand concentration at higher doses.

**Figure 4:**
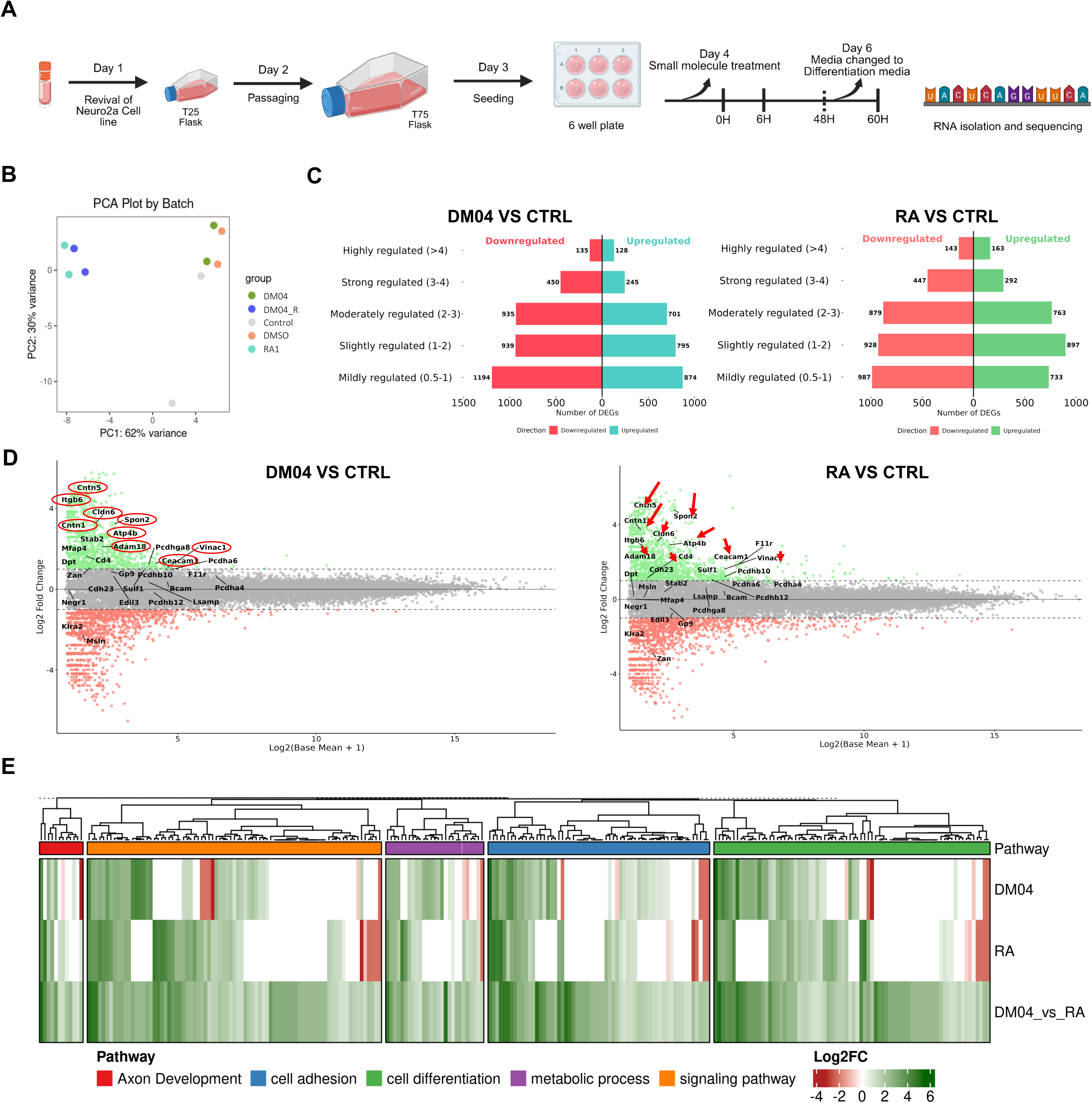
RNA-sequencing analysis of gene expression induced by DM04 and Retinoic Acid (RA) treatments. (A) Experimental Schematic. Workflow detailing the RNA-seq experimental design: Cells were seeded and cultured for 12 hours, followed by treatment with compounds (DM04 or RA), and then switched to differentiation media for 50 hours before cell harvesting, RNA isolation, and sequencing. (B) Principal Component Analysis (PCA) plot demonstrating the clear separation and clustering of experimental groups based on their RNA-seq data (C) Bi-directional bar plots showing the total number of up- and downregulated genes in DM04 vs. Control (Left) and RA vs. Control (Right). The genes are classified into five groups based on the absolute log_2_ fold change (L2FC) magnitude: Mildly (0.5 to 1), Slightly (1 to 2), Moderately (2 to 3), Strong (3 to 4), and Highly (>4) regulated. (D) MA plot comparing gene expression levels between the DM04 and Control conditions (left) MA plot comparing gene expression levels between the RA and Control conditions (right). In both MA plots, genes highlighted in red indicate significantly downregulated genes, while those in green represent significantly upregulated genes (Adjusted P-value < 0.05). (E) Heatmap visualization of pathway analysis showing the expression patterns of genes across key biological processes, including axon development, cell adhesion, cell differentiation, metabolic processes, and signaling pathways. Expression values are displayed as a color gradient of log_2_ fold change from low (red) to high (green) across the four comparison groups (DM04 vs Control, RA vs Control, DM04 vs RA, and RA vs DM04).

We extended compound testing in primary cortical neurons. Initial testing at 10 μM, the concentration optimized in Neuro2a cells, resulted in reduced neuronal viability and did not yield significant increase in neurite length, indicating toxicity in primary cultures at this dose. Based on this observation, subsequent experiments were performed using reduced concentration of DM04 (5 μM).

Fluorescence imaging revealed clear enhancement of neurite extension in DM04 treated cultures compared to controls (Figure 3F). DMSO treated neurons displayed short, sparsely branched processes typical of early-stage cultures, whereas DM04 treated neurons exhibited longer, more elaborate neurites with frequent branching points (arrowheads, Figure 3F, right). The overall neuronal density appeared similar between conditions, indicating that the observed changes were not due to altered survival or plating efficiency.

Quantification of average neurite length (Supplementary Table S5) confirmed a significant increase in DM04-treated neurons relative to DMSO (Figure 3G). Mean neurite length nearly doubled under DM04 treatment, suggesting that the RA-like signaling effects observed in Neuro2a cells may similarly operate in primary neurons to support cytoskeletal extension and maturation. Together, these results indicate that DM04 promotes robust neurite outgrowth in developing cortical neurons, reinforcing the notion that small molecules can enhance intrinsic growth capacity in postnatal neuronal populations.

A relevant consideration for the Neuro2a data is that neurite extension in this mitotic neuroblastoma line can accompany cell cycle exit, raising the possibility that DM04 acts by promoting differentiation linked proliferation arrest rather than a growth program. Several features of our dataset argue against this as the primary explanation. First, viability and confluence were comparable across DMSO, RA, DM04, and combination treatments at all concentrations tested (Figure 2B), with no morphological evidence of reduced cell density in DM04 treated cultures. Second, and most directly, primary postnatal cortical neurons (Figure 3E–G) are post-mitotic. As DM04 reproduced its neurite-outgrowth effect in this post-mitotic system, where proliferation arrest cannot contribute to the phenotype, the effect observed in Neuro2a cells is unlikely to be solely a consequence of altered proliferation.

We have compared tribenzamide-based DM04 promoted neurite outgrowth in Neuro2a cells with less hydrophobic two dibenzamide-based compounds DM01 and DM02 (Supplementary Figure S8). We also compared it with the less polar DM03 which is an esterified version of DM04. All three compounds have failed to increase neurite length in the study. This indicates the importance of the amphiphilic nature of DM04 which contains hydrophobic tribenzamide core and a terminal acidic polar functional group.

### DM04 recapitulates Retinoic Acid driven transcriptional programs, including core neuronal differentiation and ECM remodeling genes

To understand the molecular basis by which DM04 enhances neurite outgrowth and to assess whether it engages canonical Retinoic Acid (RA) signaling pathways, we performed bulk RNA-sequencing in Neuro2a cells treated with either DM04 or RA and compared their transcriptomic profiles to DMSO controls.

Principal component analysis (Figure 4B) demonstrated clear separation of treatment groups, confirming strong transcriptional divergence. Both DM04 and RA treatments led to extensive differential gene expression relative to control (Figure 4C), with a comparable number of genes classified as highly, strongly, or moderately regulated. Notably, MA plots (Figure 4D) revealed substantial overlap in the top differentially expressed genes between DM04 and RA, with shared genes highlighted using red arrows and circles. This overlap underscores that DM04 activates a transcriptional program strikingly similar to RA. Notably, this convergence occurs despite DM04’s failure to activate a RARE-luciferase reporter, suggesting that DM04 reaches RA-responsive gene targets through a route that does not depend on canonical RARE-mediated transactivation or convergence through an independent pathway.

Among the common DEGs were established components of the RA signaling pathway and neurodevelopmental regulators. These included RARβ (RA Receptor β), Crabp1 (an intracellular RA-binding protein), and RA-metabolizing enzymes like Cyp26a1 and Dhrs3, as well as key neuronal transcription factors such as Meis1, Mnx1, Zic1, Gbx2, and Fezf2 ^15^. Structural cytoskeletal genes like Tubb3 and Tubb2b markers of neuronal identity and maturation were also upregulated in both conditions ^40^. The detection of RARβ as a differentially expressed gene in this dataset also confirms that Neuro2a cells express and transcriptionally engage RARβ at baseline (see Discussion).

Beyond canonical RA targets, we observed a strong enrichment for genes involved in extracellular matrix (ECM) organization and adhesion, which are critical for creating a growth-permissive environment. Among these were Itgb6 (integrin β6)^41^, Spon2 (an ECM ligand known to promote neurite extension)^42^, Dpt (Dermatopontin), Mfap4 (a microfibril-associated ECM scaffold protein)^43^, Stab2 (a receptor involved in hyaluronan turnover)^44^, Edil3 (a matricellular protein affecting adhesion via integrin engagement), and Bcam (a cell adhesion molecule)^45^. These ECM-associated genes likely contribute to neurite elongation by remodeling the extracellular milieu and facilitating substrate interactions.

Together, these results demonstrate substantial overlap between DM04 and RA-associated transcriptional programs, without establishing that the two compounds act through the same receptor-level mechanism.

### DM04 activates canonical RA pathways and uniquely engages extracellular remodeling programs

To dissect the biological processes regulated by DM04, we performed pathway enrichment analysis on the set of genes significantly upregulated following treatment, and compared them to those induced by RA. As shown in Figure 5 (top panel), both RA and DM04 robustly activated overlapping pathways involved in neuronal differentiation, axon guidance, and retinoic acid signaling, including hallmark categories such as axonogenesis, response to retinoic acid, and neuronal cell body morphogenesis. This convergence is consistent with our earlier transcriptomic analysis and supports the notion that DM04 acts as an activator of RA-dependent gene programs.

**Figure 5:**
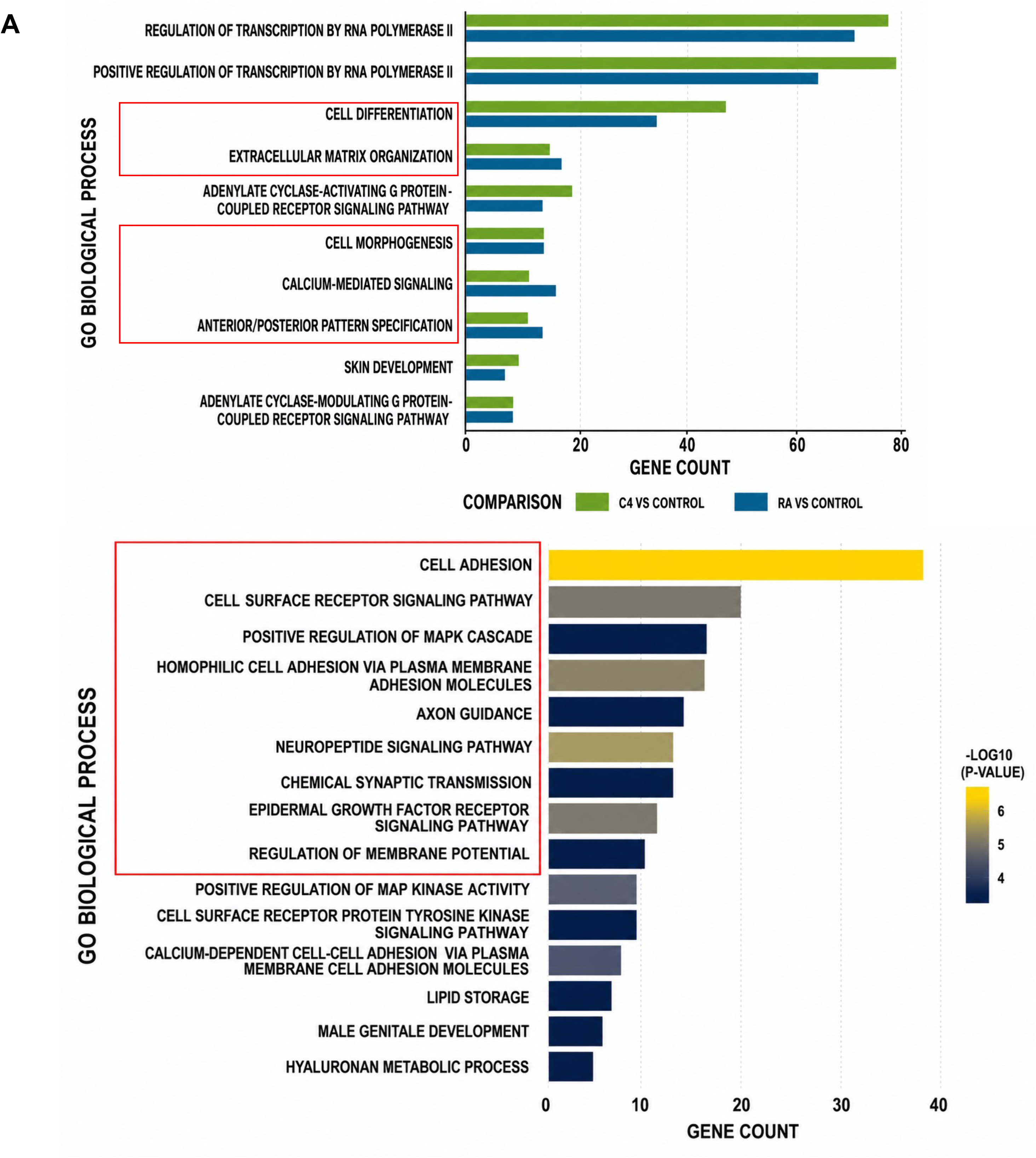
Shared and unique pathway enrichments in RA and DM04 treatment conditions. **(A)** Bar plot depicting the top enriched biological pathways based on upregulated genes in Retinoic Acid (RA) and DM04 conditions. **(Top panel)**: Pathways that are significantly enriched in both RA and DM04 treatments, highlighting strong overlap in gene programs related to neuronal differentiation, RA signaling, and axon development. **(Bottom panel)**: Pathways selectively enriched in DM04-treated cells, suggesting additional gene programs not triggered by RA alone. These include extracellular matrix remodeling, integrin engagement, and cytoskeletal dynamics, features likely contributing to the enhanced neurite outgrowth observed with DM04. Color gradient indicates adjusted p-values for enrichment significance (−log_10_ P value).

Interestingly, beyond this shared core, DM04 selectively upregulated a distinct set of gene categories not triggered by RA alone (Figure 5, bottom panel). These included pathways related to extracellular matrix (ECM) organization, cell–substrate adhesion, and integrin-mediated signaling features that may underlie the enhanced neurite outgrowth phenotype observed in primary neurons treated with DM04. The enriched categories were populated by genes such as Itgb6, Spon2, Dermatopontin (Dpt), Mfap4, Stab2, Edil3, and Bcam, all of which are associated with ECM remodeling, substrate interactions, or integrin signaling (see Supplementary Table S6). Many of these genes are known to regulate cytoskeletal dynamics, matrix viscoelasticity, or neurite anchorage, suggesting that DM04 may potentiate outgrowth through both transcriptional and microenvironmental modulation.

Together, these results indicate that while DM04 strongly overlaps with RA in activating neurodevelopmental programs, it additionally engages non-canonical, ECM-related modules that may provide a permissive substrate for process extension and stabilization.

### DM04 enhances hindlimb recovery following thoracic spinal cord injury

To evaluate whether DM04 improves motor recovery after injury, we used a thoracic (T7–T10) spinal crush model in adult mice followed by localized implantation of a gel foam soaked with either DM04 or DMSO at the injury site. Animals were maintained under standard post-operative care and allowed to recover. Animals were subjected to weekly behavioral assessments up to 42 days post-injury using the horizontal ladder rung assay, which provides sensitive measures of hindlimb placement, toe clearance, and limb elevation (Figure 6A).

**Figure 6:**
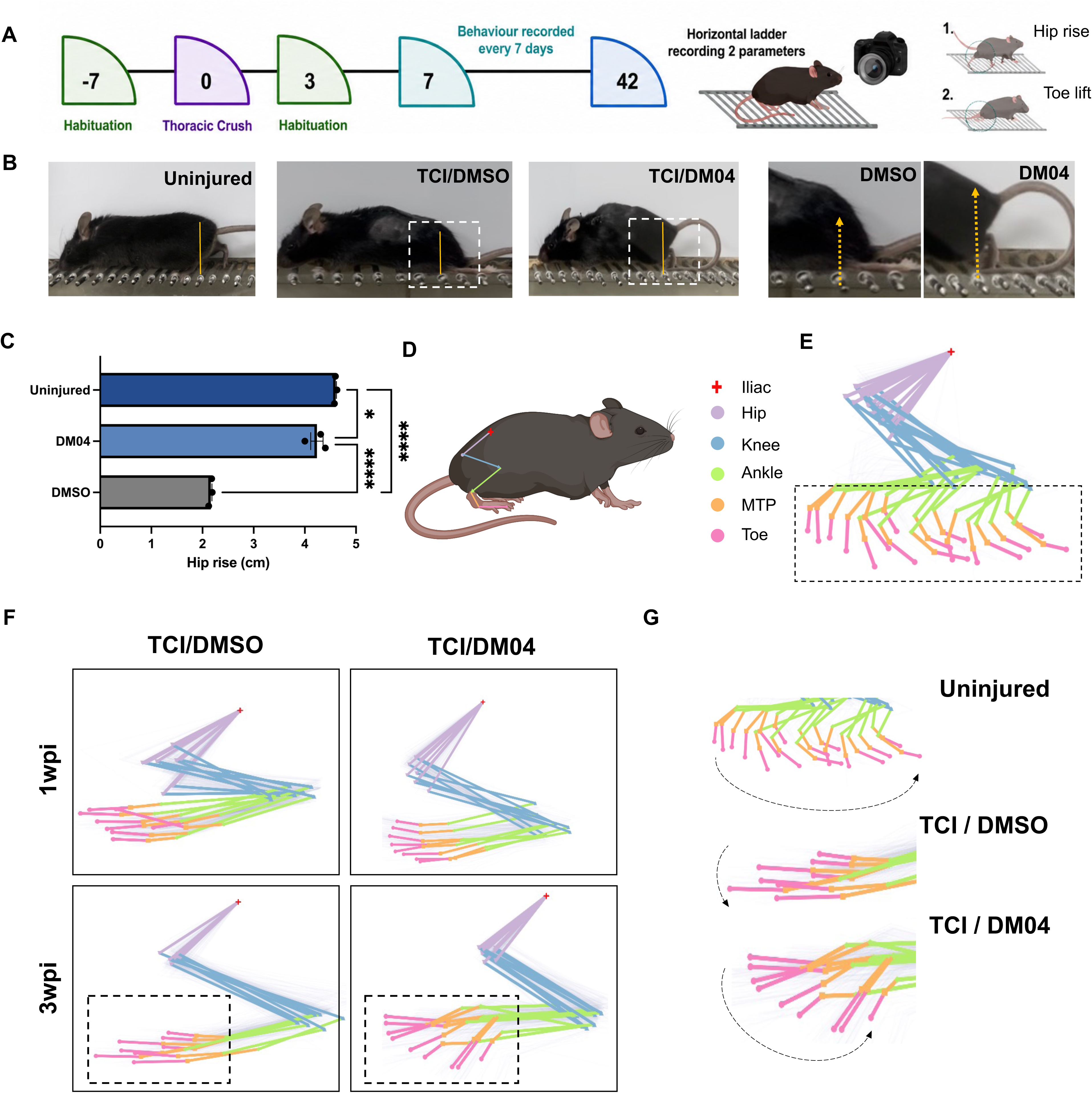
DM04 treated animals show hip rise and toe movement post 3 weeks of implantation. (A) Experimental timeline: The mice underwent 7 days of habituation, thoracic crush and DM04/DMSO implantation on day 0, and behavioral assessments every 7 days up to day 42 using a horizontal ladder test to measure hip rise and toe drag. (B) Representative images of uninjured, injured+DMSO, and injured+DM04 mice during ladder walking. Arrows indicate hip elevation.(C) Hip elevation quantification shows DM04-treated mice display significantly improved hip rise compared with DMSO-treated injured mice (mean ± SEM). (D) Diagram of tracked hindlimb joints during gait analysis, marking iliac crest, hip, knee, ankle, metatarsophalangeal joint (MTP), and toe using DeepLab Cut. (E) Representative hindlimb trajectory plots for uninjured mice during locomotion (F) Representative hindlimb trajectory of injured + DMSO and injured + DM04 treated animals at 0wpi and 3wpi where DM04 treated mice show improved gait patterns. (G) Overlaid joint trajectories for uninjured, injured+DMSO, and injured+DM04 mice demonstrate restoration of near-normal limb kinematics following DM04 treatment. Quantification of hip rise was performed on n = 3 animals per group (DMSO-treated injured, DM04-treated injured, and uninjured controls). For each animal, two frames per video were measured and averaged to yield a single value per animal. Frames were not treated as independent statistical units. Data are presented as mean ± SEM, with each data point representing one animal (n = 3/group). Groups were compared by one-way ANOVA (F(2,6) = 338.0, p < 0.0001, R² = 0.9912). Tukey’s multiple comparisons test: DMSO vs. DM04, p < 0.0001; DMSO vs. Uninjured, p < 0.0001; DM04 vs. Uninjured, p = 0.026.

During ladder walking, DM04-treated mice showed visibly improved hindlimb positioning compared to the DMSO treated mice. While DMSO treated injured mice frequently dragged the hindlimb across the rungs, DM04 treated animals demonstrated greater hip rise and more consistent foot elevation (Figure 6B). Quantification of hip rise (Supplementary Table S7) confirmed this observation, with DM04 treated injured mice exhibiting substantially higher elevation values than DMSO treated mice and approaching those of uninjured controls (Figure 6C), indicating improved weight bearing and hindlimb activation. DM04 treated injured mice exhibited significantly higher elevation values than DMSO treated mice (Tukey-adjusted p < 0.0001) and values approaching, though still modestly below, those of uninjured controls (p = 0.026).

To more precisely examine gait coordination, we performed kinematic analysis using five anatomical joint landmarks: iliac crest, hip, knee, ankle, metatarsophalangeal joint, and toe (Figure 6D)^37^. Joint trajectory plots showed that at 7 days post injury, both groups displayed disrupted and abbreviated limb movements. However, by 21 days, DM04 treated animals exhibited improvement in trajectory length, smoothness, and directional consistency (Figure 6F). In contrast, DMSO treated injured mice continued to show irregular, truncated paths with minimal recovery over the same period (Supplementary Video 3). When trajectories were overlaid with those of uninjured controls, the DM04 treated group demonstrated a closer approximation to normal gait dynamics, particularly in the amplitude and coordination of distal joint movement (Figure 6G, Supplementary Video 4).

These results indicate that a single localized application of DM04 at the time of spinal cord injury augments hindlimb motor coordination and step mechanics, suggesting that DM04 promotes functional recovery *in vivo*. The functional recovery aligns with the transcriptional and cellular effects observed in vitro, supporting a model in which DM04 enhances structural and behavioral repair following injury.

### DM04 is associated with axonal extension into the glial scar and increased PSD95-positive puncta following spinal cord injury

To determine whether DM04 treatment influences axonal responses following spinal cord injury, corticospinal tract axons were labeled by anterograde delivery of AAV-CAG-GFP into the layer 5 motor cortex prior to thoracic crush injury (Fig. 7A). The animals were injured 7 days post injection and DM04 was implanted using gel foam. Six weeks after injury, sagittal spinal cord sections were examined by immunofluorescence. In DMSO treated animals, GFP-labeled axons largely terminated at the lesion border, with almost no extension beyond the GFAP-positive glial scar. Representative sections from DM04 treated animals showed GFP-positive axonal profiles extending toward and within the glial scar, whereas GFP-positive axons in DMSO treated animals were predominantly observed near the lesion border. (Fig. 7B). To quantitatively assess GFP-labeled axonal signal beyond the glial scar, GFP fluorescence intensity was measured (Supplementary Table S8) in a region extending from the visually defined beginning of the GFAP-positive glial scar. DM04-treated animals exhibited significantly higher GFP fluorescence intensity compared with DMSO-treated animals (*P* = 0.0286, two-tailed Mann–Whitney U test; Fig. 7C).

**Figure 7:**
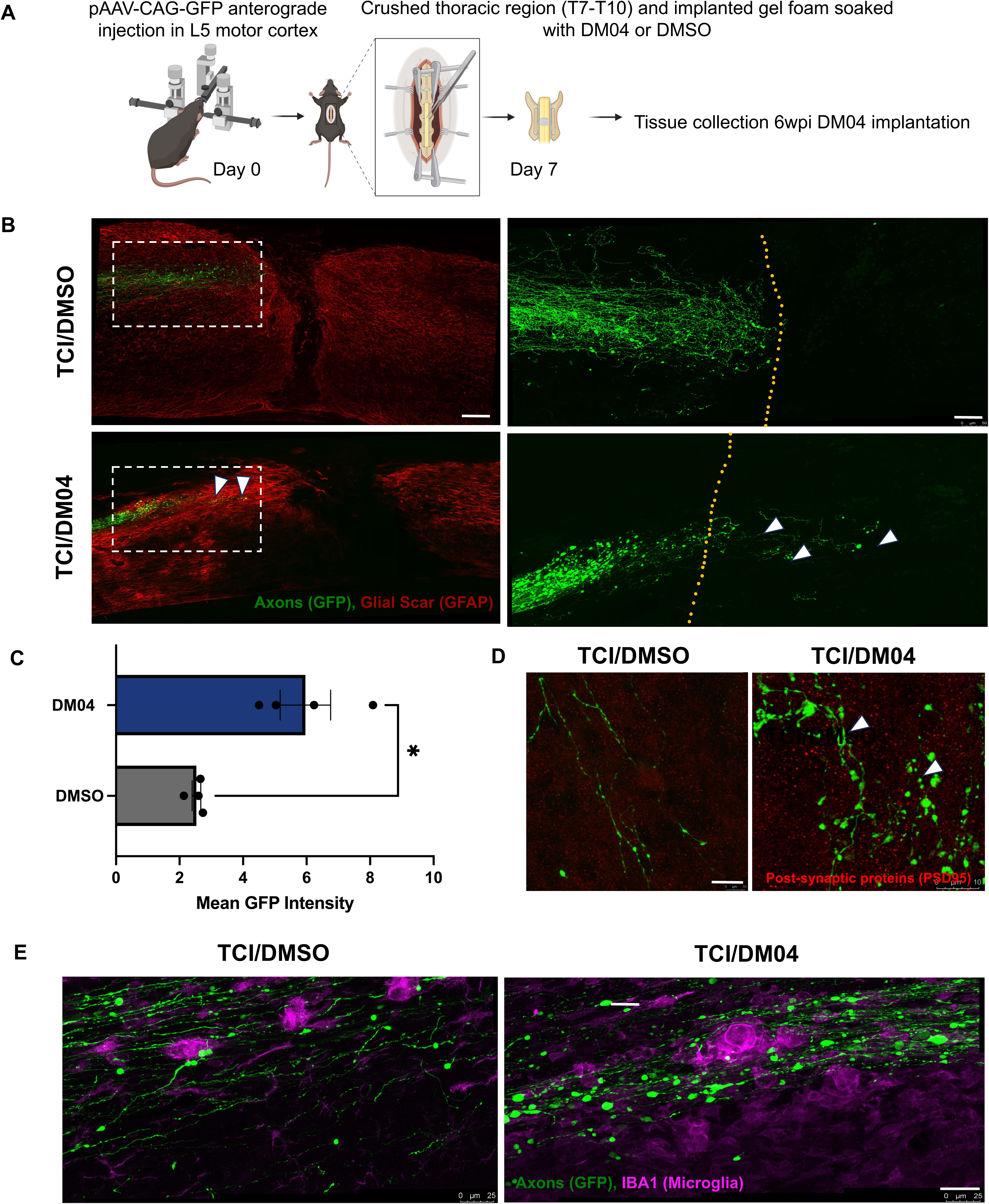
DM04 treated animals show axon extension into the glial scar and synapse formation. (A) Experimental timeline : Mice received anterograde cortical injections of pAAV-CAG-GFP into the L5 motor cortex followed by thoracic spinal cord crush injury (T7–T10) after 7 days. Gel foam soaked with either DM04 or DMSO was implanted at the lesion site, and tissues were collected 6 weeks post-injury for histological analysis. (B) Representative immunofluorescence images showing GFP-labeled corticospinal axons (green) attempting to extend (white arrows) into the GFAP-positive glial scar (red) at the lesion site ( Scale - Right : 100μm, left : 50μm). The dashed yellow line indicates the approximate lesion boundary. No quantitative analysis of axonal penetration was performed. (C) Quantification of GFP fluorescence intensity in the region extending beyond the visually defined beginning of the GFAP-positive glial scar. Each dot represents one animal. DM04 treatment significantly increased GFP fluorescence intensity compared with DMSO treatment (P = 0.0286, two-tailed Mann–Whitney U test; n = 4 animals per group) (D) Representative images of PSD95 immunostaining (red), a postsynaptic marker, in the vicinity of GFP-positive axons (green) where DM04 samples show PSD95-positive puncta (white arrows) associated with GFP-labeled axons, suggesting enhanced postsynaptic specialization and potential synapse formation. These observations are qualitative and were not quantified (Scale - 10μm). (E) Representative images of GFP-positive axons (green) and IBA1 positive microglia (magenta) within the lesion area, illustrating the cellular environment surrounding extending axons following DMSO or DM04 treatment (Scale - 25μm). No qualitative differences in microglial distribution or GFAP+ scar morphology were observed between treatment groups. For the qualitative assessments, 4 animals were used for the study in both groups.

To assess whether axonal growth was accompanied by evidence of synaptic specialization, sections were immunostained for the postsynaptic marker PSD95. Representative images from DM04 treated animals showed increased PSD95-positive puncta in close proximity to GFP-positive axons compared with DMSO treated controls, consistent with the possibility of enhanced postsynaptic organization at sites of axonal contact (Fig. 7D).

Finally, immunostaining for IBA1 revealed the distribution of microglia surrounding GFP-positive axons within the lesion area. GFP-labeled axons were observed traversing regions containing IBA1-positive cells in both treatment groups, with DM04-treated animals displaying qualitatively greater axonal presence within the injured tissue (Fig. 7E). Qualitative comparison of GFAP and Iba1 immunoreactivity between DMSO and DM04 treated animals did not reveal overt differences in glial scar morphology or microglial distribution at the lesion site (Fig. 7B, E), suggesting that DM04 treatment did not grossly alter the reactive glial or innate immune response to injury within the timeframe examined. Together, these qualitative findings indicate that DM04 treatment is associated with increased axonal extension toward the injury site and evidence of postsynaptic specialization near GFP-labeled axons.

### DM04 induces neural lineage commitment in human iPSCs

Neural induction from pluripotent stem cells is a widely used model to test the neurogenic potential of small molecules and signaling cues. Retinoic acid (RA) is a key developmental morphogen that promotes neural fate specification and is routinely used to induce neural progenitor cells (NPCs) from human induced pluripotent stem cells (hiPSCs)^16,46^. Given that our transcriptomic data revealed upregulation of canonical RA signaling genes such as RARβ, Crabp1, and Sox11 following DM04 treatment (see Figure 4), we hypothesized that DM04 may act as an RA pathway agonist and might therefore enhance neural induction even in vitro.

To directly test this hypothesis, we implemented a dual-SMAD inhibition-based neural induction protocol using hiPSCs treated with either RA, DM04, both in combination, or vehicle controls (Figure 8A). Cells were maintained for 10 days and then subjected to immunocytochemistry to detect Pax6, a hallmark transcription factor indicative of early neural fate specification. In the RA-only condition, robust Pax6 expression was observed with characteristic rosette-like NPC clusters (Figure 8C, second row, middle panel). Strikingly, DM04 treatment alone also resulted in a similarly high level of Pax6+ cells (Figure 8C, fourth row, middle panel), demonstrating that DM04 is independently capable of promoting neural differentiation.

**Figure 8.**
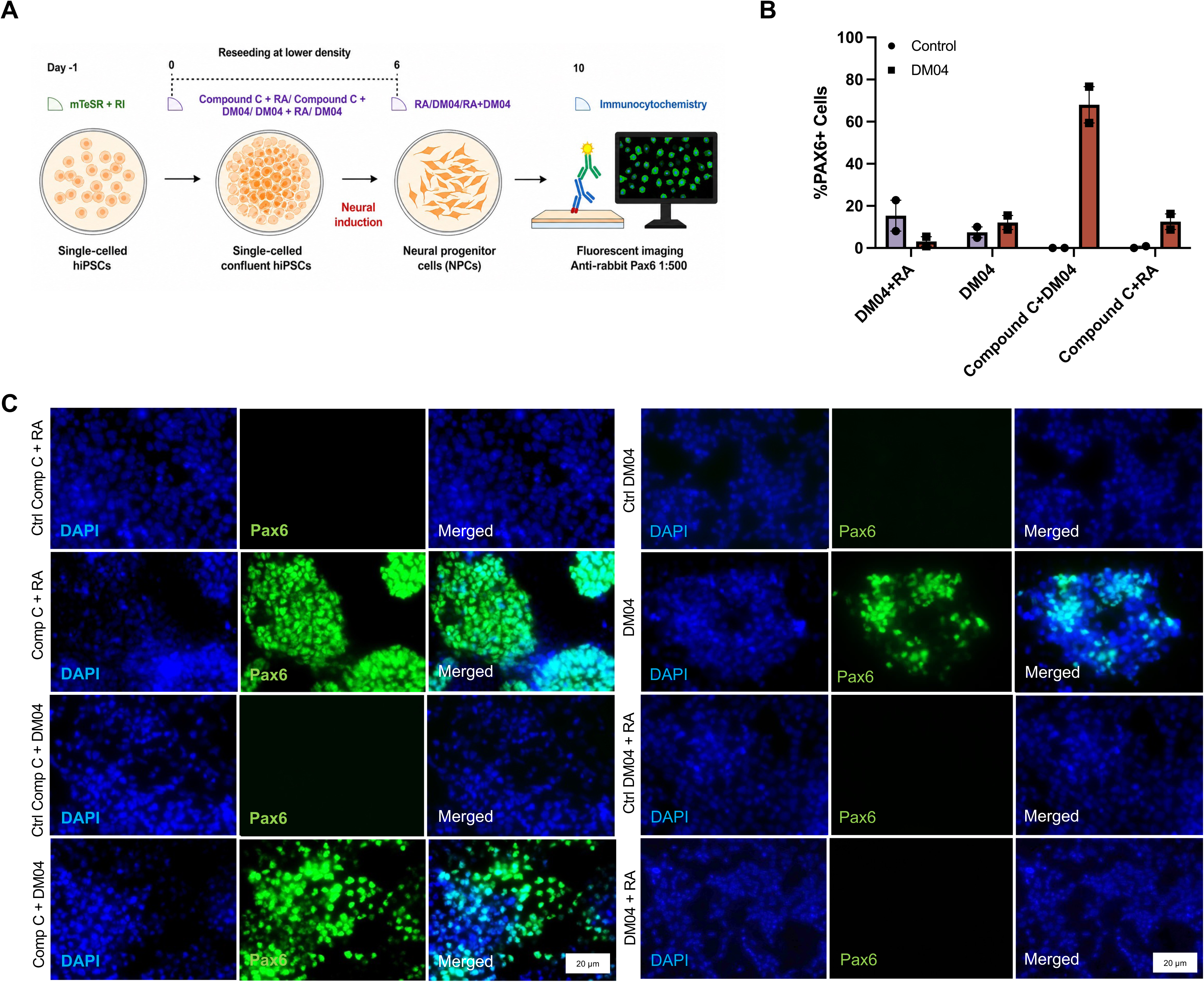
DM04 and RA promote neural induction in hiPSCs. (A) Schematic of the experimental workflow for neural differentiation. Single-cell hiPSCs were plated and treated with RA, DM04, or their combination. Cells were transitioned through a neural induction protocol to generate neural progenitor cells (NPCs), followed by immunocytochemistry on Day 10. (B) Quantification of the percentage of Pax6+ cells per field across treatment groups. Data are shown as individual values from n = 2 independent biological replicates with the mean + SEM (C) Fluorescence imaging of NPCs stained for Pax6 (green) and DAPI (blue) across different treatment groups. Pax6-positive cells (green) indicate successful neural induction. Both RA and DM04 treatments enhanced neural induction as seen via Pax6 expression, whereas the combinatorial treatment failed to induce neurogenesis. (Scale - 20 μm)

Together, these findings establish that DM04 can effectively recapitulate RA-induced neural induction in hiPSCs, inducing Pax6+ neural progenitors *in vitro*. The ability of DM04 to promote neurogenesis in a human stem cell model complements our *in vivo* results and positions DM04 as a promising neuroactive small molecule for both regenerative medicine and developmental modeling.

The effects were consistent across replicate experiments and comparable in intensity and distribution to the RA-only condition, supporting our hypothesis that DM04 resembles RA activity. However, the combination of RA and DM04 did not produce additive effects. Instead, this group showed reduced Pax6 staining relative to single-agent treatments (Figure 8C, bottom row, right panel), suggesting potential pathway interference, receptor saturation, or feedback inhibition when both agents are co-applied. Negative controls treated with vehicle (Figure 8C, top row) or non-neurogenic combinations (e.g., Ctrl Compound C + DM04; Figure 8C, third row) showed minimal Pax6 staining, confirming that the observed effects were specific to RA and DM04 action.

## Discussion

In this study, we report a novel small molecule, DM04, that enhances axon extension and functional recovery following spinal cord injury. Through a combination of transcriptomic profiling, behavioral assays, and cellular models, we demonstrate that DM04 promotes increased growth and activation of pro-growth transcriptional programs in neurons, promotes neurogenesis and facilitates improved locomotor coordination following spinal injuries *in vivo*. Pathway enrichment analysis revealed strong overlap between DM04 and known retinoic acid (RA)–regulated genes, and direct testing in human iPSCs confirmed that DM04 can drive neural lineage commitment, comparable to RA. Collectively, our data position DM04 as a potent neuroactive compound capable of influencing both regenerative potential through axonal extension and neural induction.

The rationale for pursuing DM04 was shaped by both the well-established neurogenic and regenerative roles of retinoic acid (RA) and the practical limitations of RA itself as a pharmacological agent. RA is a key developmental morphogen that regulates neuronal differentiation, axon guidance, and plasticity, and multiple studies have shown that RA signaling can re-engage growth-associated transcriptional programs in the injured CNS^12,14,15,38,46,47^. However, RA is chemically unstable, rapidly oxidized and metabolized in biological environments, and exhibits narrow dose tolerability, which has constrained its therapeutic use^17^. In our transcriptomic experiments, DM04 treatment induced a gene expression signature that closely overlapped with RA-regulated pathways, including upregulation of several classical RA-responsive genes. This raised the hypothesis that DM04 may function as a chemically more degradation-resistant RA-mimetic, though we have not directly compared its metabolic stability or receptor-complex behaviour with those of RA. Motivated by this, we evaluated DM04 in both spinal cord injury and human iPSC neural induction assays. In both contexts, DM04 reproduced key phenotypic outcomes associated with RA treatment, including enhanced axon outgrowth and Pax6-positive neural lineage commitment. While these results indicate that DM04 engages RA-like transcriptional programs, the precise receptor-level mechanism, whether via RARβ or alternative retinoid pathway components remains to be determined.

From a translational standpoint, the identification of a small molecule such as DM04 offers potential advantages over viral-vector–based gene therapies. AAV or lentivirus-mediated gene delivery faces challenges including host immune responses, packaging limitations, and restrictions on cell-type targeting^48^. In principle, small molecules can be synthesized at scale and delivered without genetic manipulation, realizing this potential for DM04 specifically will depend on the dose optimization, pharmacokinetic characterization, and mechanistic validation outlined above. The present findings in both injury repair and stem-cell differentiation models motivate this further preclinical characterization. This broader potential is supported by recent literature on small molecule discovery in regenerative medicine^49–53^.

A notable and somewhat unexpected finding was that DM04 did not activate a RARE-luciferase reporter, despite favorable docking predictions and despite closely recapitulating RA’s transcriptional and phenotypic effects. This dissociation suggests that DM04 does not act as a conventional RAR agonist driving canonical RARE-dependent transactivation unlike known agonists.^59,60^ The RARE-luciferase assay did not include a constitutive Renilla luciferase transfection control. The luciferase activity was therefore normalized to vehicle-treated controls at each time point rather than to an internal transfection control. This limitation should be considered when interpreting the magnitude of reporter activity, although the absence of DM04 induced reporter activation was consistent across the independent experiments.

Several non-mutually-exclusive explanations merit future investigation. DM04 may engage RARβ in a manner that alters coregulator recruitment or chromatin interactions without activating a minimal RARE reporter construct. It may act through non-genomic or rapid RAR signaling not captured by a transcriptional reporter. It may indirectly elevate endogenous RA levels or signaling tone via modulation of RA-metabolizing enzymes or it may act through a distinct nuclear receptor or signaling node that converges on an overlapping set of downstream targets. Resolving this will require complementary approaches to test whether DM04’s phenotypic effects are RARβ-dependent at all.

Importantly, the present study does not establish that the biological effects of DM04 are RARβ-dependent. The overlap with RA-responsive transcriptional programs and the computational prediction of RARβ engagement provide a rationale for this hypothesis, but receptor dependence will require direct perturbation or biochemical target-engagement studies.

A related open question is which RAR isotype(s) DM04 might engage. Our RNA-sequencing data show expression of RARβ transcripts in Neuro2a cells, but RARα and RARγ have not been profiled in our hands. Isotype-resolved profiling (qPCR or Western blot for RARα, RARβ, RARγ), together with coactivator recruitment assays and broader nuclear receptor profiling (e.g. RXRs, PPARs), will be needed to establish receptor selectivity and formally exclude off-target engagement.

Looking ahead, several promising avenues emerge from this work. While our findings suggest that DM04 activates retinoid-responsive pathways and enhances axon extension, the precise molecular target of this compound remains to be determined. Future studies will aim to clarify the molecular basis of DM04’s RA-like activity in the absence of RARE transactivation. Direct biochemical binding assays and profiling of RA-metabolizing enzyme activity will help determine whether DM04 acts through RARβ non-canonically, through upstream retinoid metabolism, or through a distinct target altogether. In parallel, it will be important to characterize the pharmacokinetic properties of DM04, including its metabolic stability, biodistribution, and ability to cross the blood–brain barrier, as well as its safety profile in long-term administration.

A related limitation is that native RA itself was not tested *in vivo* in this study. Our rationale for pursuing DM04 as an alternative to RA rests on the extensively documented instability and short *in vivo* half-life of RA and its rapid autoinductive metabolism, rather than on a direct *in vivo* comparison. Because RA is also susceptible to photoisomerization and oxidative degradation under the same conditions used for local, sustained-release delivery in this study, a locally implanted native-RA formulation would be expected to lose most of its activity before it could support the multi-day signaling window relevant to axon extension, making a like-for-like *in vivo* comparison technically difficult to interpret. A formal head-to-head comparison of RA and DM04 delivered via the same local route remains an important experiment for directly establishing DM04’s translational advantage over RA.

Notably, DM04 was not uniformly well tolerated across cell types at 10 µM — the concentration that produced maximal neurite outgrowth in Neuro2a cells — DM04 reduced viability in primary cortical neurons, requiring a lower working concentration (5 µM) in that system (Figure 3E–G). This cell type dependent tolerability underscores that the present toxicity assessment relied on a single AO/PI viability readout in one immortalised cell line, which can miss toxicity manifesting as metabolic stress, mitochondrial dysfunction, or membrane damage without outright cell death. A fuller safety profile, including orthogonal *in vitro* assays such as LDH release and mitochondrial membrane potential (e.g. JC-1/TMRM) in both Neuro2a and primary neuronal cultures, alongside standard *in vivo* haematological and histopathological screening, will be required before DM04 can be considered for further pharmacological development.

We also considered whether DM04’s effect on neurite outgrowth in Neuro2a cells could reflect an anti-proliferative or cell cycle arrest mechanism rather than a specific growth promoting program, given that Neuro2a is a mitotic line in which differentiation and cell cycle exit are often coupled. Viability and confluence data, together with the uniform application of the differentiation protocol across treatment groups, argue against this as the dominant explanation. The reproduction of the neurite-outgrowth phenotype in post-mitotic primary cortical neurons (Figure 3E–G), where cell-cycle-linked confounds do not apply, provides the strongest evidence that DM04 promotes a genuine growth-associated program rather than acting solely through proliferation blockade. A dedicated cell-cycle assay (e.g., Ki67 or EdU incorporation) would provide more direct confirmation and represents a logical next step. Neuro2a was used for RNA-sequencing because of its scalability, homogeneity, and cost effectiveness, with the key neurite outgrowth finding then validated in post-mitotic primary cortical neurons and in a fully human hiPSC neural induction system. We note as a limitation that our transcriptomic dataset (Figures 4–5), which underlies the mechanistic core of this study, was generated exclusively in Neuro2a cells and has not yet been extended to primary neurons or hiPSC-derived neurons. We restricted transcriptomic profiling to this homogeneous neuronal population deliberately: bulk RNA-sequencing of the lesion site *in vivo*, which comprises a heterogeneous mixture of neurons, glia, infiltrating immune cells and fibrotic cells, would confound cell-intrinsic, DM04-driven transcriptional changes in neurons with injury-associated and non-neuronal responses. Cell-type-resolved profiling of the lesion site, for example by single-nucleus RNA-sequencing, will be a priority for future studies.

The observation that DM04 does not produce additive effects when combined with RA suggests potential saturation of the RA signaling pathway, or the existence of homeostatic feedback mechanisms that dampen further response. These findings motivate additional investigations into the dynamics of receptor occupancy, downstream signaling crosstalk, and target gene regulation. Our findings provide early evidence that DM04 warrants further preclinical characterization, including dose optimization, formal pharmacokinetics, and mechanistic validation, before its translational potential can be more fully assessed.

Our use of the term ‘axon extension’ rather than ‘regeneration’ reflects the fact that the present study demonstrates enhanced axonal growth without direct evidence of long-distance regrowth across the lesion site or reconnection with distal targets. This distinction is consistent with a broader pattern in CNS injury research, where interventions that enhance intrinsic neuronal growth capacity, such as DM04, are often necessary but not sufficient to achieve functional circuit repair, particularly when a lesion cavity or dense glial/fibrotic scar presents a physical and biochemical barrier to sustained axon growth. Combining DM04 with strategies that address these extrinsic barriers, for example, biomaterial scaffolds to bridge the lesion cavity and provide a growth-permissive substrate, or co-treatment with agents that neutralize inhibitory extracellular matrix components, may be necessary to convert local axon extension into longer-range regenerative growth and, ultimately, functional recovery.

Future work will also explore the therapeutic potential of DM04 in more complex or chronic models of CNS injury, including spinal cord contusion and cortical stroke, as well as its impact on recovery in aged animals. Finally, there is significant opportunity in testing combinatorial strategies where DM04 is paired with pro-regenerative interventions^35^ such as biomaterial scaffolds^54^, electrical or optogenetic stimulation^55^, or epigenetic reprogramming^36^. Together, these future directions will help determine the full therapeutic potential of DM04 in neural repair.

## Materials and Methods

### Sequence Retrieval, Template Selection, and Homology Modeling

The primary amino acid sequence of mouse retinoic acid receptor beta (Mus musculus RARβ) was retrieved from the UniProtKB database under accession identifier P22605^62^. To identify suitable structural templates for comparative modeling, the RARβ sequence was queried against the Research Collaboratory for Structural Bioinformatics (RCSB) Protein Data Bank (PDB) using the Basic Local Alignment Search Tool for proteins (BLASTp)^63^. Based on significant sequence identity, high query coverage (>95%), and high crystallographic resolution, two homologous structures were selected as dual templates: the human retinoic acid receptor beta ligand-binding domain (PDB ID: 1XDK, 1.60 Å resolution)^64^ and the retinoic acid receptor complex (PDB ID: 5UAN, 1.95 Å resolution)^65^.

Three-dimensional structural modeling was carried out using MODELLER version 10.8^66^ via an automodel comparative multi-template protocol. Atomic spatial coordinates of the native agonist, all-trans retinoic acid (Ret), were preserved and transferred from the template binding pocket into the target model during optimization. An ensemble of 100 candidate models was generated using standard conjugate gradient minimization followed by molecular dynamics with simulated annealing. Candidate structures were evaluated using the Discrete Optimized Protein Energy (DOPE) scoring function and GA341 assessment scores^67^. The top-ranking model with the lowest DOPE score was selected, and stereochemical quality was validated by Ramachandran plot analysis using PROCHECK and MolProbity ^68^, confirming >95% of residues within favored and allowed dihedral regions without stereochemical clashes.

### Molecular Docking (AutoDock Vina and CB-Dock)

Molecular docking of small molecules (DM01, DM02, DM03, DM04, and reference agonist all-trans retinoic acid / 9CR_RET) into the RARβ LBD was executed using the cavity-detection guided docking algorithm CB-Dock ^69^ coupled with AutoDock Vina ^70^. The modeled RARβ structure was prepared by removing crystallographic waters, adding polar hydrogens, and computing Kollman partial charges with AutoDockTools ^71^. Ligand geometries were energetically optimized, and Gasteiger charges and active torsional degrees of freedom were assigned.

CB-Dock performed curvature-based surface analysis to detect concave topological binding pockets on the receptor. The primary canonical ligand-binding cavity (Cavity 1, volume: 1595 Å³) was mapped with grid center coordinates of X = 14.991, Y = 31.297, Z = 103.532, and dimensions of 25 × 25 × 25 Å. Local conformational searches were conducted in AutoDock Vina with an exhaustiveness of 8. Binding affinities were scored in kcal/mol, and top-ranking poses were selected for subsequent molecular dynamics.

### Molecular Dynamics Simulations (GROMACS)

All-atom MD simulations of RARβ in its apo state and complexed with DM04 and all-trans retinoic acid were performed using GROMACS 2026.3 with native CUDA GPU acceleration^72^. The protein was parameterized with the AMBER ff14SB force field^73^. Small-molecule topologies were parameterized using the General AMBER Force Field (GAFF2) ^74^with semi-empirical AM1-BCC quantum partial charges derived via Antechamber (AmberTools) ^75^.

Each system was solvated in a rhombic dodecahedron box with a minimum solute-wall clearance of 1.0 nm using explicit TIP3P water molecules ^76^ and neutralized at 0.15 M NaCl. Energy minimization was conducted with steepest descent until maximum force converged below 1000 kJ/mol/nm. Positional restraints were applied during 100 ps of NVT equilibration at 300 K using the velocity-rescaling thermostat (τ_t = 0.1 ps) ^77^, followed by 100 ps of NPT equilibration at 300 K and 1.0 bar with the C-rescale barostat (τ_p = 2.0 ps) ^78^. Unrestrained 100 ns production runs were executed with a 2.0 fs timestep, LINCS constraints on hydrogen bonds ^79^, Particle Mesh Ewald (PME) electrostatics (1.0 nm cutoff) [80], and complete GPU offloading (-nb gpu -pme gpu -bonded gpu -update gpu). Trajectories were analyzed for backbone RMSD, C-α RMSF, radius of gyration (Rg), ligand RMSD, solvent accessible surface area (SASA), and internal hydrogen bonds.

### Synthesis of Methyl 3-nitrobenzoate (Compound 1)

A solution of 3-nitrobenzoic acid (5.9 mmol, 1 g, 1 eq.) in methanol (50 mL, anhydrous) was equilibrated at room temperature for 10 minutes and cooled down at 0 °C. Thionyl chloride (14 mmol, 1.77g, 2.5 eq.) was added dropwise to the solution at 0 °C. Then the reaction mixture was stirred at room temperature for 12 h. The disappearance of the starting material confirmed the completion of the reaction. The volatiles were removed on the rotavapor and the resulting mixture was partitioned between ethyl acetate and water. The organic layer was dried over sodium sulphate, filtered, and concentrated. Column chromatography (0 to 25% ethyl acetate in hexane, v/v) afforded the desired product as a white solid (1.06 g, 96%).

^1^H NMR (400 MHz, CDCl_3_) δ 8.87 (s, 1H), 8.42 (d, *J* = 8.1 Hz, 1H), 8.38 (d, *J* = 7.8 Hz, 1H), 7.67 (t, *J* = 8.0 Hz, 1H), 4.00 (s, 3H).

^13^C NMR (100 MHz, CDCl_3_) δ 164.9, 148.2, 135.2, 131.8, 129.6, 127.4, 124.6, 52.8.

### General method for the reduction of nitro group

To a solution of nitro aryl and arylamide compound (0.1 mmol) in ethyl acetate (EtOAc) (10 mL), Pd/C (10% by wt.) was added and the reaction was initiated under an atmosphere of H₂(g) with constant stirring. The progress of the reaction was monitored using thin layer chromatography (TLC). The disappearance of the starting material confirms the completion of the reaction. The reaction mixture was filtered, and the filtrate was dried over rotavapor to afford the desired reduced product as a solid, which is used in the next step without further characterization.

### General method for the amide coupling

To a mixture of aryl acid (1.2 mmol) and aryl amine (1 mmol) in dichloromethane (10 mL, anhydrous), 2-chloro-1-methylpyridinium iodide (1.5 mmol) and triethyl amine (3 mmol) were added. The reaction mixture was stirred at 60 °C under reflux for 6 h in an inert atmosphere. The disappearance of the starting material confirms the completion of the reaction. The volatiles were removed on the rotavapor. Column chromatography (20 to 60% ethyl acetate in hexane, v/v) afforded the desired product as a white solid.

### General method for the saponification of ester benzamides

To a solution of aryl amide derivative (1 mmol) in tetrahydrofuran (20 mL), 1M lithium hydroxide (20 mL) was added, and the reaction was stirred for 12 h at room temperature. The disappearance of the starting material confirms the completion of the reaction. The solution was then poured into water (20 mL). The pH of the reaction solution was adjusted to 4. The aqueous layer was extracted with EtOAc (2×30 mL). The organic layers were combined, dried over Na_2_SO_4_, and concentrated on rotavap to afford the desired product as a solid.

#### Methyl 3-benzamidobenzoate (Compound 2)

^1^H NMR (400 MHz, CDCl_3_) δ 8.16 – 8.14 (m, 1H), 8.06 (ddd, *J* = 8.1, 2.2, 1.0 Hz, 1H), 7.98 (s, 1H), 7.89 (dd, *J* = 8.3, 1.3 Hz, 1H), 7.84 – 7.81 (m, 1H), 7.56 (dt, *J* = 2.7, 1.9 Hz, 1H), 7.53 – 7.49 (m, 1H), 7.49 – 7.44 (m, 2H), 3.92 (s, 3H).

^13^C NMR (100 MHz, CDCl_3_) δ 166.7, 165.8, 138.1, 134.6, 132.1, 131.0, 129.3, 128.9, 127.0, 125.6, 124.7, 121.0, 52.3. HRMS-ESI (m/z): Calculated for C_15_H_14_NO_3_ (M+H)^+^: 256.0974, found 256.0973.

#### 3-benzamidobenzoic acid (DM01)

^1^H NMR (400 MHz, DMSO-*d_6_*) δ 12.95 (s, 1H), 10.42 (s, 1H), 8.04 (ddd, *J* = 8.1, 2.2, 1.1 Hz, 1H), 7.98 (dd, *J* = 8.2, 1.5 Hz, 2H), 7.71 – 7.65 (m, 1H), 7.62 – 7.45 (m, 4H).

^13^C NMR (100 MHz, DMSO-*d_6_*) δ 167.6, 166.1, 139.8, 135.1, 132.2, 131.6, 129.3, 128.9, 128.1, 124.9, 121.5. HRMS-ESI (m/z): Calculated for C_14_H_12_NO_3_ (M+H)^+^: 242.0817, found 242.0811.

#### Methyl 3-(3-nitrobenzamido) benzoate (Compound 3)

^1^H NMR (400 MHz, CDCl_3_) δ 10.79 (s, 1H), 8.84 (s, 1H), 8.50 – 8.38 (m, 3H), 8.11 (ddd, *J* = 8.1, 2.1, 1.0 Hz, 1H), 7.86 (t, *J* = 8.0 Hz, 1H), 7.77 – 7.71 (m, 1H), 7.55 (t, *J* = 7.9 Hz, 1H), 3.89 (s, 3H).

^13^C NMR (100 MHz, CDCl_3_) δ 166.5, 164.0, 148.2, 139.6, 136.3, 134.7, 130.7, 130.5, 129.7, 126.8, 125.4, 125.1, 122.9, 121.4, 52.7. HRMS-ESI (m/z): Calculated for C_15_H_13_N_2_O_5_ (M+H)^+^: 301.0824, found 301.0814.

#### 3-(3-nitrobenzamido) benzoic acid (DM02)

^1^H NMR (400 MHz, DMSO-*d_6_*) δ 13.02 (s, 1H), 10.76 (s, 1H), 8.84 (t, *J* = 1.9 Hz, 1H), 8.48 – 8.41 (m, 3H), 8.07 (ddd, *J* = 8.1, 2.1, 1.0 Hz, 1H), 7.86 (t, *J* = 8.0 Hz, 1H), 7.74 – 7.70 (m, 1H), 7.52 (t, *J* = 7.9 Hz, 1H).

^13^C NMR (100 MHz, DMSO-*d_6_*) δ 167.5, 163.9, 148.2, 136.4, 134.7, 131.7, 130.7, 129.5, 126.8, 125.3, 125.0, 122.9, 121.7. HRMS-ESI (m/z): Calculated for C_14_H_11_N_2_O_5_ (M+H)^+^: 287.0668, found 287.0662.

#### Methyl 3-(3-benzamidobenzamido) benzoate (DM03)

^1^H NMR (500 MHz, DMSO-*d_6_*) δ 10.49 (d, *J* = 19.6 Hz, 2H), 8.41 (dt, *J* = 54.8, 1.8 Hz, 2H), 8.08 (ddd, *J* = 8.1, 2.2, 1.0 Hz, 1H), 8.03 (ddd, *J* = 8.2, 2.1, 0.9 Hz, 1H), 8.01 – 7.98 (m, 2H), 7.75 – 7.69 (m, 2H), 7.64 – 7.60 (m, 1H), 7.58 – 7.50 (m,4H), 3.88 (s, 3H).

^13^C NMR (125 MHz, DMSO-*d_6_*) δ 166.6, 166.2, 166.1, 151.7, 139.8, 132.2, 129.6, 129.1, 128.9, 128.1, 125.2, 124.7, 124.0, 123.1, 121.3, 120.4, 52.7. HRMS-ESI (m/z): Calculated for C_22_H_19_N_2_O_4_ (M+H)^+^: 375.1345, found 375.1345.

#### 3-(3-benzamidobenzamido) benzoic acid (DM04)

^1^H NMR (400 MHz, DMSO-*d_6_*) δ 12.94 (s, 1H), 10.47 (s, 2H), 8.39 (dt, *J* = 32.3, 1.7 Hz, 2H), 8.07 – 7.98 (m, 4H), 7.70 (ddd, *J* = 8.3, 7.7, 4.6 Hz, 2H), 7.61 (dt, *J* = 2.7, 2.1 Hz, 1H), 7.58 – 7.47 (m, 4H).

^13^C NMR (100 MHz, DMSO-*d_6_*) δ 167.6, 166.2, 166.1, 139.9, 139.8, 132.2, 131.7, 129.3, 129.1, 128.9, 128.1, 124.9, 124.8, 124.0, 123.1, 121.6, 120.5

HRMS-ESI (m/z): Calculated for C_21_H_17_N_2_O_4_ (M+H)^+^: 361.1188, found 361.1181.

HPLC: DM04 purity > 97% (Supplementary Data 5,6)

### Neuro2a Cell culture

Mouse neuroblastoma (Neuro2a or N2a) cell line was maintained in Dulbecco’s Modified Eagle Medium (DMEM, Gibco #11885084) supplemented with 10% fetal bovine serum and 1% penicillin-streptomycin-gentamicin at 37°C in a humidified 5% CO₂ incubator. For experiment, cells were thawed from frozen stocks, expanded in T75 flasks and passaged at a 1:1 ratio using 0.1% trypsin-EDTA (Sigma, #T4799). For toxicity and viability assays, cells were seeded in 96 well plates at a density of 60,000 cells per well in 150 µL of complete medium. For neurite growth assays and Bulk-RNA sequencing, cells were seeded in 12 well plates and 6 well plates at densities of 60,000 cells/well (in 1mL medium) and 120,000 cells/well (in 2mL medium) respectively. Following a 12-hour adhesion period, cells were treated with the small molecules DM04, all-trans Retinoic Acid (RA) or a combination of both. Compounds were diluted in complete medium to final concentrations of 10, 25, 50 and 100 µM, with the final DMSO concentration not exceeding 0.1% (v/v). A vehicle control (0.1% DMSO) was included in all experiments.

### Toxicity and viability assessment

Cell morphology was monitored at every 6 hours for 36 hours post treatment using phase contrast microscopy. At the 48 hour endpoint, viability was quantified using acridine orange (AO) and propidium iodide (PI) staining. Briefly, the culture medium was aspirated and cells were washed with phosphate buffer saline (PBS, MP BIOMEDICALS #092810307) and detached using 0.1% trypsin-EDTA. The reaction was neutralised with complete medium and the cells were centrifuged at 1500 x g for 3 min. The pellet was resuspended in PBS and an equal volume of AO/PI staining solution (10 µg/mL AO, 5 µg/mL PI in PBS) was added and incubated in dark for 5 min. Viable (AO⁺/PI⁻) and non-viable (PI⁺) cells were quantified using an automated cell counter. Data are presented as the mean percentage of viable cells ± SEM from three independent experiments (Figure 2B, Supplementary Table S3).

### Immunocytochemistry and Quantification of Neurite length

To assess differentiation, Neuro2a cells were treated for 48 hours with DM04, RA, DM04+RA, or DMSO in complete medium. The medium was then replaced with a serum free medium with 1% penicillin-streptomycin-gentamycin for an additional 12 hours. Post incubation, the spent medium was aspirated out and cells were washed with PBS three times. Cells were fixed with 4% paraformaldehyde (PFA,Sigma, #30525-89-4) for 15 min at room temperature, permeabilised with 0.3% Triton X-100 and blocked with 1% bovine serum albumin (BSA, Sigma, #A7906). Cells were incubated overnight at 4°C with primary antibody against βIII-tubulin (Rabbit anti-βIII-tubulin, CST, #2128S, 1:500). After washing, cells were incubated for 2 hours at room-temperature with an Alexa Fluor 488-conjugated secondary antibody (Goat anti-rabbit IgG, Invitrogen #A-11008, 1:500). Fluorescent images were acquired using an Olympus microscope under consistent exposure settings. Neurites were manually traced and their lengths were measured using Olympus CellSen software. The experiment was repeated four times and approximately neurites from 200 cells from each batch and sample were traced. Statistical analysis was performed using GraphPad Prism (10.3.0.). Data from four independent experiments were analyzed by Kruskal–Wallis followed by Dunn’s test. Supplementary Table S4).

### Read Alignment and Differential Gene Expression Analysis

All surfaces and equipment were treated with RNaseAWAY (MoleculeBioProducts #7003) to prevent degradation. The spent media was aspirated out and 800µL of Trizol in ice cold condition was added to each well. A cell scraper was used to detach the cells and collected in 2ml MCTs before snap freezing them in liquid nitrogen. Total RNA was isolated from treated cells (DMSO, DM04, RA) in 6 well plates using a NucleoSpin RNA Mini kit for RNA purification (MacheryNagel #740955.50) according to the manufacturer’s instructions. RNA concentration and integrity were determined using a Qubit and only samples with an RNA Integrity Number (RIN) score between 9 and 10 were used for library preparation.

RNA libraries were sequenced on an Illumina NovaSeq 6000 platform generating 30 million paired-end reads per library. Quality control was performed with FastQC^29^, and adapter trimming and quality filtering were executed using fastp (v0.23)^28^. Clean reads were mapped to the mouse reference genome (GRCm38/mm10) using STAR (v2.7)^30^ in two-pass mode under default parameters. Gene-level read counts were determined using featureCounts (Subread package)^31^ against GENCODE/Ensembl gene models. Low-abundance genes (fewer than 10 counts across all libraries) were filtered out prior to statistical testing.

Differential expression analysis was conducted on raw integer count matrices using DESeq2 (v1.44)^32^. Library size factors were estimated via the median-of-ratios method, and dispersions were fitted using a Negative Binomial generalized linear model with empirical Bayes shrinkage. Multiple testing correction was performed using the Benjamini-Hochberg False Discovery Rate (FDR) procedure coupled with DESeq2’s automated independent filtering. Genes with an absolute log2 fold change ≥ 1.0 (|log₂FC| ≥ 1.0) and an FDR-adjusted p-value < 0.05 (padj < 0.05) were designated as differentially expressed genes (DEGs). To visualize sample clustering, raw counts were transformed using the Variance Stabilizing Transformation (VST), and Principal Component Analysis (PCA) was performed on the top 1,000 most variable genes.

Biological pathway and functional enrichment analyses of DEGs were performed using the Database for Annotation, Visualization, and Integrated Discovery (DAVID v2021)^33^ interfacing with the KEGG pathway database^34^ and Gene Ontology (Biological Process). Pathways exhibiting a fold enrichment > 1.5 and a Benjamini-Hochberg adjusted p-value < 0.05 (FDR < 0.05) were considered significantly altered.

### pAAV-CAG-GFP production

Recombinant adeno-associated virus serotype 9 (AAV9) particles were produced in HEK293T cells and purified using a commercial AAV purification kit (Takara, #6675). HEK293T cells were transfected with pAAV-CAG-GFP, pAAV2/9n, and pAdDeltaF6 plasmids (Source - Regalla laboratory, CSIR-CCMB) using polyethylenimine (PEI, 40 kDa; Polysciences, #24765). Sixty-Seventy hours after transfection, cells were harvested and AAV particles were isolated according to the manufacturer’s protocol. Capsid integrity was assessed by resolving purified viral particles on a 10% SDS-PAGE gel (35 mA, 120 min), followed by Coomassie Brilliant Blue staining and overnight destaining in a methanol-acetic acid solution. Viral genome titers were determined by quantitative PCR using TB Green (Takara, #RR82WR). Prior to qPCR, viral preparations were treated with DNase I (Thermo Fisher Scientific, #EN0521) at 37°C for 30 min to remove residual plasmid DNA, followed by four 10-fold serial dilutions. Viral genome copy numbers were calculated using a plasmid-derived standard curve. AAV preparation yielded titers of ≥ 1 × 10^11 viral genomes (vg)/µL, and intact capsid formation was confirmed by SDS-PAGE analysis.

### Animal Husbandry

All the animal procedures were approved by the Institutional Animal Ethics Committee (IAEC) at CSIR-CCMB. Animals were housed under a 12-hour day and night cycle. All experiments were performed on wild-type C57BL/6J mice of both sexes. All animal procedures were approved by the Animal Ethics Committee at CSIR CCMB and adhered to the IAEC guidelines.

### Anterograde injections, thoracic Crush and compound treatment

Animals were randomly allocated to the respective treatment groups prior to experimentation. Adult C57BL/6J mice of mixed sex (>12 weeks old, 23-26g) were used for the *in vivo* studies. Both male and female animals were included in the study; however, owing to the limited sample size, the study was not powered to evaluate sex-dependent effects. Mice were anesthetised with a mixture of Ketamine (50mg/mL) and Xylazine (100mg/mL). Anterograde injections of pAAV-CAG-GFP were performed using the stereotaxic setup. Four points : (M/L=+2, A/P=0), (M/L=+2, A/P=+1), (M/L=-2, A/P=0), and (M/L=-2, A/P=+1) were marked relative to the bregma (M/L=0, A/P=0). After drilling at these coordinates, the 0.4 µL of the AAV was injected per point to label the axons. Seven days post the anterograde injections, thoracic crush injury was performed by procedure as described previously^35,36^. For the thoracic crush injury, a midline incision exposed the spinal column, followed by blunt muscle dissection. Mice were secured in a custom spine stabilizer, and a laminectomy was performed at the T7-T10 vertebrae using spring scissors to expose the spinal cord. The spinal cord was then carefully crushed with equal force using blunt forceps (width-1mm). A gel foam was cut at a dimension of 1mm X 1mm X 1mm and soaked in 20µL of either DMSO(vehicle) or DM04 (10µM) and implanted on the injury site. The local implant therefore contained 0.2 nmol DM04 per 20 μL preparation (approximately 72 ng based on the molecular weight of DM04). This represents a localized exposure rather than a systemically defined pharmacological dose, and tissue exposure and pharmacokinetic parameters were not determined. The muscles cut were carefully put to cover the injury site and the incision made was sutured. Betadine was applied on top of the sutured area and 2 units of Meloxicam were administered. Animals were kept on a heating pad until ambulatory and monitored for 6 weeks post-surgery.

### Immunohistochemistry

Six weeks post-injury, animals were perfused in accordance with IAEC guidelines and tissue processed as described previously^35,36^. The vertebral column was dissected and washed in PBS. The entire cord was fixed overnight in 4% paraformaldehyde (PFA) with gentle rocking. The spinal cord was then dissected from the vertebral column, and the lesion area was embedded in 12% gelatin. Following embedding, the tissue was post-fixed once again in 4% PFA before sectioning. Sagittal sections were prepared using a vibratome and processed for immunohistochemistry. Sections were permeabilized and blocked in PBS containing 0.3% Triton X-100 and 5% fetal bovine serum for one hour at room temperature to minimize non-specific binding. Primary antibody against GFAP (Cell Signaling Technology, #3670; dilution 1:500)/ Iba1 (Cell Signaling Technology, #17198S; dilution 1:500)/PSD95 (Cell Signaling Technology, #3540; dilution 1:500) was diluted in a blocking solution and incubated with the sections overnight at 4°C with gentle rocking. After incubation, sections were washed three times in PBS and incubated with species appropriate fluorescent secondary antibodies for two hours at room temperature in the dark. Following secondary incubation, sections were rinsed thoroughly in PBS and mounted on glass slides using DAPI Fluoroshield (Sigma, #F6057), which served both as a nuclear stain and mounting medium. The lesion center was identified based on the peak of GFAP immunoreactivity. Sections (30 µm) spanning the lesion were imaged on Leica STED confocal microscope (63x objective and 100x objective). Histological characterization was performed with treatment identities masked during imaging and quantification.

### Machine Learning–Based Locomotion Analysis

Locomotor behavior was analyzed using DeepLabCut (DLC), an open-source deep learning framework for markerless motion tracking ^37^. A new DLC project was created and high-quality hindlimb locomotion videos were imported under consistent lighting conditions. Representative frames were extracted and manually labeled for key hindlimb landmarks: iliac crest, hip, knee, ankle, metatarsophalangeal joint (MTP), and toe. Labeled frames were reviewed to ensure consistency and then used to train a ResNet-50–based neural network for up to 50,000 iterations, with training parameters such as learning rate and batch size adjusted to optimize tracking accuracy. Model performance was evaluated using a validation dataset, and the final trained network was applied to the full set of locomotion videos to automatically extract joint trajectories. These trajectories were then used for quantitative gait analysis.

### Behaviour Assays

Prior to the experimental procedures, animals were habituated to a horizontal ladder in which they were trained to traverse a ladder with evenly spaced rungs to evaluate stepping accuracy and limb coordination to minimize stress and ensure consistent performance during locomotion trials as described previously^35^. Following surgical intervention, behavioral recordings were conducted on a weekly basis up to six weeks to assess locomotor recovery and coordination over time. High-resolution videos were recorded from a lateral view to capture detailed limb movements. Two primary gait parameters-hip rise and toe fall were analyzed to quantify vertical limb displacement and step precision. Behavioral assessments and subsequent video analyses were performed by investigators blinded to the treatment group. The hip rise videos and screenshots were renamed and given to the blinded observer to calculate hip rise.

### Maintenance of Human Induced Pluripotent Stem Cells (hiPSCs)

Episomal hiPSC lines (Gibco, Cat. #A18945) were routinely maintained under feeder-free conditions. Cells were cultured on Matrigel-coated plates (Corning, Cat. #354234) in mTeSR™1 medium (STEMCELL Technologies, Cat. #85850), which supports long-term self-renewal and maintenance of pluripotency. Cultures were passaged every 3–4 days at 60–70% confluency using Accutase (Thermo Fisher Scientific, Cat. #A1110501) to ensure optimal growth and viability.

### Neural Induction

Neural induction was performed with minor modifications to established protocols. On Day 0, hiPSCs were dissociated into single cells using Accutase (5–7 min, 37 °C) and seeded at high density (1 × 10⁶ cells/cm²) on Matrigel-coated plates in mTeSR™1 medium. Upon reaching confluency, cells were assigned to treatment groups: (1) Compound C + RA + SHH, (2) Compound C + DM04 + SHH, (3) DM04 only, and (4) DM04 + RA + SHH, with respective DMSO controls. On Day 1, neural induction medium (NIM) composed of DMEM/F12, N2 and B27 supplements, GlutaMAX™, and 0.2% heparin was added, along with Compound C (1 µM) and DM04 (10 µM). From Day 3 to 6, RA (100 nM) and DM04 (10 µM) were included. On Day 6, cells were gently dissociated and replated at lower density on Matrigel-coated coverslips. Compound C and DM04 were withdrawn, and cells were treated with RA and SHH (100 µg/mL) until Day 10, except for the DM04-only group. Cells were then fixed in 4% PFA and processed for immunocytochemistry. To assess successful neural induction, at Day 10 post induction, cells were incubated overnight at 4°C with primary antibody Pax6 (Rabbit anti-Pax6, CST, #42-6600, 1:500). After washes, cells were incubated for 2 hours at room-temperature with an Alexa Fluor 488-conjugated secondary antibody (Goat anti-rabbit IgG, Invitrogen #A-11008, 1:500). Fluorescent images were acquired using Zeiss Apotome microscope under consistent exposure settings.

### Primary Neuron Culture and Compound Treatment

Plates were coated with 25 μg/ml Poly-D-Lysine (Gibco, #A3890401) in 1X PBS and incubated at 37 °C for 2 hours, followed by three rinses with autoclaved Milli-Q water and air-drying under a laminar flow hood. C57BL/6J P0 pups were sacrificed in accordance with IAEC guidelines, and motor cortices were dissected using a brain matrix and transferred to an ice-cold dissection solution. The dissection solution was made from two components: Solution A (1X HBSS without Ca²⁺ and Mg²⁺) and Solution B (9.9 mM HEPES), mixed in a 5 ml:2.8 ml ratio and supplemented with 33.3 mM D-glucose and 43.8 mM sucrose in sterile Milli-Q water. Tissue was incubated in 0.1% trypsin (Sigma, #T4799) in 1X HBSS at 37 °C for 20 minutes, followed by quenching in 10% FBS (Gibco, #10438026) in dissection solution for 5 minutes at room temperature. The tissue was gently triturated 5–10 times in 1 ml of plating medium consisting of Neurobasal medium (Gibco, #21103049) with 1X B27 (Gibco, #17504044), 10% FBS, 1X Penicillin-Streptomycin, and gentamicin. After 24 hours, the medium was replaced with fresh Neurobasal containing 1X B27, 1X PSG, and 1 µM Ara-C (Sigma, #C1768). Neurons were then treated with small molecule DM04 (5 µM) or DMSO (control), and treatment efficacy was assessed after 48 hours via immunostaining for βIII-tubulin (CST, #2128S) (Figure 3F, Supplementary Table S5).

### Luciferase assay

Neuro2a (N2a) cells were cultured in Dulbecco’s Modified Eagle Medium (DMEM; Gibco) supplemented with 10% fetal bovine serum (FBS) and 1% penicillin-streptomycin at 37°C in a humidified incubator with 5% CO₂. Cells were first seeded in 12-well plates overnight prior to transfection. Cells were transiently transfected with the pGL3-RARE-Luciferase (Addgene plasmid # 13458)^61^ luciferase reporter plasmid using polyethyleneimine (PEI) at a 1:1 DNA:PEI (w/w) ratio. As a positive control for transfection, cells were also transfected with pAAV-CAG-GFP. Additionally, cells were co-transfected with pAAV-CAG-GFP and pGL3-RARE-Luciferase to ensure the latter does not affect the transfection and DMSO as a negative control. 24h post transfection, cells transfected with pGL3-RARE-Luciferase were washed and seeded in white bottom 96-well plates and were left to adhere to the surface overnight. They were then treated with vehicle (DMSO), DM04, or all-trans retinoic acid (RA) as a positive control. Luciferase activity was measured at 0, 12, 24, and 48 h following treatment. At each time point, luciferase activity was quantified using the Steady-Glo Luciferase Assay System (Promega, Cat. No. E2510) according to the manufacturer’s instructions. Briefly, an equal volume of Steady-Glo reagent was added directly to each well containing culture medium, and the plate was incubated for 5 min at room temperature to allow cell lysis and stabilization of the luminescent signal. Luminescence was measured using a microplate luminometer. Since a Renilla luciferase internal control was not included, firefly luciferase activity was normalized to the corresponding DMSO-treated control for each time point and expressed as fold change relative to the vehicle control. Data are presented as mean ± SEM from three independent biological replicates, each with three technical replicates. Statistical analysis was performed using two-way ANOVA followed by Šídák’s multiple-comparison test.

## Supporting information

All supplementary figures and tables

## Authors contributions

Ishwariya Venkatesh (I.V.) and Debabrata Maity (D.M.) conceived and designed the study. Katha Sanyal (K.S.), Dhanuush Balakannan (D.B.), Sk Rajibul Haque (S.R.H.), Rutuja Arun Pendharkar (R.A.P.), Prakash Chermakani (P.C.), Manojkumar Kumaran (M.K.), and Pratikhya Acharya (P.A.) performed the experiments, provided technical expertise, and contributed to data acquisition, analysis, and interpretation. Manuscript drafting was led by K.S., D.B., and I.V., with input from D.M. Figure preparation was carried out by K.S. and D.B. All authors reviewed, revised, and approved the final manuscript. *D.M. and I.V. jointly supervised the study and contributed equally as corresponding authors. ‡K.S., D.B., and S.R.H. contributed equally to this work.

## Acknowledgements

The authors gratefully acknowledge financial support from the Council of Scientific and Industrial Research (CSIR–HCP, FBR), CSIR–Centre for Cellular and Molecular Biology (CSIR-CCMB), the Science and Engineering Research Board (SERB, SRG), the Department of Biotechnology (DBT, Biomedical Sciences), the Anusandhan National Research Foundation (ANRF-ARG), and the BFI Biome programme, all awarded to I.V. This work was partially supported by the National Laboratories Scheme under the ULIP sub-scheme [MLP002604]. We sincerely thank the CCMB core facilities and their staff for their invaluable technical assistance throughout this study, particularly Zareena (Tissue Culture), Sairam and Prashanth (Animal House), Dr. Nitla Mahesh and Suman Bhandari (Microscopy), and the FBC facility. We also acknowledge the laboratories of Dr. Vinay Nandicoori, Dr. Santosh Chauhan, Dr. Sriram Varahan for providing access to resources and technical support. We are especially grateful to our lab manager, Dhruva Kesireddy, grants manager, Dr. Shringika Soni, and LabTech support, Mr. Venkateshwarulu, whose dedicated assistance and efficient management of laboratory operations were instrumental in the successful completion of this work.

## Supporting Information

The Supporting Information contains the following material: additional computational characterization of the RARβ–DM04 interaction (Supplementary Figures S1–S2), physicochemical stability and aggregation data for DM04 (Supplementary Figures S3–S6), RARE-luciferase reporter and structure-activity data (Supplementary Figure S7–S8), and supporting quantitative tables for cell viability, neurite outgrowth, luciferase activity, RNA-sequencing gene lists and hip rise measurements (Supplementary Tables S2–S7). The supplementary data also includes the behaviour videos of DMSO treated (Supplementary videos 1 and 3) and DM04 treated animals (Supplementary videos 2 and 4) where the first two videos were 1wpi while the others were 3wpi. The Supplementary Data 5 and 6 includes the HPLC purity results for DM04.

## Supplementary Figures

Figure S1. Additional computational docking analysis of the RARβ–DM04 complex, including detailed hydrogen-bonding and hydrophobic contact residues within the ligand-binding pocket (related to Figure 1).

Figure S2. Molecular dynamics trajectory analysis (backbone RMSD and Cα RMSF) of apo-RARβ, RARβ–RA, and RARβ–DM04 complexes over a 100 ns simulation.

Figure S3. HPLC stability profile of DM04 following UV irradiation (360 nm) for up to 60 min.

Figure S4. HPLC stability profile of DM04 following oxidative treatment with H₂O₂ (1 mM, 3 h).

Figure S5. Concentration-dependent UV–Vis absorbance spectra of DM04 (5–25 μM).

Figure S6. Dynamic light scattering (DLS) analysis of DM04 aggregation propensity (5–25 μM).

Figure S7. Time-course RARE-luciferase reporter activation by RA and DM04 over 48 hours.

Figure S8. Comparative neurite outgrowth analysis of DM01, DM02, and DM03 relative to DM04 in Neuro2a cells.

## Supplementary Table

Table S1. Percentage yield of compounds in the study.

Table S2. Quantitative RARE-luciferase reporter data corresponding to Figure S7.

Table S3. Cell viability (AO/PI) quantification data corresponding to Figure 2B.

Table S4. Statistical summary of Neuro2a neurite outgrowth analysis corresponding to Figure 3D.

Table S5. Quantification of neurite length in primary cortical neurons corresponding to Figure 3G.

Table S6. List of extracellular matrix- and integrin-associated genes uniquely enriched by DM04 treatment, corresponding to Figure 5.

Table S7. Hip Rise measurements related to Figure 6.

Table S8. Mean GFP Intensity of axons entering glial scar.

Table S9. Pax6 positive cell count percentages.

## Supplementary Data

Supplementary Video 1. 1wpi rung ladder video (DMSO)

Supplementary Video 2. 1wpi rung ladder video (DM04)

Supplementary Video 3. 3wpi rung ladder video (DMSO)

Supplementary Video 4. 3wpi rung ladder video (DM04)

Supplementary Data 5 and 6 : HPLC purity results for DM04.

## Notes

### Competing Interest Statement

The authors have declared no competing interest.

### Summary of Updates

Figure 3 - Neurite growth data replotted with improved statistics. Figure 7 - Regeneration quantified. Figure 8 - Quantification done. Supplementary Figures 1 to 6 - Molecular Docking Results and Stability Data included. Overall wordings updated, consistency checked. Claims validated.

