## Supplementary Data 5 for "A tribenzamide small molecule with retinoic acid-like activity promotes axon growth and motor recovery after spinal cord injury"

### [1]H NMR spectra of Compound-1 in $CDCl_3$

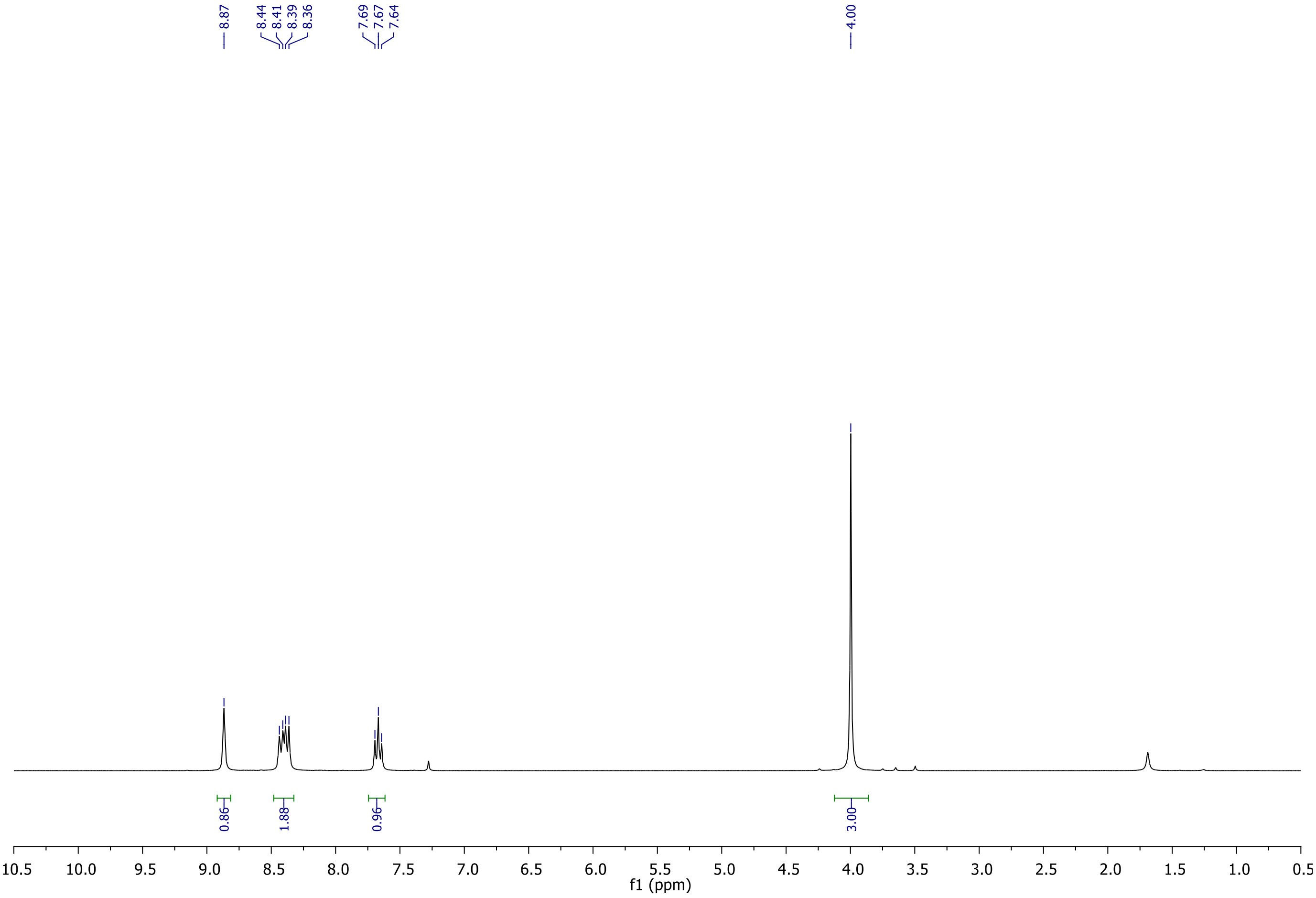

**$^{13}C$ NMR spectra of Compound-1 in $CDCl_3$**

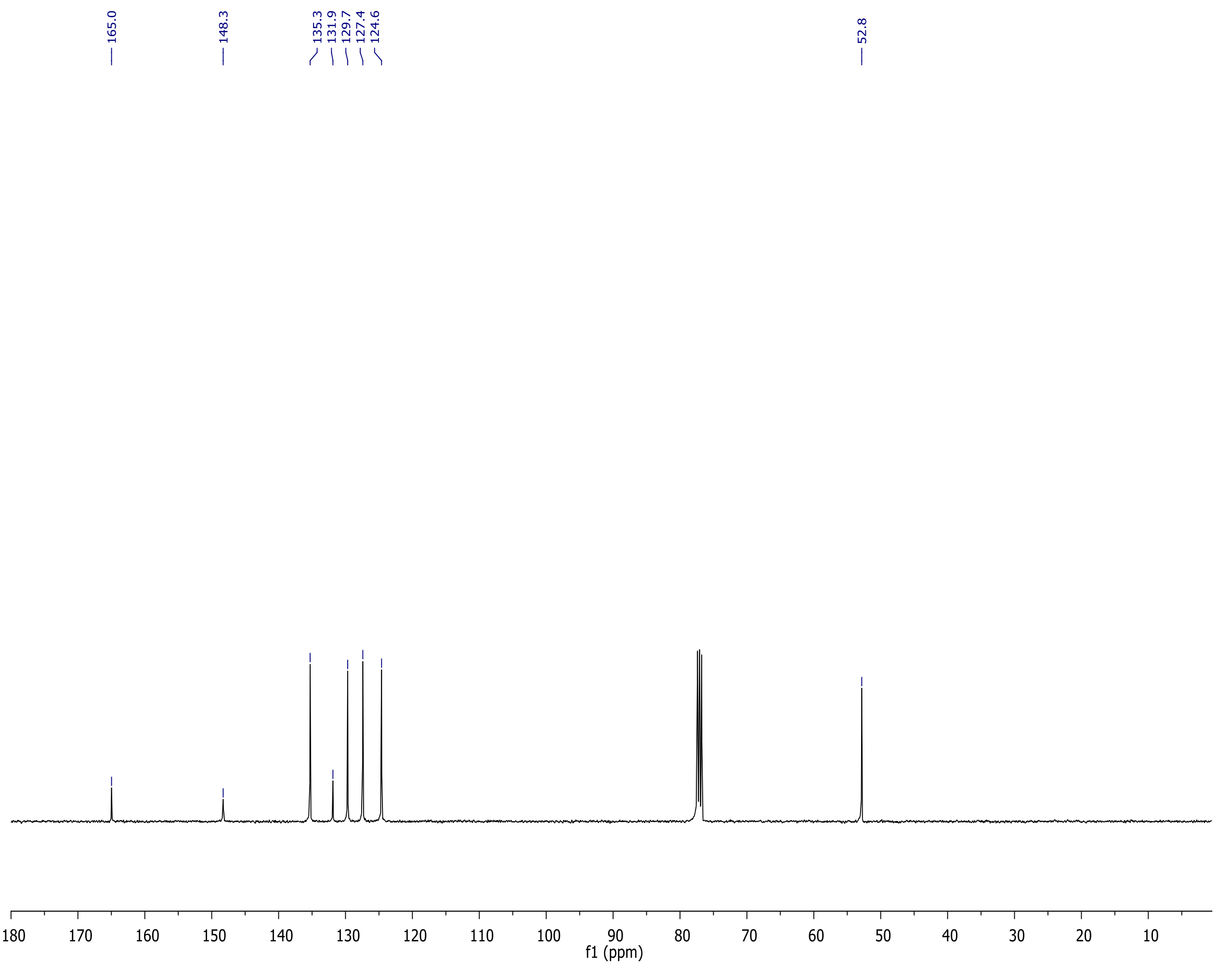

**$^1$H NMR spectra of Compound-3 in $CDCl_3$**

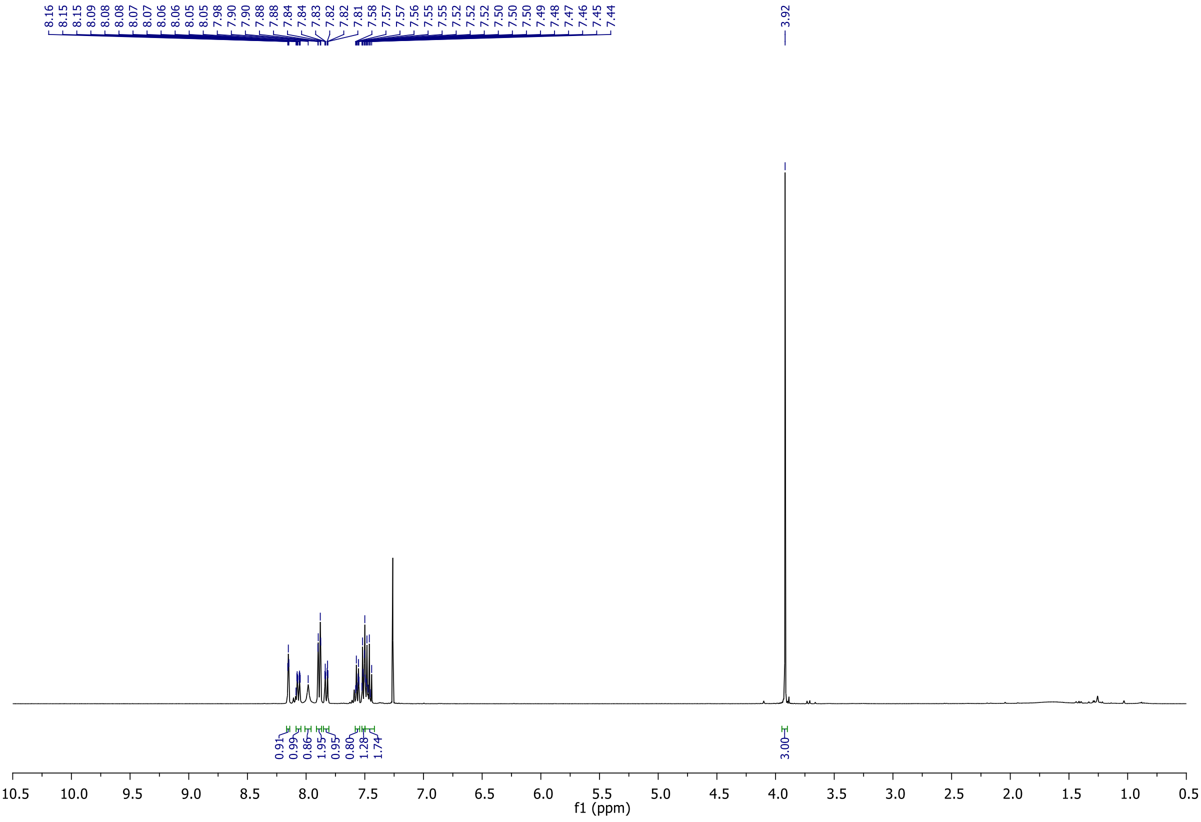

**$^{13}C$ NMR spectra of Compound-3 in $CDCl_3$**

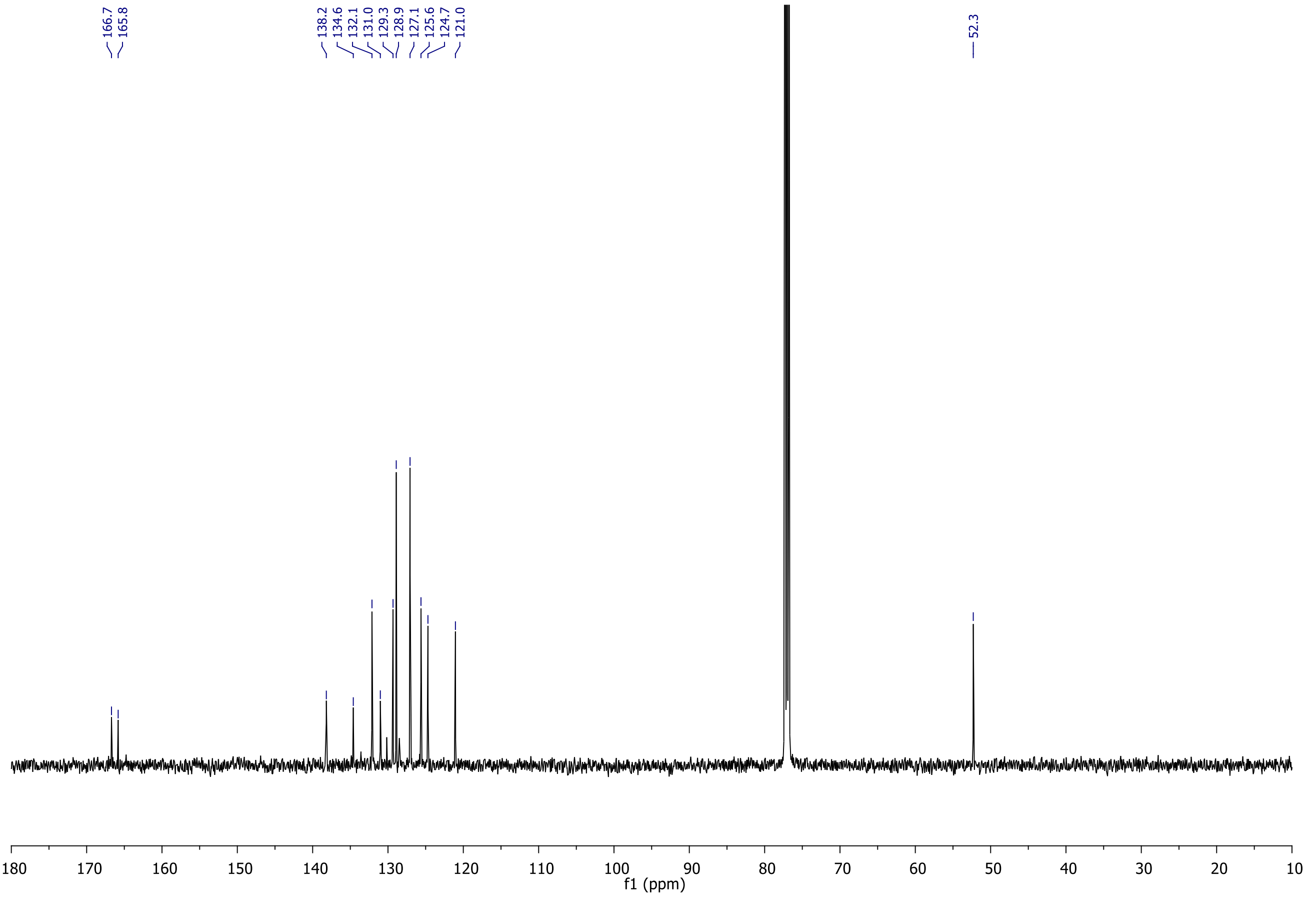


**HRMS spectrum of Compound-3**

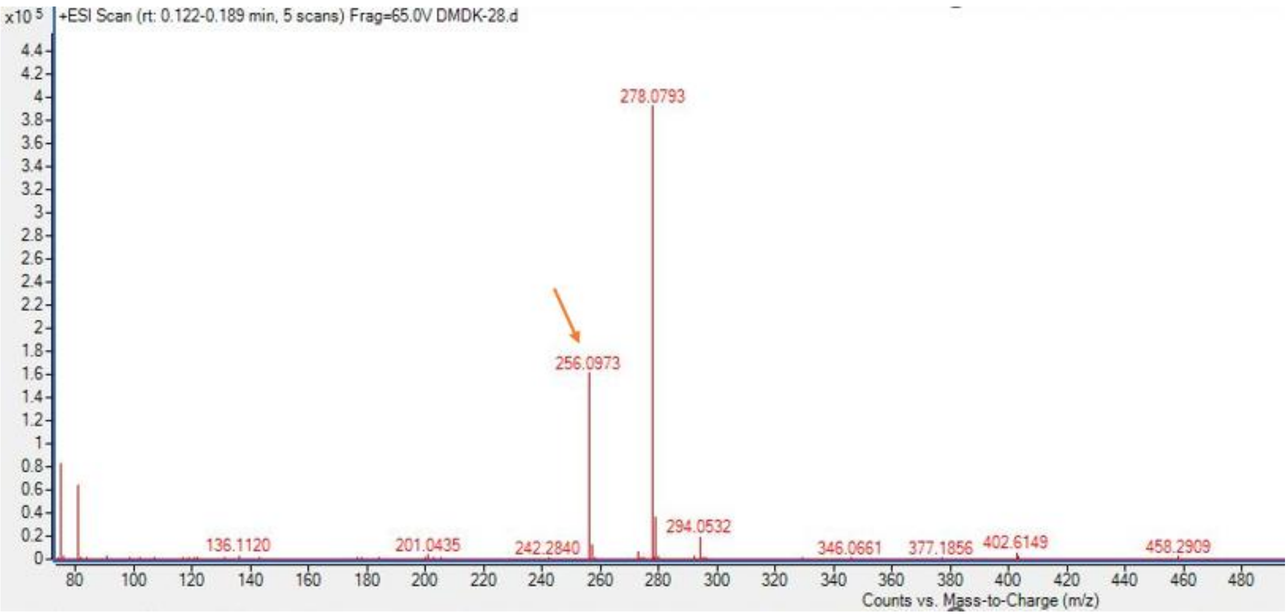


Calculated for $C_{15}H_{14}NO_3$ $(M+H)^+$: 256.0974, found 256.0973.

**[1]H NMR spectra of DM-01 in DMSO-$d_6$**

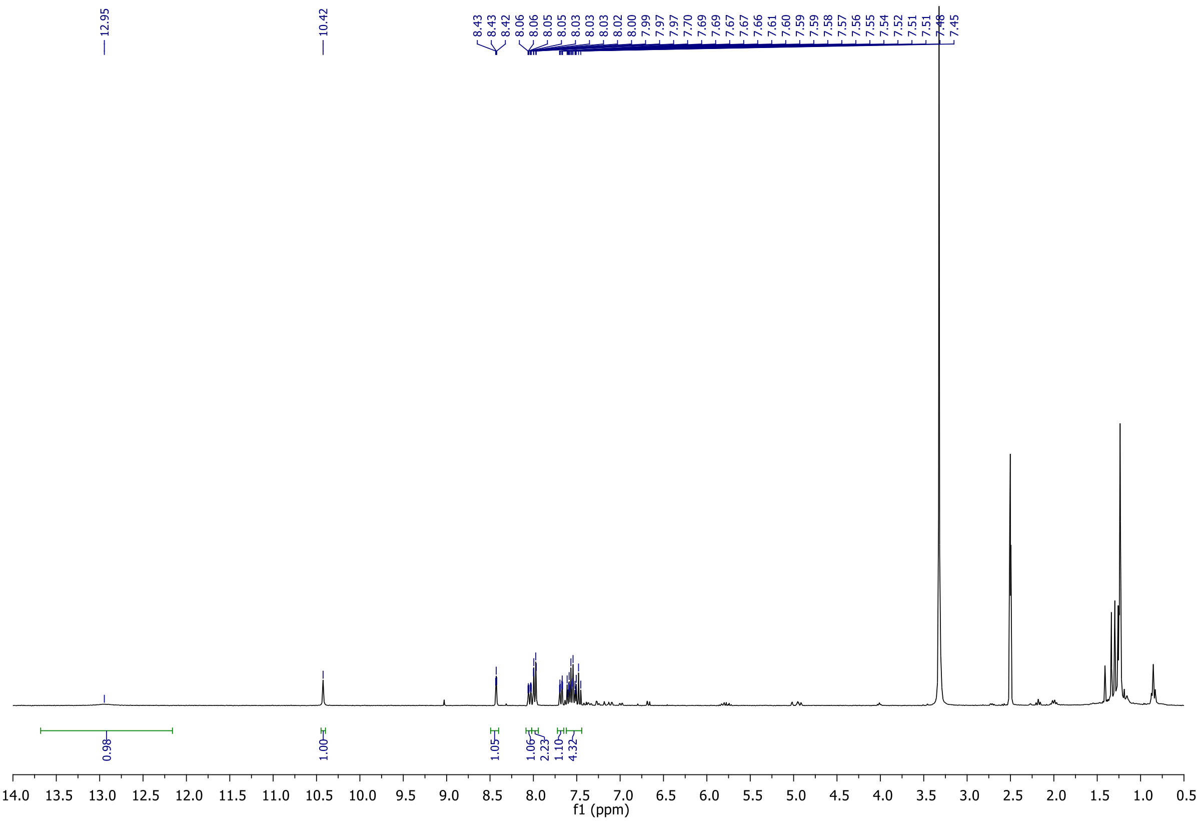

### $^{13}C$ NMR spectra of DM-01 in DMSO-$d_6$

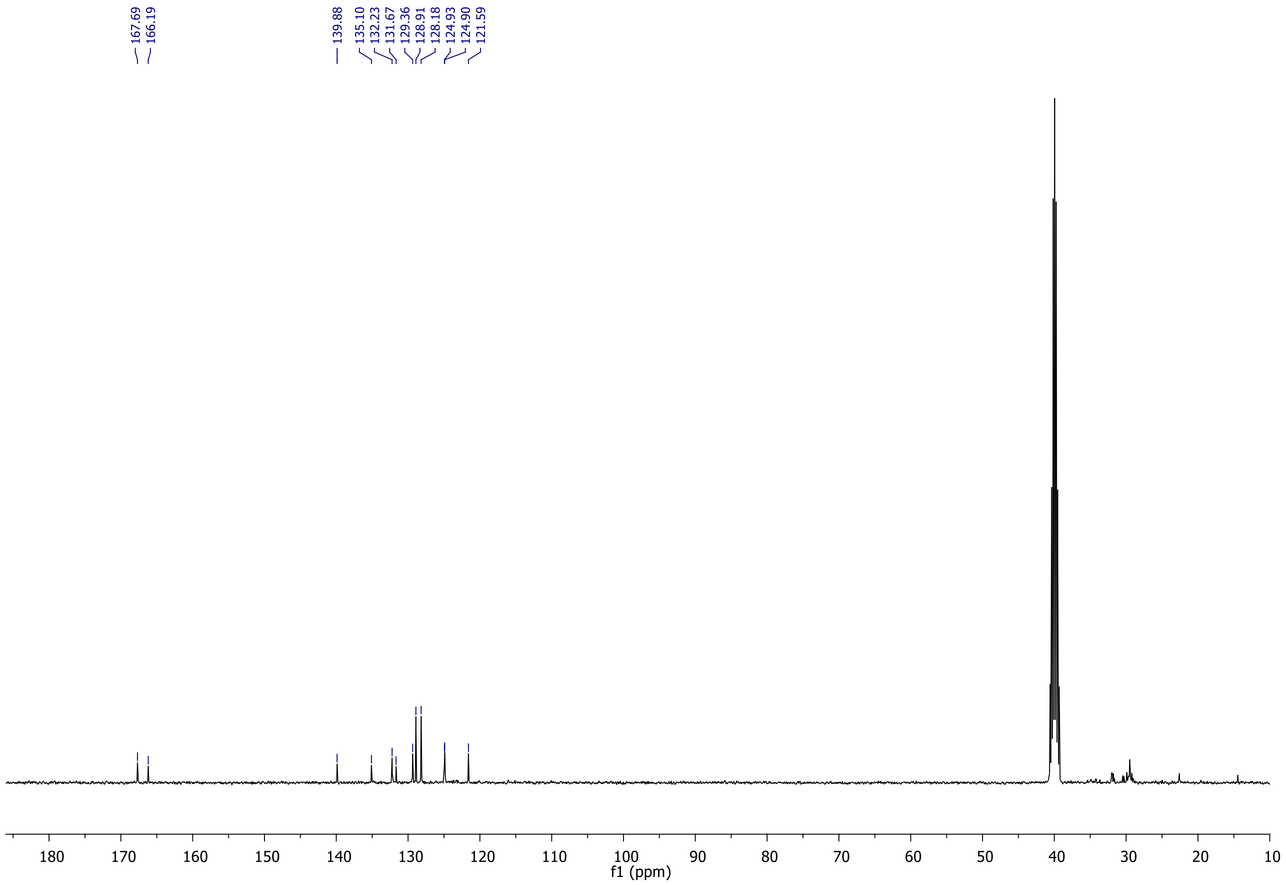


### HRMS spectrum of DM-01

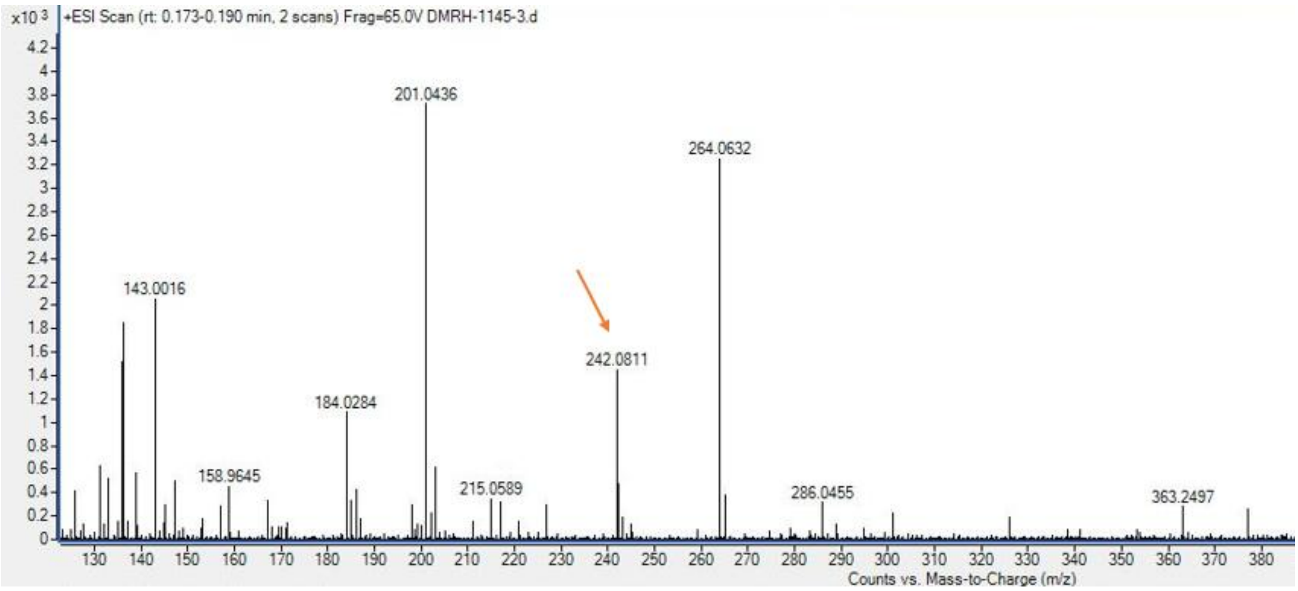


Calculated for $C_{14}H_{12}NO_3$ $(M+H)^+$: 242.0817, found 242.0811.

### $^1H$ NMR spectra of Compound-4 in $CDCl_3$

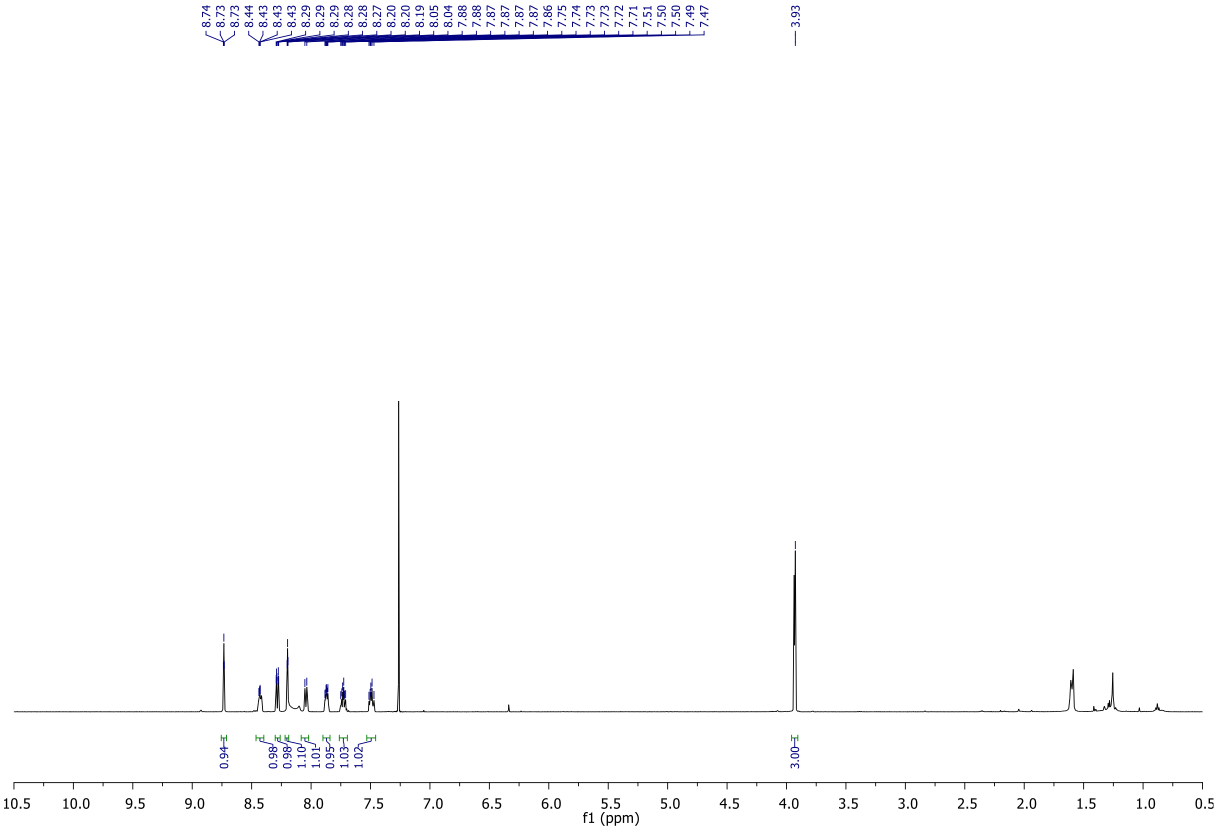

### $^{13}C$ NMR spectra of Compound-4 in $CDCl_3$

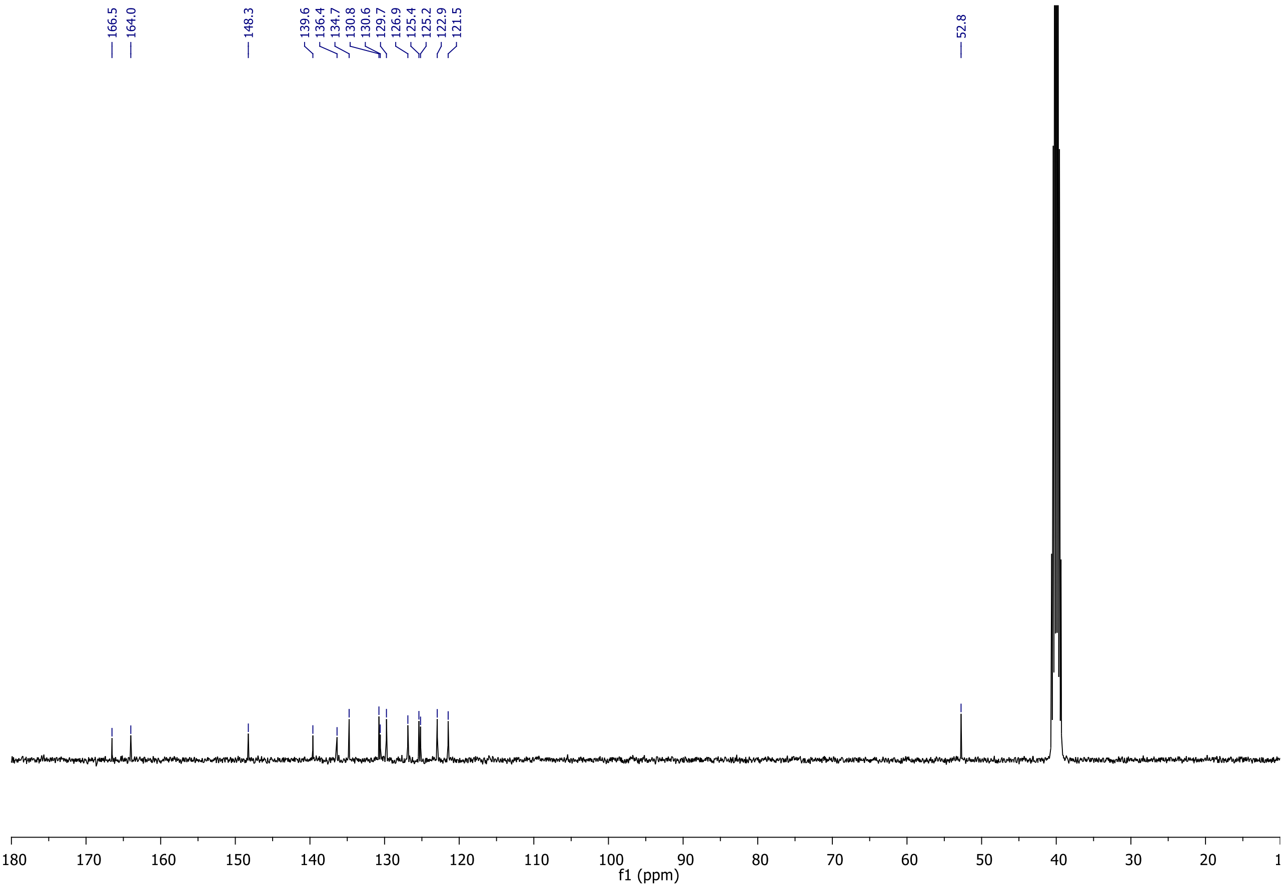


### HRMS spectrum of Compound-4

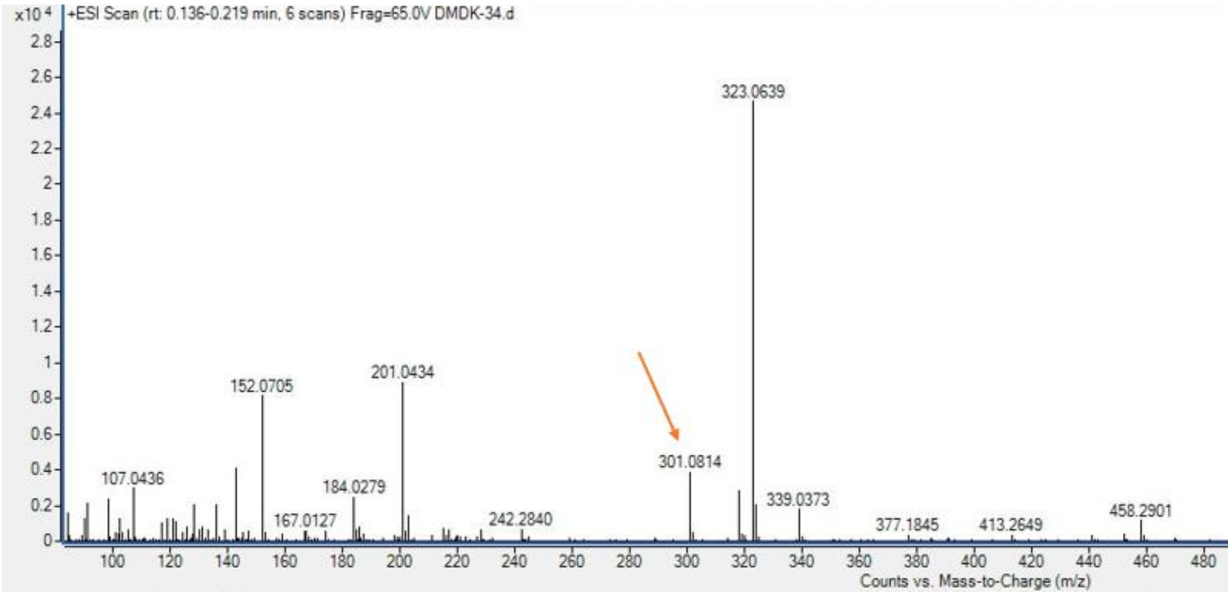


Calculated for $C_{15}H_{13}N_2O_5$ $(M+H)^+$: 301.0824, found 301.0814.

### $^{1}$H NMR spectra of DM-02 in DMSO-$d_6$

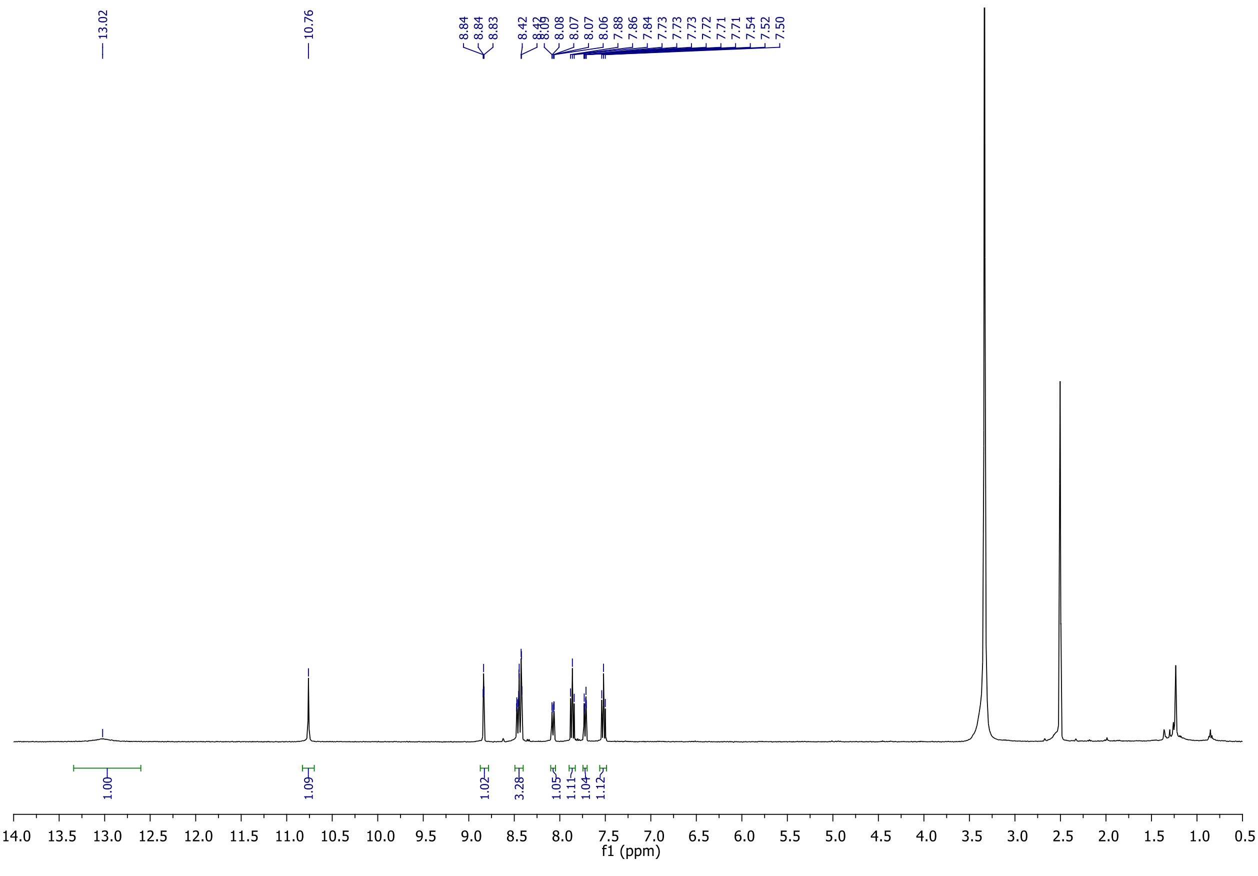

**$^{13}C$ NMR spectra of DM-02 in DMSO-$d_6$**

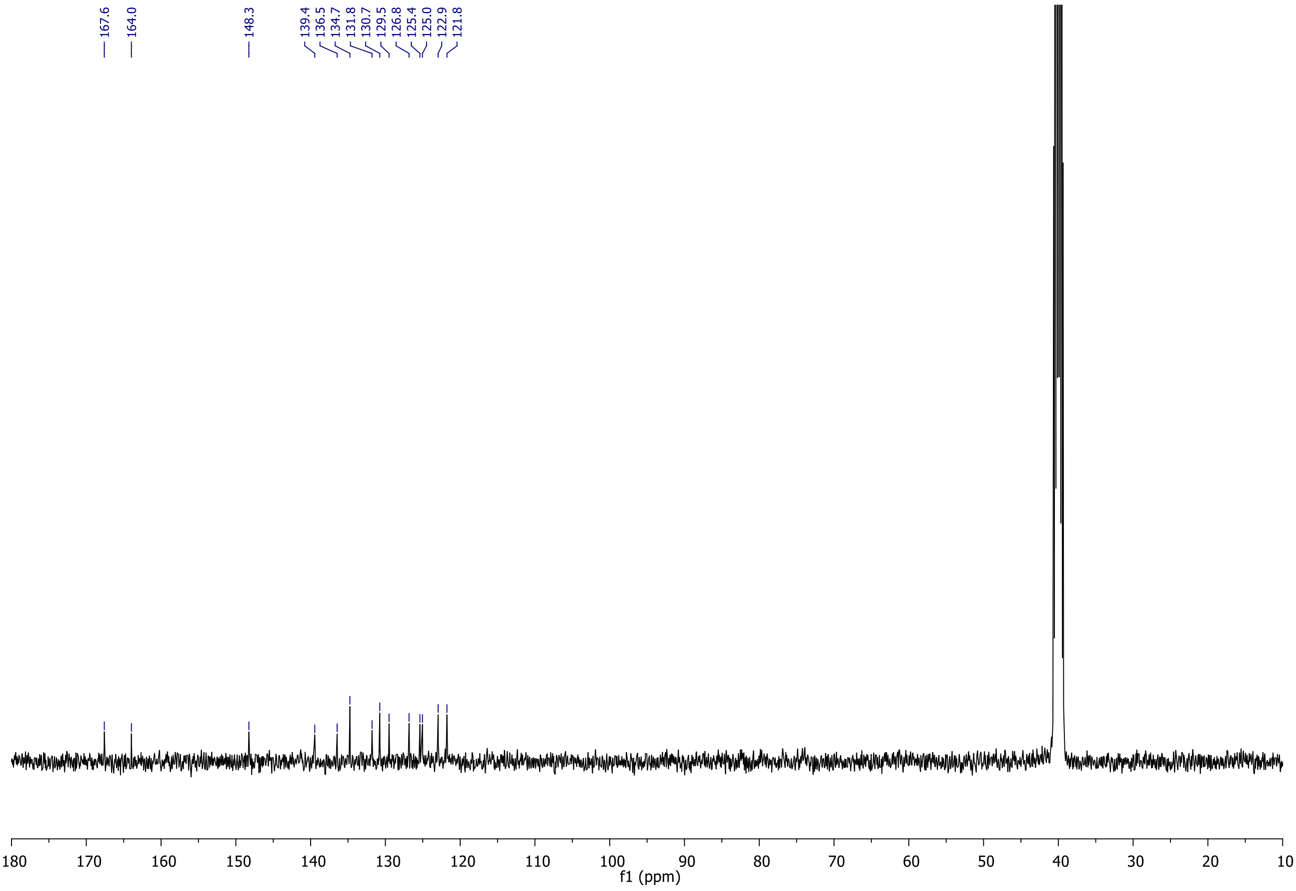


**HRMS spectrum of DM-02**

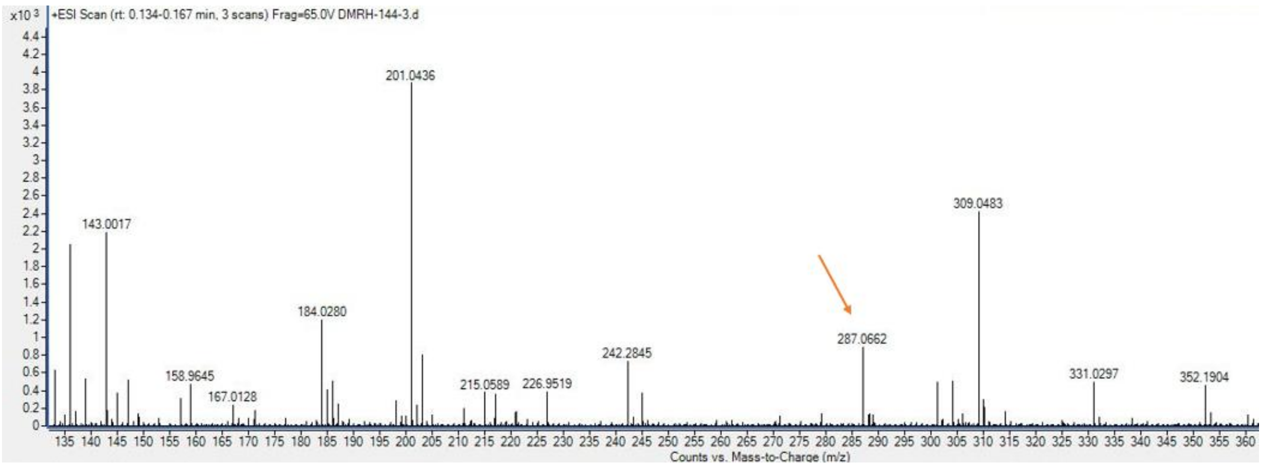


Calculated for $C_{14}H_{11}N_2O_5$ $(M+H)^+$: 287.0668, found 287.0662.

### $^{1}$H NMR spectra of DM-03 in DMSO-$d_6$

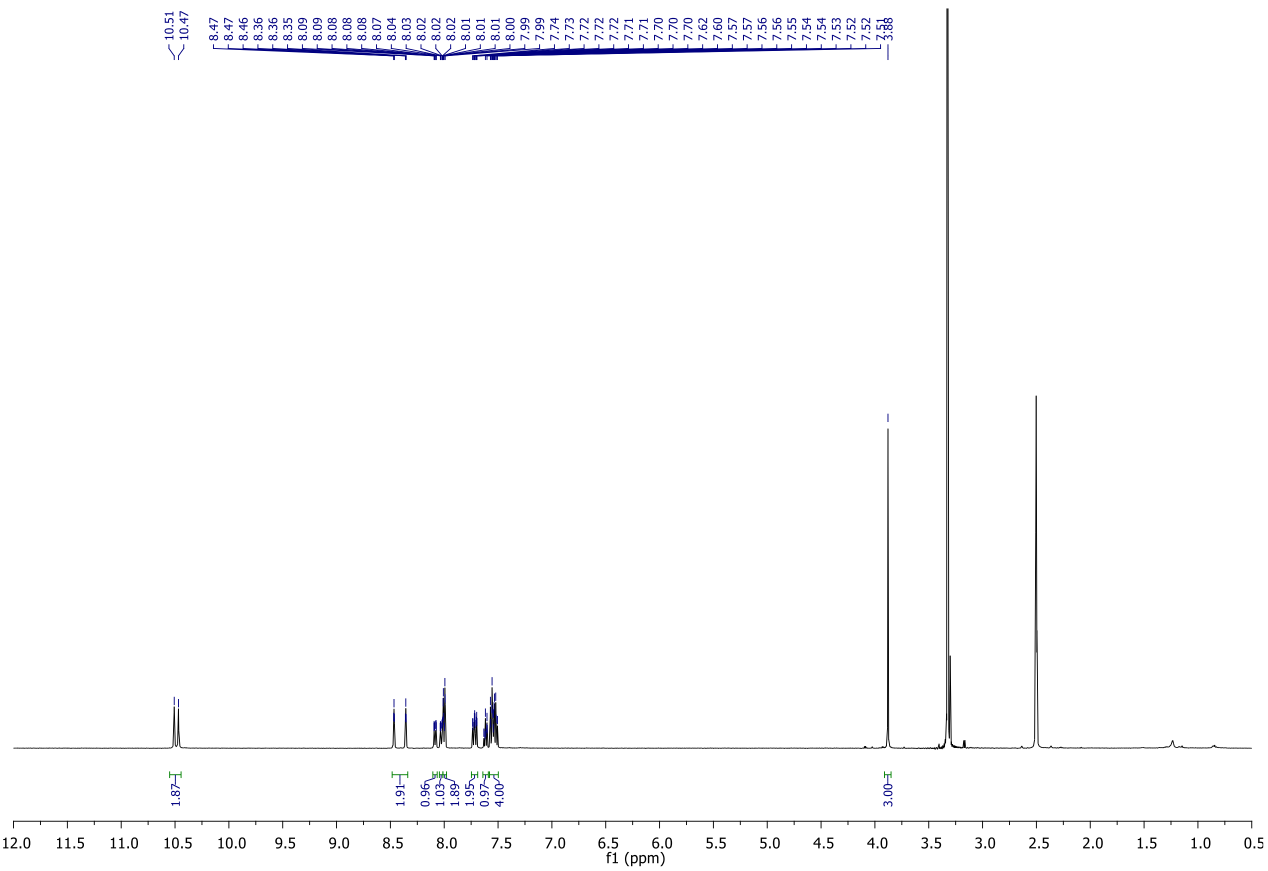

**$^{13}C$ NMR spectra of DM-03 in DMSO-$d_6$**

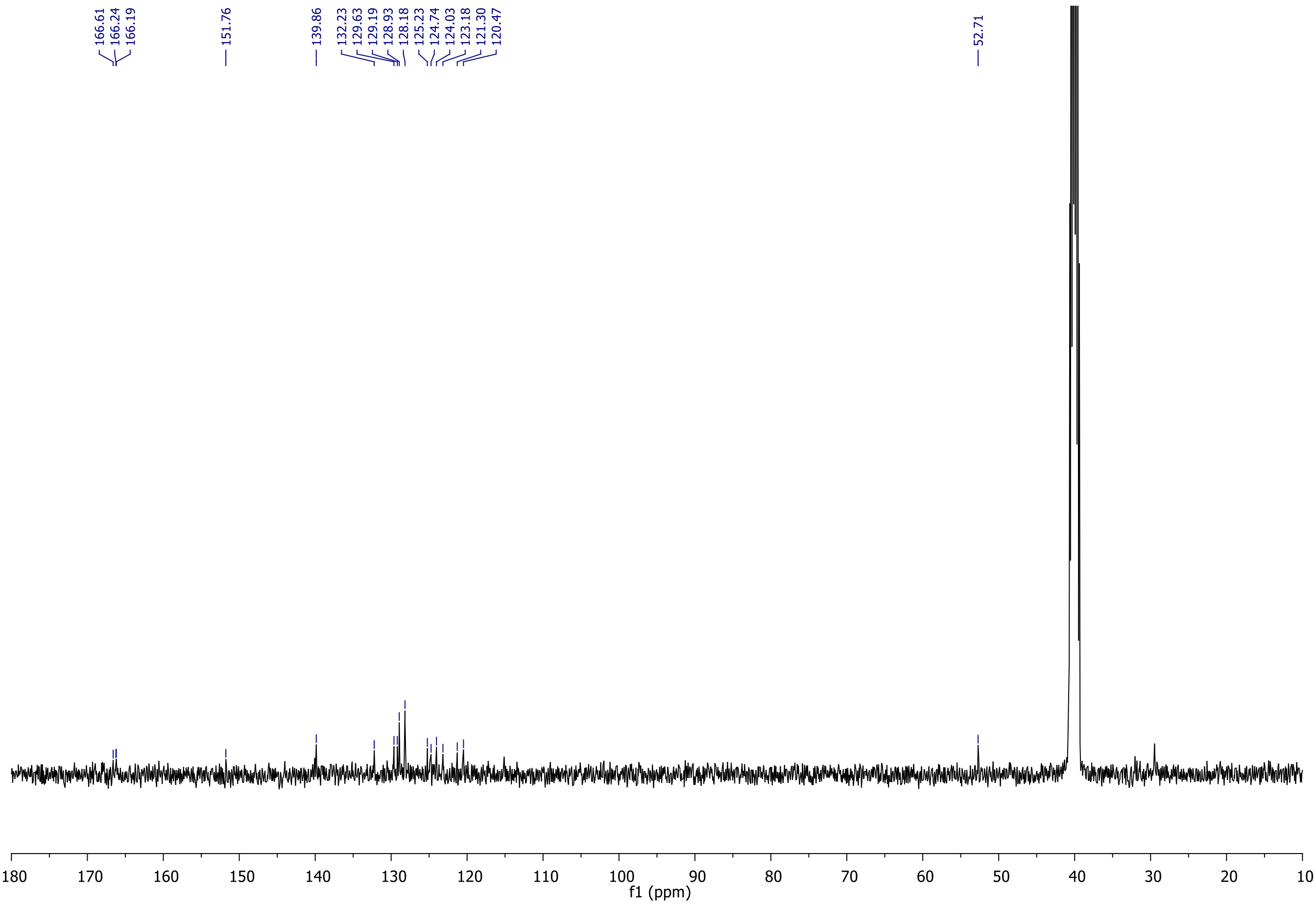


**HRMS spectrum of DM-03**

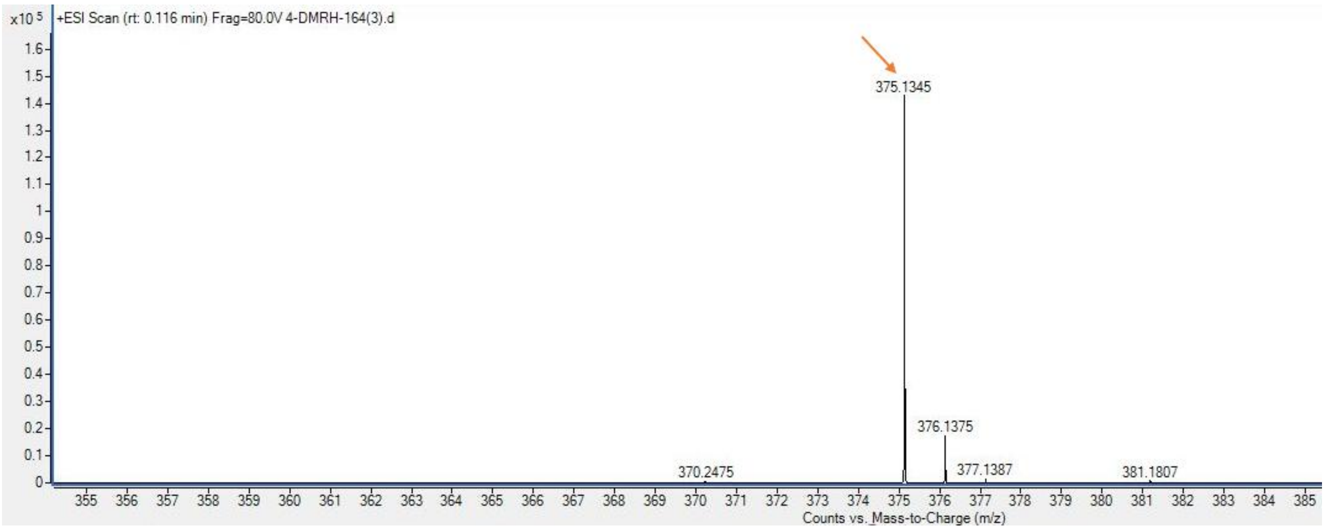


Calculated for $C_{22}H_{19}N_2O_4$ $(M+H)^+$: 375.1345, found 375.1345.

**$^{1}$H NMR spectra of DM-04 in DMSO-$d_6$**

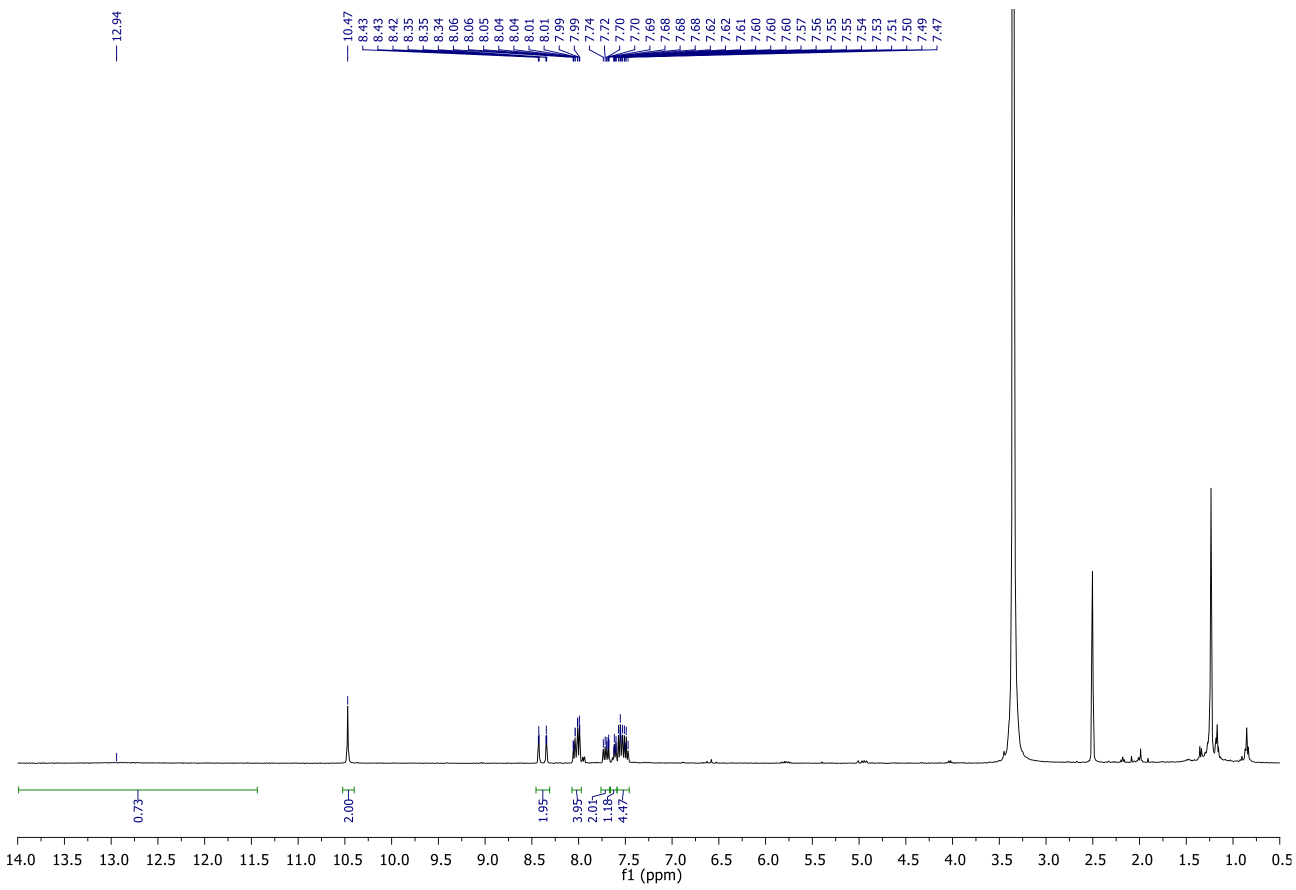

**$^{13}C$ NMR spectra of DM-04 in DMSO-$d_6$**

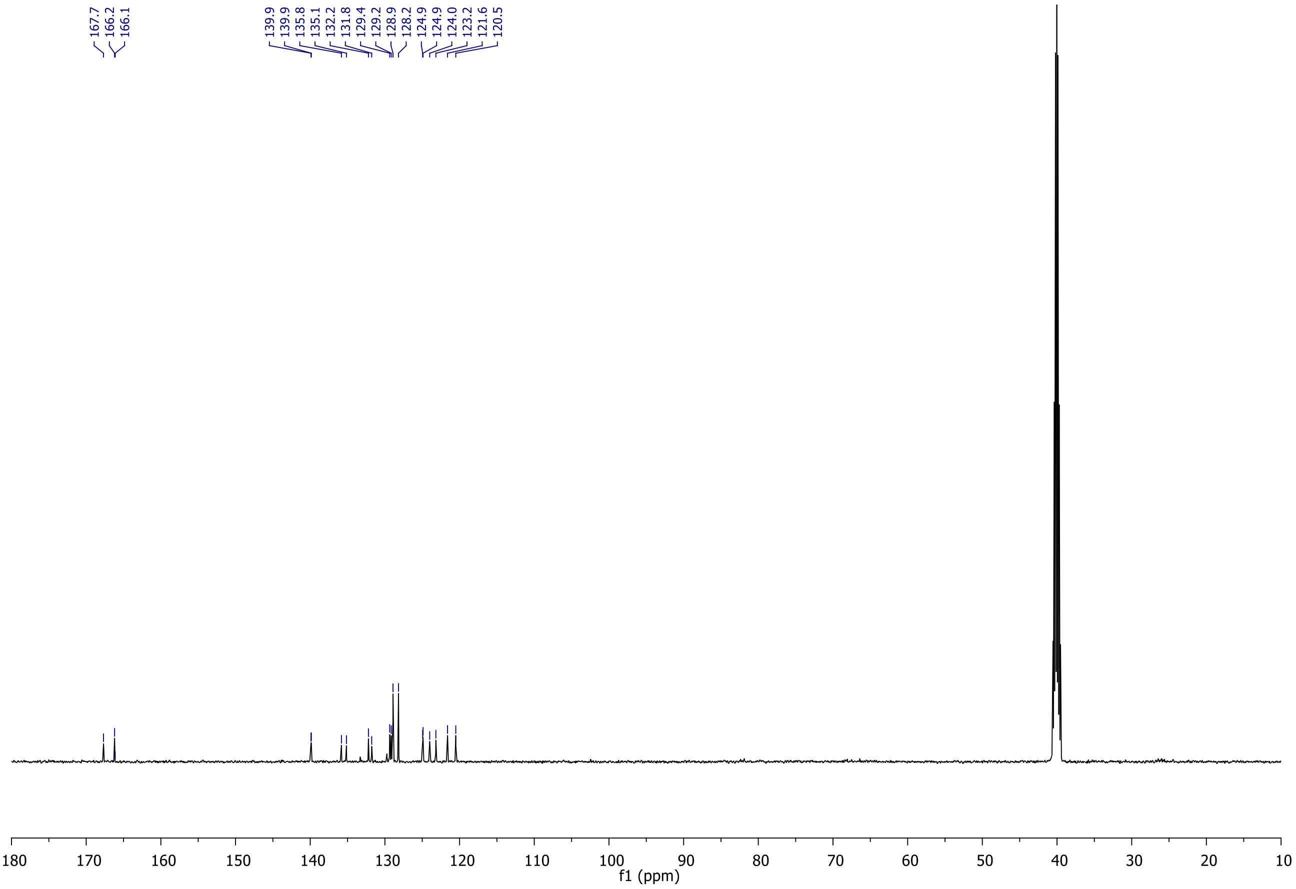


**HRMS spectrum of DM-04**

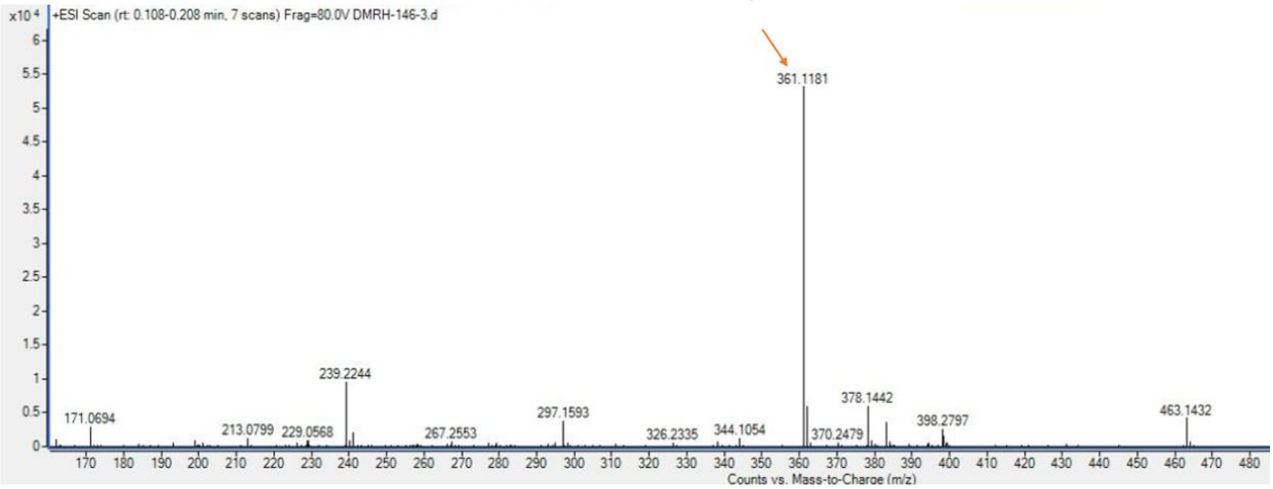


Calculated for $C_{21}H_{17}N_2O_4$ $(M+H)^+$: 361.1188, found 361.1181.
