## Supplementary Data 6 for "A tribenzamide small molecule with retinoic acid-like activity promotes axon growth and motor recovery after spinal cord injury"

### Maity Lab CSIR-IICT

#### <Sample Information>

| | | | |
|---|---|---|---|
| Sample Name | : DMRH-04 | | |
| Sample ID | : DMRH-04 | | |
| Data Filename | : DMRH-04-Purity.lcd | | |
| Method Filename | : Purity analysis10-90h.lcm | | |
| Batch Filename | : DMRH-04.lcb | | |
| Vial # | : 1-4 | Sample Type | : Unknown |
| Injection Volume | : 3 uL | | |
| Date Acquired | : 11/12/2025 4:17:23 PM | Acquired by | : System Administrator |
| Date Processed | : 11/12/2025 4:44:51 PM | Processed by | : System Administrator |

#### <Chromatogram>

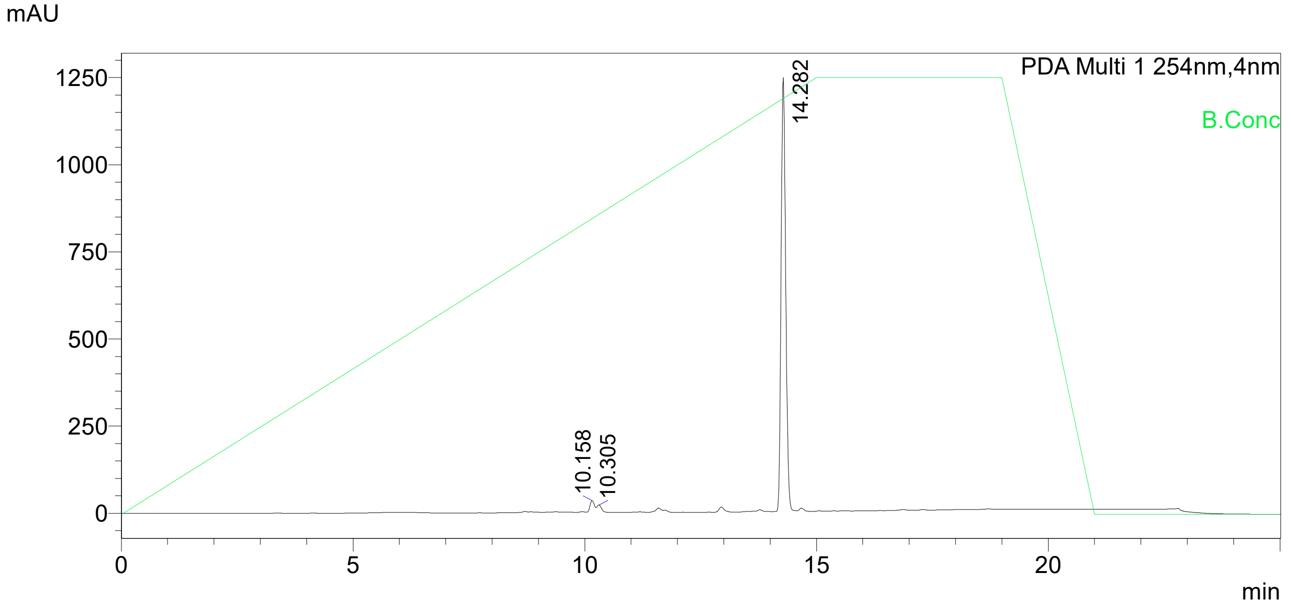


UV Spectrum

| | |
|---|---|
| Peak# | : 3 |
| Retention Time | : 14.282 min |
| Compound Name | : |
| Spectrum Operation | : None |

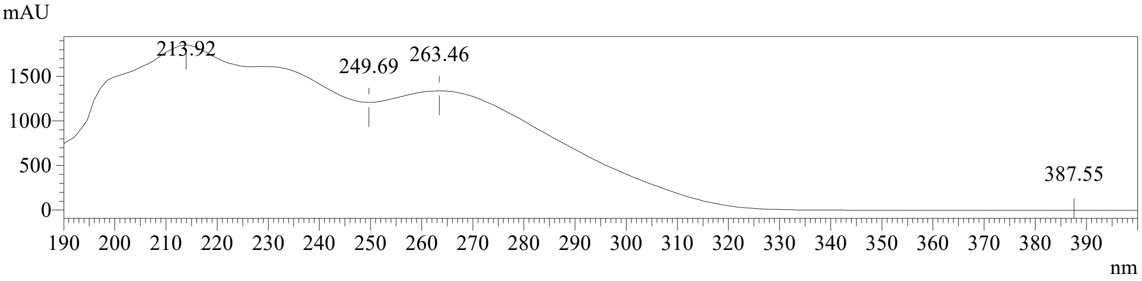


Peak Table

PDA Ch1 254nm

| Peak# | Ret. Time | Area | Height | Area% |
|---|---|---|---|---|
| 1 | 10.158 | 145158 | 25689 | 1.662 |
| 2 | 10.305 | 58965 | 11704 | 0.675 |
| 3 | 14.282 | 8528631 | 1243661 | 97.663 |
| Total | | 8732753 | 1281053 | 100.000 |
