## Supplementary material for "A tribenzamide small molecule with retinoic acid-like activity promotes axon growth and motor recovery after spinal cord injury": All supplementary figures and tables

### Supplementary Information

***Supplementary Tables***

- Table S1 - Percentage yield for the compounds used in the study
- Table S2 - Luciferase assay luminometer readings
- Table S3 - Toxicity analysis
- Table S4 - Average neurite outgrowth - Neuro2a (Raw file attached separately)
- Table S5 - Average neurite outgrowth – Primary Neuron (Raw file attached separately)
- Table S6 - RNA seq (Raw file attached separately)
- Table S7 - Hip rise values
- Table S8 - Mean GFP Intensity of Axons into the glial scar
- Table S9 - Pax6+ Cell Quantification during Neural Induction

***Supplementary figures***

- Figure S1 - Molecular docking and interaction profile of RARβ with native retinoic acid and candidate ligands
- Figure S2 - Molecular docking replicates and 100 ns molecular dynamics (MD) simulation analyses of RARβ system
- Figure S3 - HPLC analysis demonstrated that stability of 10 μM of DM04 is unchanged after 60 min irradiation of UV light (360 nm)
- Figure S4 - HPLC analysis demonstrated that stability of 10 μM of DM04 is unchanged after 1mM peroxide treatment for 3 hr
- Figure S5 - UV-Vis spectra of DM04 with increasing concentration in water
- Figure S6 - . Size distribution of DM04 at different concentration in water
- Figure S7 - DM04 failed to bind to the RAR elements from the pGL3-RARE-Luciferase plasmid
- Figure S8 - DM04 significantly increases neurite outgrowth *in vitro* in N2A cells.

***Supplementary Data***

- Data S1 - Behavior videos (Attached separately)

  Supplementary Video 1 – 1wpi (DMSO)

  Supplementary Video 1 – 1wpi (DM04)

  Supplementary Video 1 – 3wpi (DMSO)

  Supplementary Video 1 – 3wpi (DM04)
- Data S5 and S6 - NMR data (Attached).

| Compound | % yield |
|---|---|
| **Compound-1** | 96 |
| **Compound-2** | 98 |
| **Compound-3** | 85 |
| **DM01** | 98 |
| **Compound-4** | 85 |
| **DM02** | 98 |
| **Compound-5** | 98 |
| **DM03** | 78 |
| **DM04** | 98 |

**Table S1 : Percentage yield for the compounds used in the study.**

| Replicate | Treatment | Timepoint | Reading 1 | Reading 2 | Reading 3 |
|---|---|---|---|---|---|
| Replicate 1 | RA | 48 h | 7270 | 8302 | 10008 |
| | RA | 24 h | 31256 | 35948 | 35151 |
| | RA | 12 h | 12955 | 9814 | 9648 |
| | RA | 0 h | 281 | 346 | 396 |
| | DM04 | 48 h | 639 | 393 | 314 |
| | DM04 | 24 h | 2628 | 3409 | 4369 |
| | DM04 | 12 h | 2629 | 2309 | 2053 |
| | DM04 | 0 h | 389 | 437 | 392 |
| | DMSO | 48 h | 337 | 307 | 292 |
| | DMSO | 24 h | 1321 | 942 | 1197 |
| | DMSO | 12 h | 902 | 1091 | 1198 |
| | DMSO | 0 h | 320 | 315 | 249 |
| Replicate 2 | RA | 48 h | 26981 | 34727 | 29580 |
| | RA | 24 h | 8339 | 17741 | 4168 |
| | RA | 12 h | 787 | 495 | 602 |
| | RA | 0 h | 367 | 359 | 698 |
| | DM04 | 48 h | 852 | 774 | 726 |
| | DM04 | 24 h | 517 | 587 | 675 |
| | DM04 | 12 h | 394 | 405 | 404 |
| | DM04 | 0 h | 626 | 404 | 350 |
| | DMSO | 48 h | 701 | 809 | 718 |
| | DMSO | 24 h | 516 | 528 | 453 |
| | DMSO | 12 h | 339 | 486 | 351 |
| | DMSO | 0 h | 367 | 366 | 522 |
| Replicate 3 | RA | 48 h | 27649 | 16575 | 26906 |
| | RA | 24 h | 45217 | 25018 | 39482 |
| | RA | 12 h | 11742 | 7229 | 11403 |
| | RA | 0 h | 899 | 546 | 968 |
| | DM04 | 48 h | 1831 | 1236 | 1135 |
| | DM04 | 24 h | 1512 | 997 | 1798 |
| | DM04 | 12 h | 1656 | 1065 | 1253 |
| | DM04 | 0 h | 1025 | 687 | 1057 |
| | DMSO | 48 h | 1381 | 996 | 743 |
| | DMSO | 24 h | 1214 | 905 | 669 |
| | DMSO | 12 h | 1060 | 883 | 487 |
| | DMSO | 0 h | 684 | 553 | 424 |

#### Table S2 – Luciferase binding : Luminometer readings

| | Replicate | DMSO | DM04 | RA | DM04 + RA |
|---|---|---|---|---|---|
| **100µM** | 1 | 95 | 83.0986 | 38.6364 | 68.1818 |
| | 2 | 62.0915 | 72.6444 | 61.039 | 53.8462 |
| | 3 | 76.6447 | 71.2531 | 78.0669 | 74.6888 |
| **50µM** | 1 | 78.6008 | 98.5437 | 99.0385 | 64.7059 |
| | 2 | 65.4135 | 73.1518 | 67.019 | 69.1176 |
| | 3 | 71.134 | 68.6047 | 61.413 | 77.6573 |
| **25µM** | 1 | 71 | 100 | 78.0952 | 98.4615 |
| | 2 | 59.3625 | 64.7887 | 79.7059 | 69.7368 |
| | 3 | 88.4615 | 79.6089 | 96.8137 | 73.8854 |
| **10µM** | 1 | 51 | 86.3014 | 75 | 70.8661 |
| | 2 | 56.0784 | 100 | 92.0502 | 66.9492 |
| | 3 | 95.671 | 100 | 68.6047 | 71.3115 |

**Table S3 – Toxicity analysis**

| **100uM** | **DMSO** | **DM04** | **RA** | **DM04+RA** |
|---|---|---|---|---|
| Replicate 1 | 125.68 | 139.65 | 119.39 | 125.83 |
| Replicate 2 | 133.44 | 179.98 | 180.64 | 197.56 |
| Replicate 3 | 165.96 | 187.34 | 192.51 | 227.18 |
| Replicate 4 | 159.24 | 182.06 | 188.43 | 227.31 |
| **50uM** | **DMSO** | **DM04** | **RA** | **DM04+RA** |
| Replicate 1 | 145.86 | 161.9 | 218.75 | 211.16 |
| Replicate 2 | 170.35 | 190.16 | 196.06 | 202.05 |
| Replicate 3 | 189.3 | 193.04 | 201.06 | 164.92 |
| Replicate 4 | 151.97 | 160.06 | 152.87 | 203.33 |
| **25uM** | **DMSO** | **DM04** | **RA** | **DM04+RA** |
| Replicate 1 | 169.12 | 219.73 | 250.26 | 267.23 |
| Replicate 2 | 178.07 | 211.54 | 191.83 | 196.42 |
| Replicate 3 | 191.84 | 222.59 | 170.27 | 140.32 |
| Replicate 4 | 149.16 | 217.56 | 193.6 | 194.09 |
| **10uM** | **DMSO** | **DM04** | **RA** | **DM04+RA** |
| Replicate 1 | 183.37 | 315.37 | 311.09 | 252.03 |
| Replicate 2 | 166.89 | 278.95 | 201.27 | 196.42 |
| Replicate 3 | 189.79 | 272.62 | 154.24 | 223.2 |
| Replicate 4 | 168.58 | 269 | 146.87 | 216.76 |

**Table S4 – Average neurite outgrowth - Neuro2a**

| | DMSO | DM04 |
|---|---|---|
| Replicate 1 | 150.3273 | 264.8801 |
| Replicate 2 | 120.3068 | 298.9996 |
| Replicate 3 | 161.9661 | 295.0038 |
| Replicate 4 | 150.41 | 290.7389 |

**Table S5 – Average neurite outgrowth – Primary Neuron**

| Group | Animal | Frame | Reading 1 (cm) | Reading 2 (cm) | Reading 3 (cm) |
|---|---|---|---|---|---|
| Uninjured | Animal 1 | Frame 1 | 4.348 | 4.478 | 4.524 |
| Uninjured | Animal 1 | Frame 2 | 4.654 | 4.696 | 4.783 |
| Uninjured | Animal 2 | Frame 1 | 4.436 | 4.565 | 4.826 |
| Uninjured | Animal 2 | Frame 2 | 4.657 | 4.696 | 4.611 |
| Uninjured | Animal 3 | Frame 1 | 4.565 | 4.522 | 4.739 |
| Uninjured | Animal 3 | Frame 2 | 4.699 | 4.565 | 4.565 |
| DM04 | Animal 1 | Frame 1 | 4.478 | 4.467 | 4.478 |
| DM04 | Animal 1 | Frame 2 | 4.262 | 4.478 | 4.522 |
| DM04 | Animal 2 | Frame 1 | 4.174 | 4.089 | 4.131 |
| DM04 | Animal 2 | Frame 2 | 4.522 | 4.478 | 4.522 |
| DM04 | Animal 3 | Frame 1 | 3.523 | 3.479 | 3.696 |
| DM04 | Animal 3 | Frame 2 | 4.35 | 4.392 | 4.565 |
| DMSO | Animal 1 | Frame 1 | 2.087 | 2 | 2.13 |
| DMSO | Animal 1 | Frame 2 | 2.348 | 2.306 | 2.304 |
| DMSO | Animal 2 | Frame 1 | 2.305 | 2.219 | 2.263 |
| DMSO | Animal 2 | Frame 2 | 2 | 2.13 | 2.174 |
| DMSO | Animal 3 | Frame 1 | 2.174 | 2.174 | 2.131 |
| DMSO | Animal 3 | Frame 2 | 2.043 | 2.13 | 2.087 |

**Table S7 – Hip rise values**

| Sample | Animal Number | Area | Mean Intensity |
|---|---|---|---|
| **DMSO** | 1 | 39118.414 | p |
| **DMSO** | 2 | 39118.414 | 3.599 |
| **DMSO** | 3 | 39118.414 | 3.215 |
| **DM04** | 1 | 39118.414 | 5.035 |
| **DM04** | 2 | 39118.414 | 8.089 |
| **DM04** | 3 | 39118.414 | 4.501 |
| **DM04** | 4 | 39118.414 | 6.234 |

**Table S8 – Mean GFP Intensity of Axons into the glial scar**

| | Control 1 | Control 2 | Control 3 | DM04 1 | DM04 2 | DM04 3 |
|---|---|---|---|---|---|---|
| ***DM04+RA*** | 8 | 22.7 | | 5.4 | 0.9 | |
| ***DM04*** | 4.9 | 10 | | 8.8 | 15.45 | |
| ***Compound C+DM04*** | 0 | 0 | | 59.4 | 76.6 | |
| ***Compound C+RA*** | 0 | 0.86 | | 8.8 | 16.2 | |

**Table S9 – Pax6+ cells% - Quantification during Neural Induction**

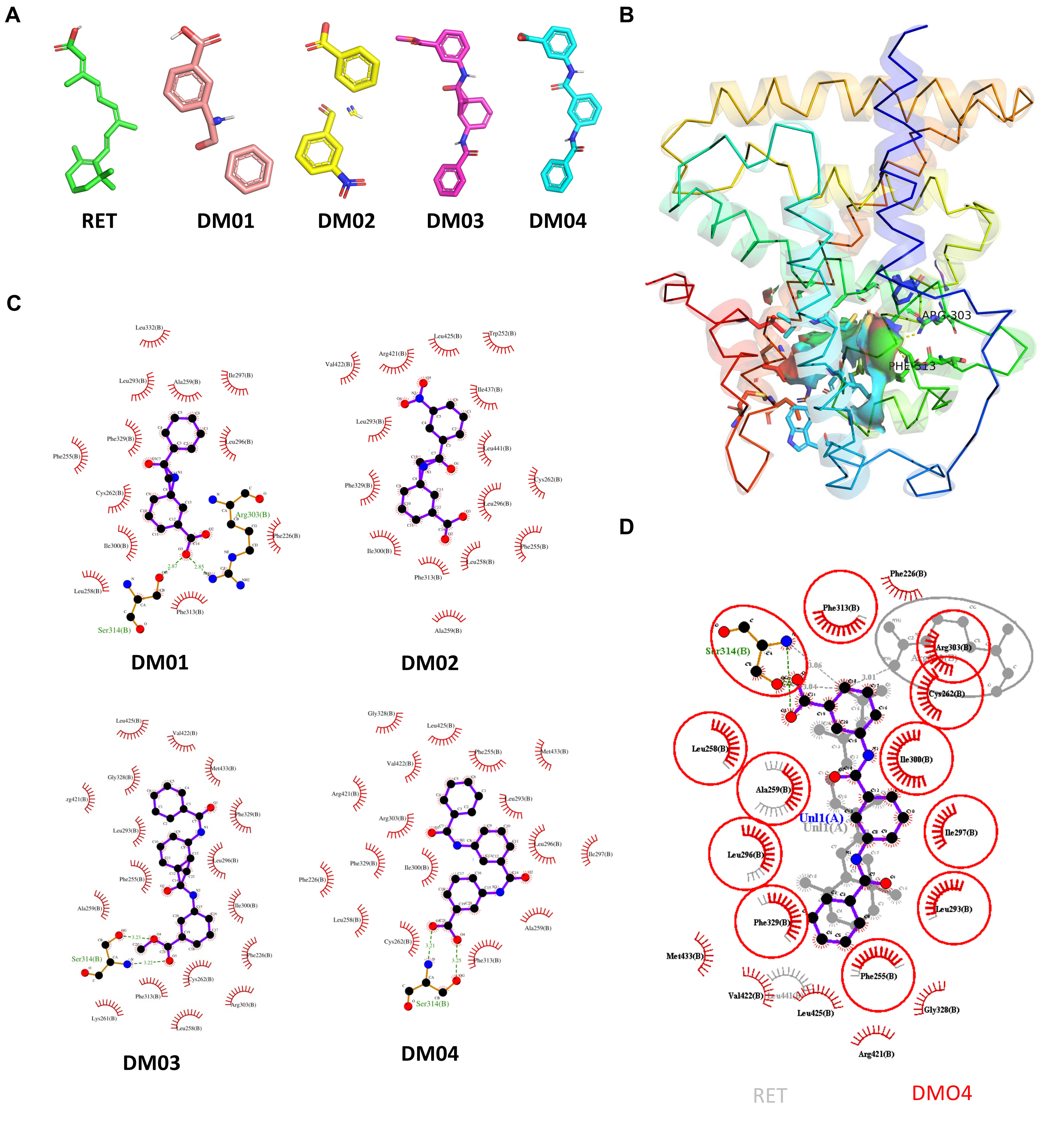

**Figure S1. Molecular docking and interaction profile of RARβ with native retinoic acid and candidate ligands. (A**) Three-dimensional conformations of the native control retinoic acid (RET) and four selected small-molecule candidate ligands (DM01, DM02, DM03, and DM04). **(B)** Three-dimensional ribbon representation of the RARβ receptor, showing the active ligand-binding pocket and key interacting residues (including Arg303 and Phe313).**(C)** Two-dimensional Lig-Plot interaction diagrams illustrating the binding profiles of DM01, DM02, DM03, and DM04 within the RARβ binding site. Green dashed lines denote conventional hydrogen bonds along with bond distances (Å), while red spiked arcs indicate active site residues involved in hydrophobic contacts.**(D)** Comparative superimposed 2D interaction map of native retinoic acid (RET, grey) versus the top-scoring candidate compound (DM04, black/red), highlighting shared hydrophobic and hydrogen-bonding contact residues inside the binding pocket.

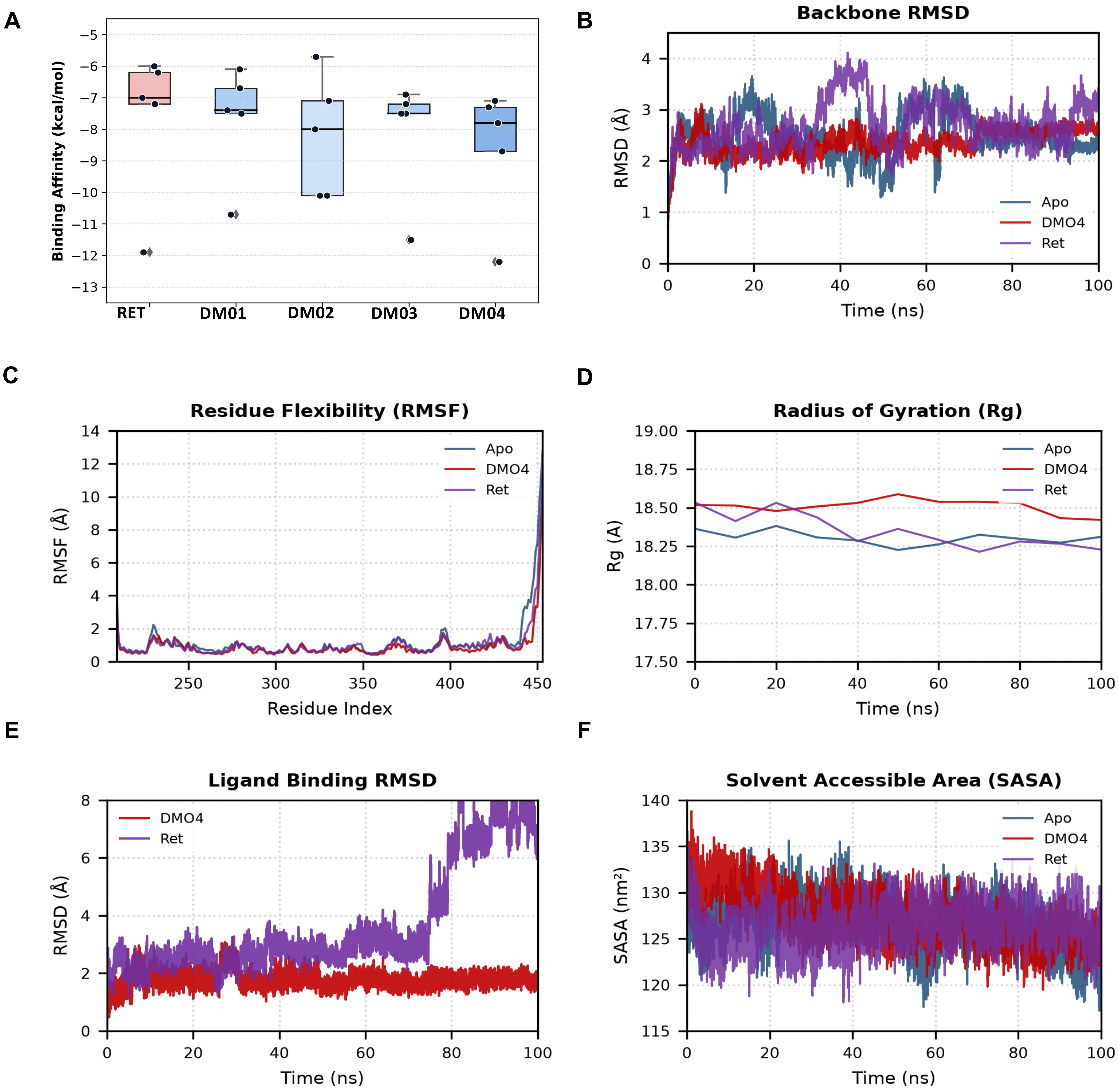

**Figure S2. Molecular docking replicates and 100 ns molecular dynamics (MD) simulation analyses of RARβ systems.(A)** Box plot depicting the binding affinity scores (kcal/mol) from 5 docking replicates for the control retinoic acid (RET) and screened candidate compounds (DM01–DM04), showing median values and individual run distributions.**(B)** Time evolution of protein backbone Root Mean Square Deviation (RMSD, Å) across the 100 ns MD trajectory for apo-RARβ (Apo, blue), the retinoic acid complex (Ret, purple), and the lead candidate complex (DMO4, red).**(C)** Root Mean Square Fluctuation (RMSF, Å) per residue index, highlighting structural flexibility and local residue dynamics for Apo, Ret, and DMO4 systems over the simulation period.**(D)** Radius of Gyration (Rg, Å) as a function of time (ns), illustrating overall protein compactness and conformational stability across the three systems.**(E)** Ligand Root Mean Square Deviation (RMSD, Å) assessing positional and conformational stability within the binding pocket for DMO4 (red) compared to native Ret (purple) throughout the 100 ns trajectory.**(F)** Solvent Accessible Surface Area (SASA, $nm^2$) trajectory profiles of Apo, Ret, and DMO4 complexes, representing solvent exposure and folding stability over 100 ns.

A

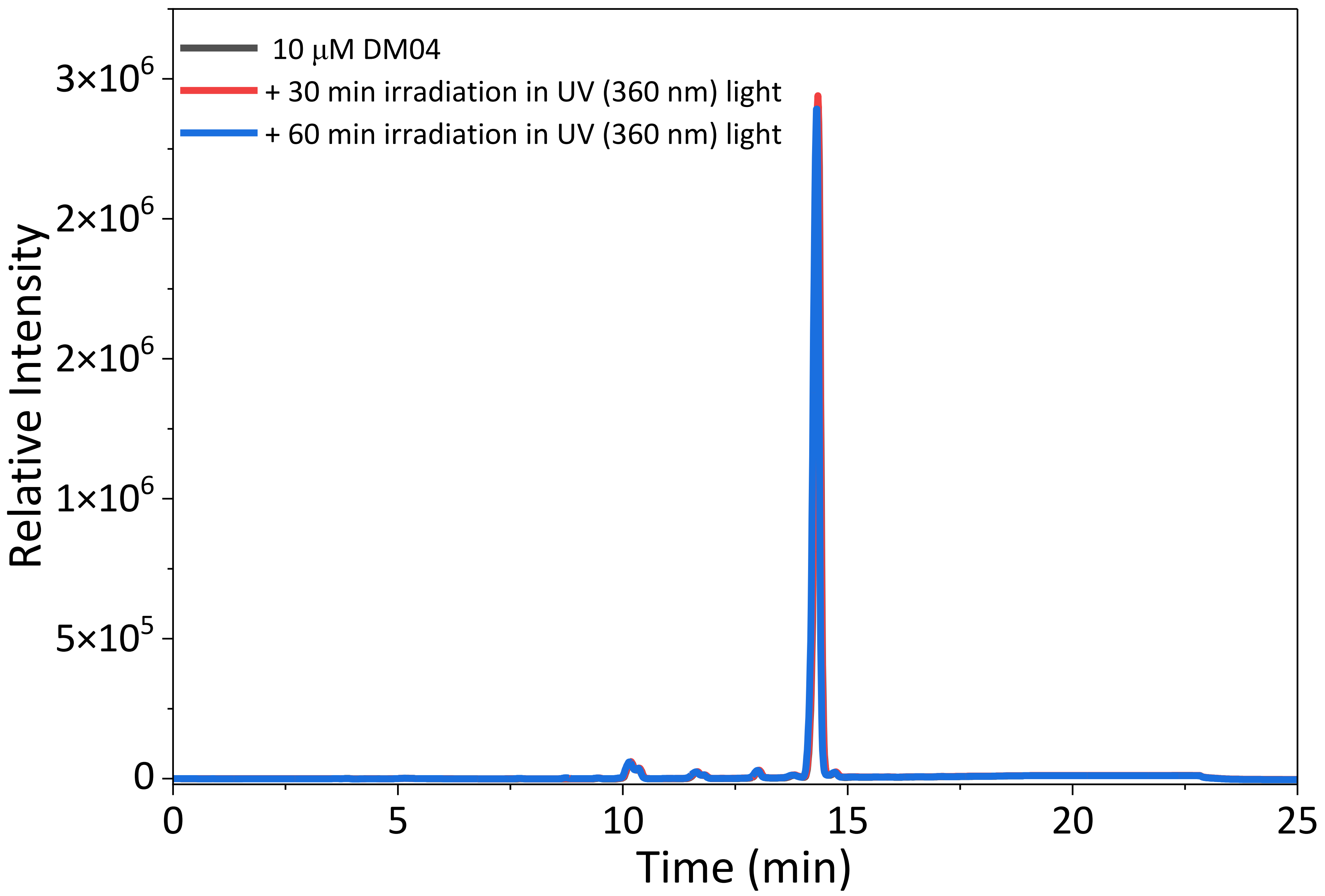

B

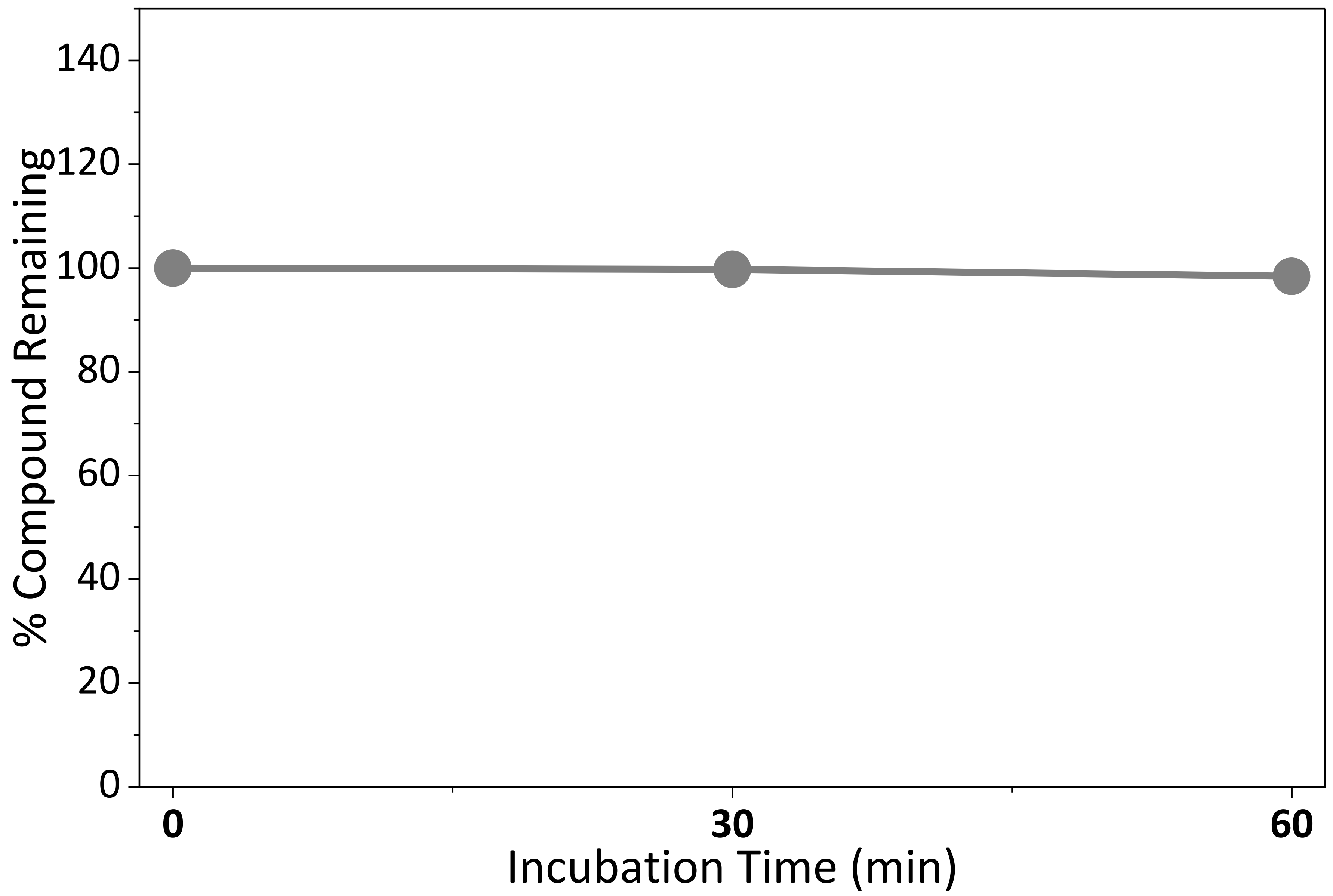

**Figure S3. HPLC analysis demonstrated that stability of 10 μM of DM04 is unchanged after 60 min irradiation of UV light (360 nm). (A)** HPLC chromatogram plot **(B)** Quantification of % DM04 remaining at different time points following UV light irradiation

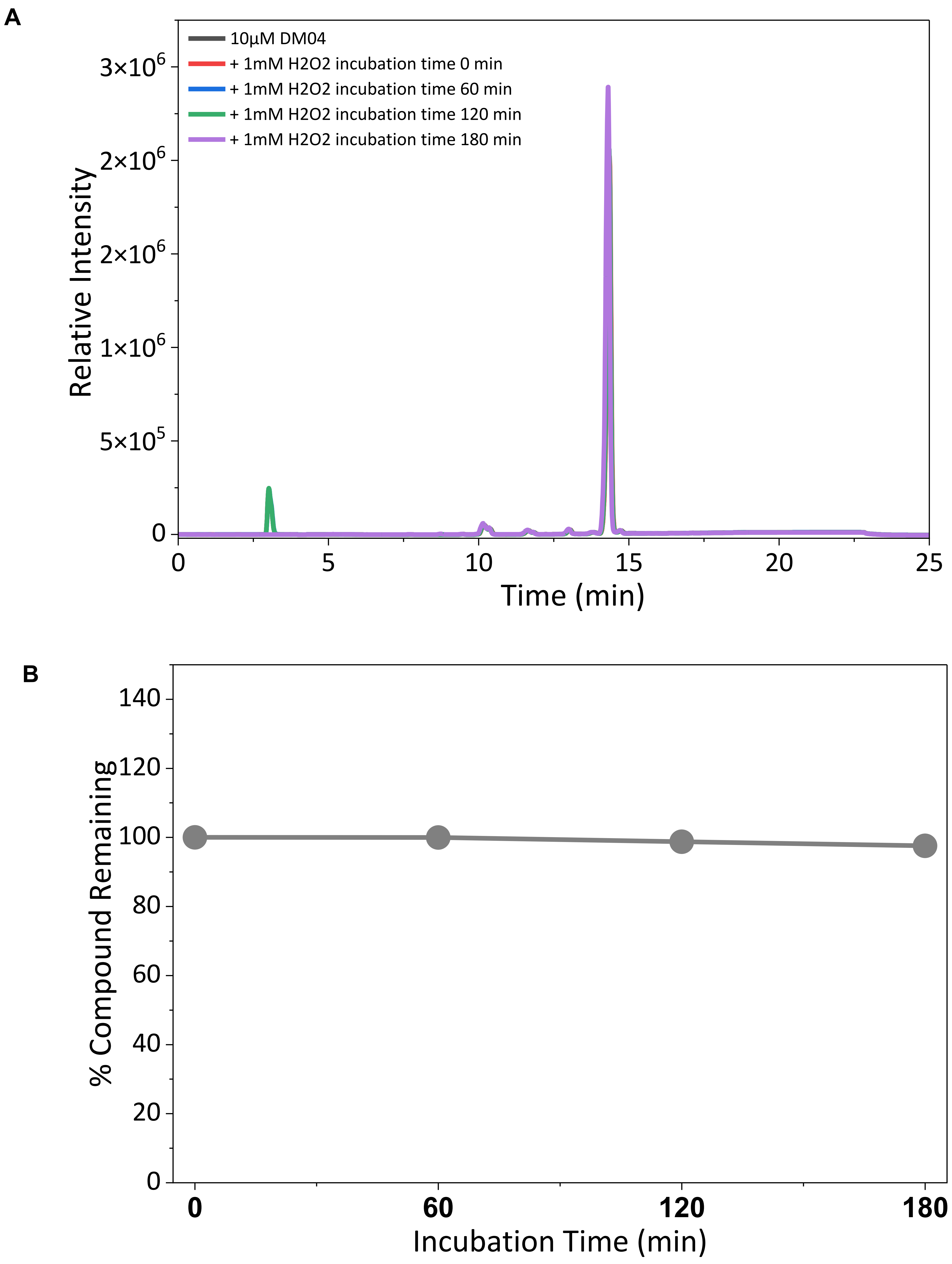

**Figure S4. HPLC analysis demonstrated that stability of 10 µM of DM04 is unchanged after 1mM peroxide treatment for 3 hr (A)** HPLC chromatogram plot **(B**) Quantification of % DM04 remaining at different time points following incubation in $H_2O_2$.

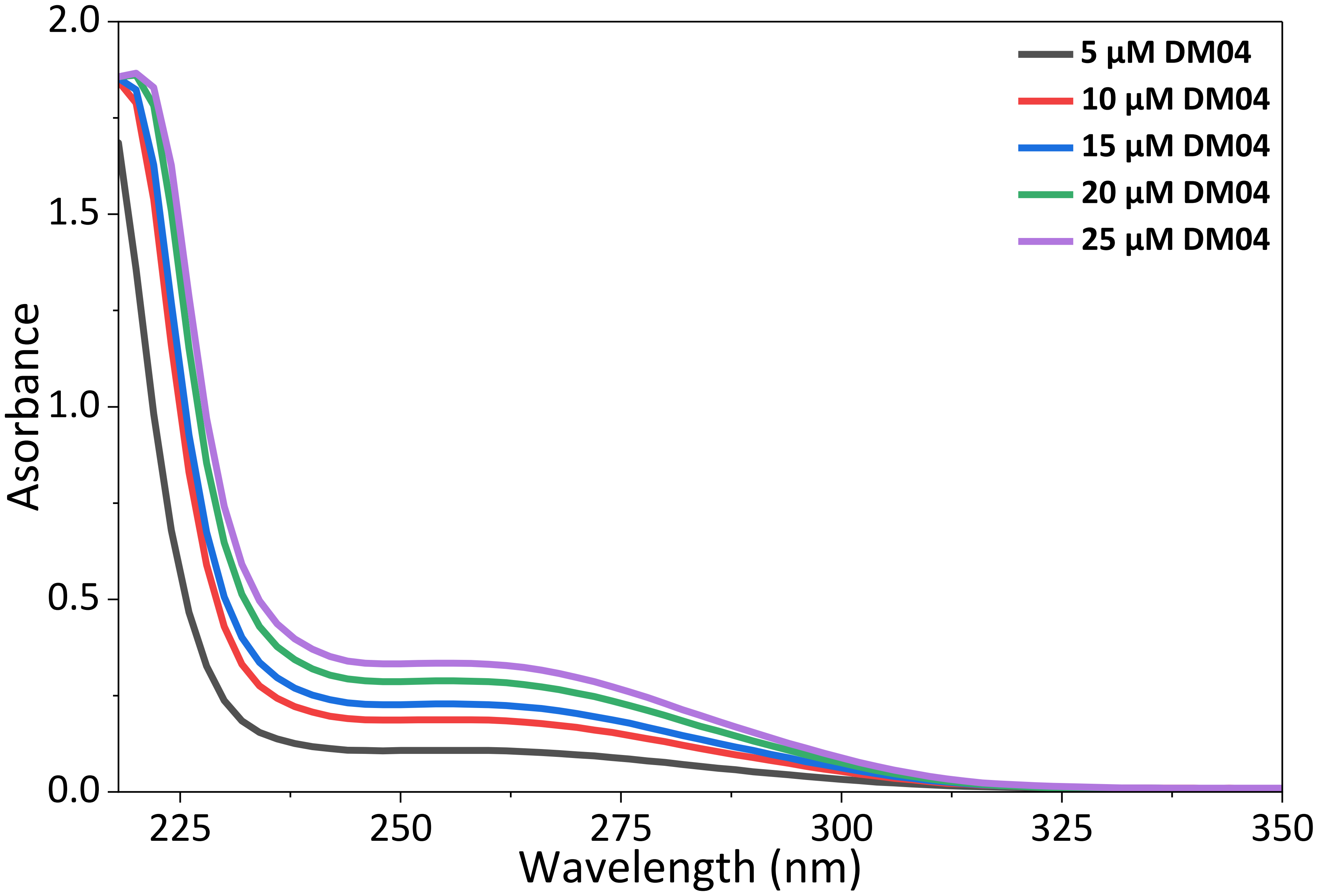

**Figure S5. UV-Vis spectra of DM04 with increasing concentration in water.** UV–Vis absorption spectra of DM04 recorded in water at concentrations ranging from 5 to 25 µM. Absorbance intensity increased in a concentration-dependent manner across the measured wavelength range, with prominent absorption observed in the UV region.

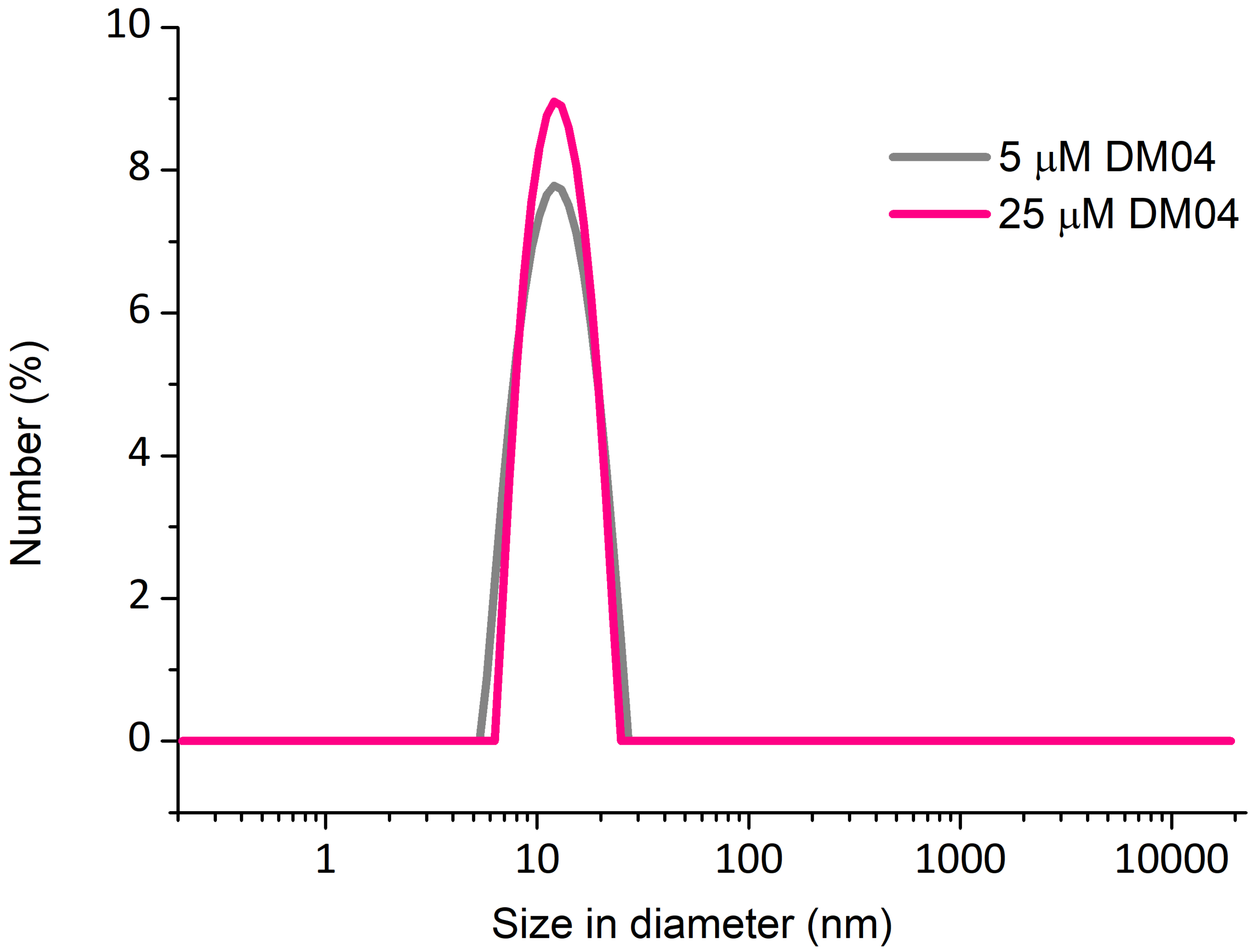

**Figure S6. Size distribution of DM04 at different concentration in water.** Size distribution profiles of DM04 measured at 5 μM and 25 μM in water. The particle size distribution is presented as number (%) as a function of particle diameter (nm).

A

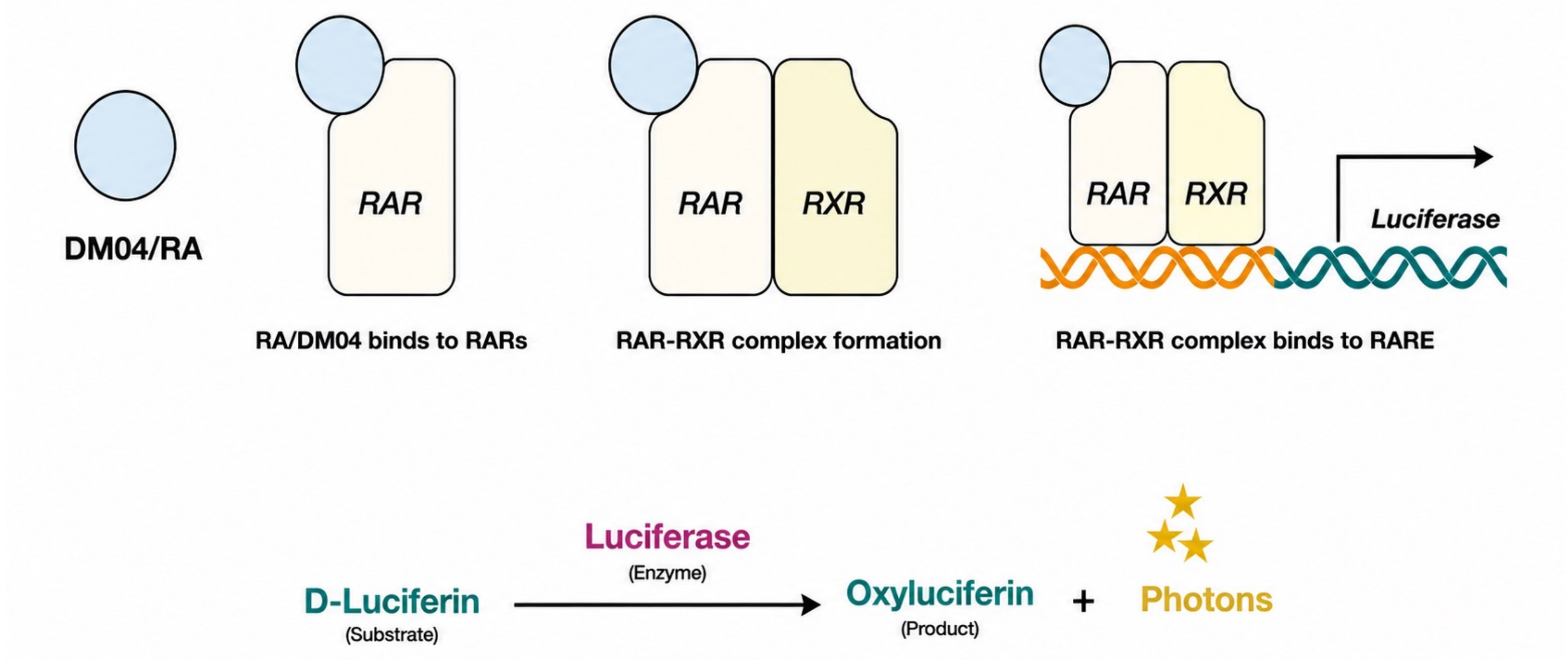

B

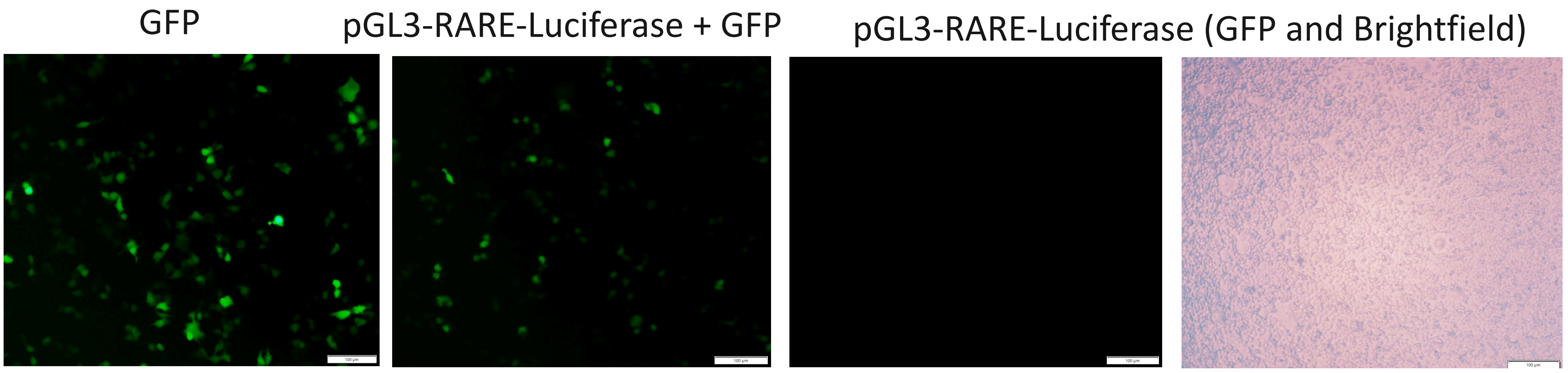

C

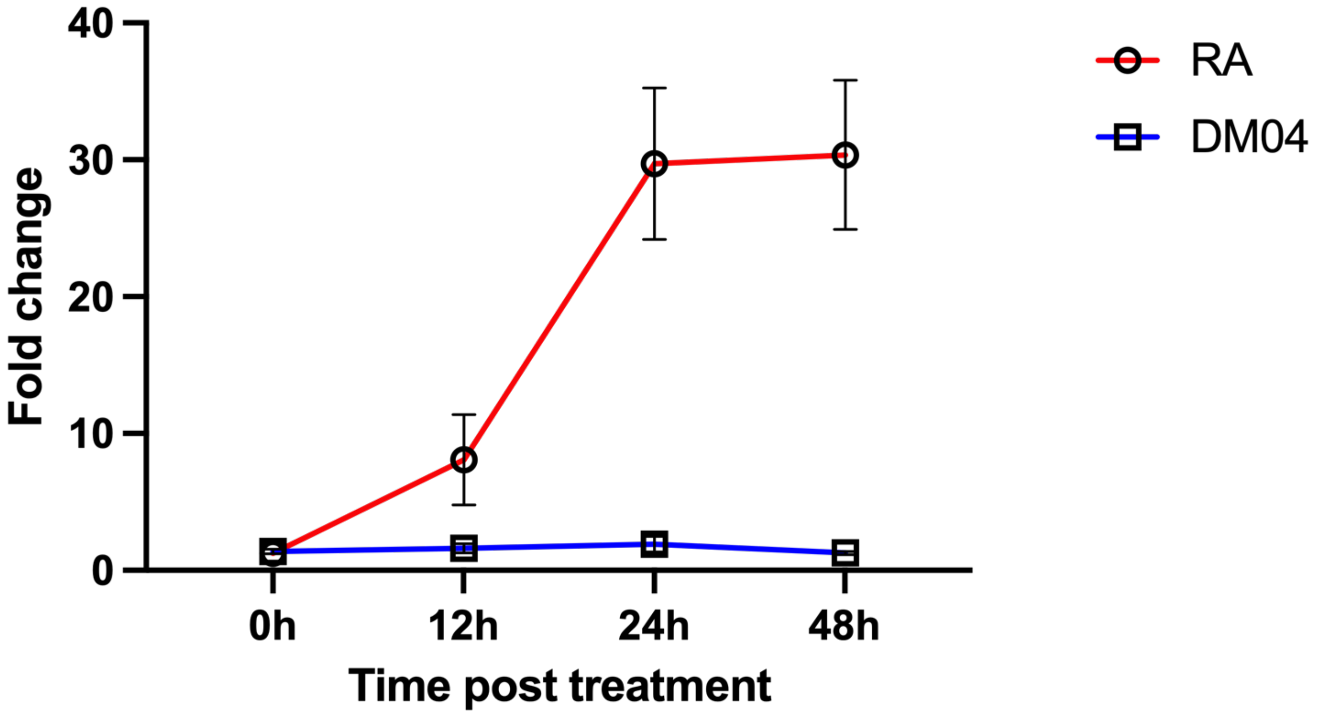

**Figure S7 : DM04 failed to bind to the RAR elements from the pGL3-RARE-Luciferase plasmid.**
**(A)** Schematic overview of the luciferase assay. DM04 is expected to bind to the RAR, form complex with RXR and bind to the RARE elements in the pGL3-RARE-Luciferase plasmid, and there by activate luciferase. Upon substrate (D-luciferin) treatment, photons get released and can be detected in the luminometer. **(B)** Representative images of control pAAV-CAG-GFP transfection, and co-transfection of pGL3-RAR-Luc and pAAV-CAG-GFP and brightfield images of transfected N2A with pGL3-RARE-Luc plasmid. **(C)**Log2Fold change of luciferase activity plotted for RA and DM04 treatment. Data are presented as mean ± SEM from three independent biological replicates. Statistical analysis was performed using a two-way ANOVA followed by Šídák's multiple-comparison test. Significant effects of treatment ($F(1,4) = 288.8$, $\mathbf{P < 0.0001}$), time ($F(1.04,4.16) = 9.85$, $\mathbf{P = 0.0326}$), and treatment × time interaction ($F(3,12) = 9.67$, $\mathbf{P = 0.0016}$) were observed. Šídák's post hoc comparisons did not reveal significant differences between treatments at individual time points in DM04 samples.

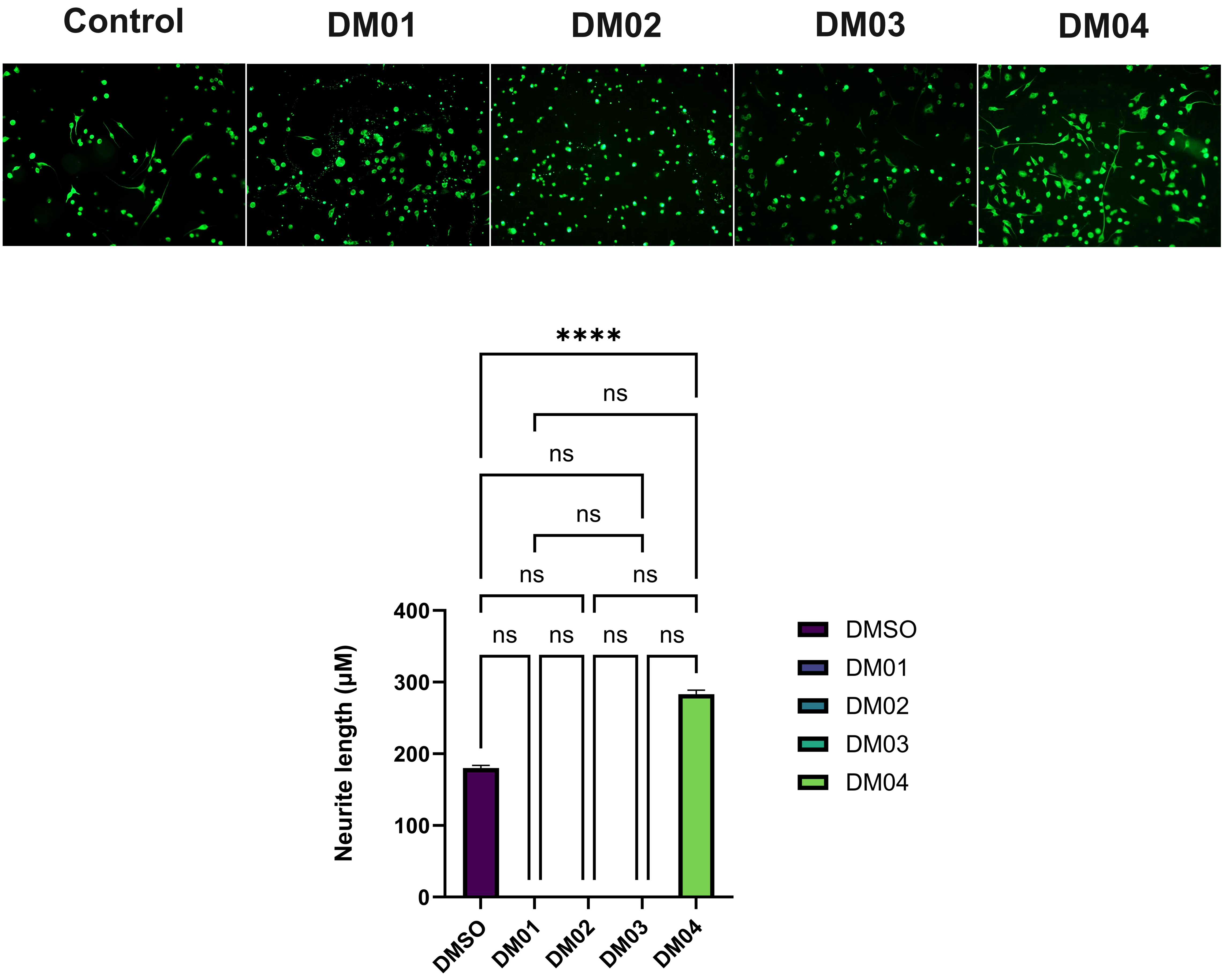

**Figure S8. DM04 significantly increases neurite outgrowth *in vitro* in N2A cells.** Representative fluorescence images showing N2A cells treated with DMSO (control) and DM01–DM04. Quantification of neurite length demonstrates a significant increase following DM04 treatment compared with DMSO and the other DM compounds. Data are presented as mean ± SEM. Statistical significance was determined using the Kruskal–Wallis test with multiple-comparison analysis; ns, not significant; ****P < 0.0001.

### Supplementary Data 5

**[1]H NMR spectra of Compound-1 in $CDCl_3$**

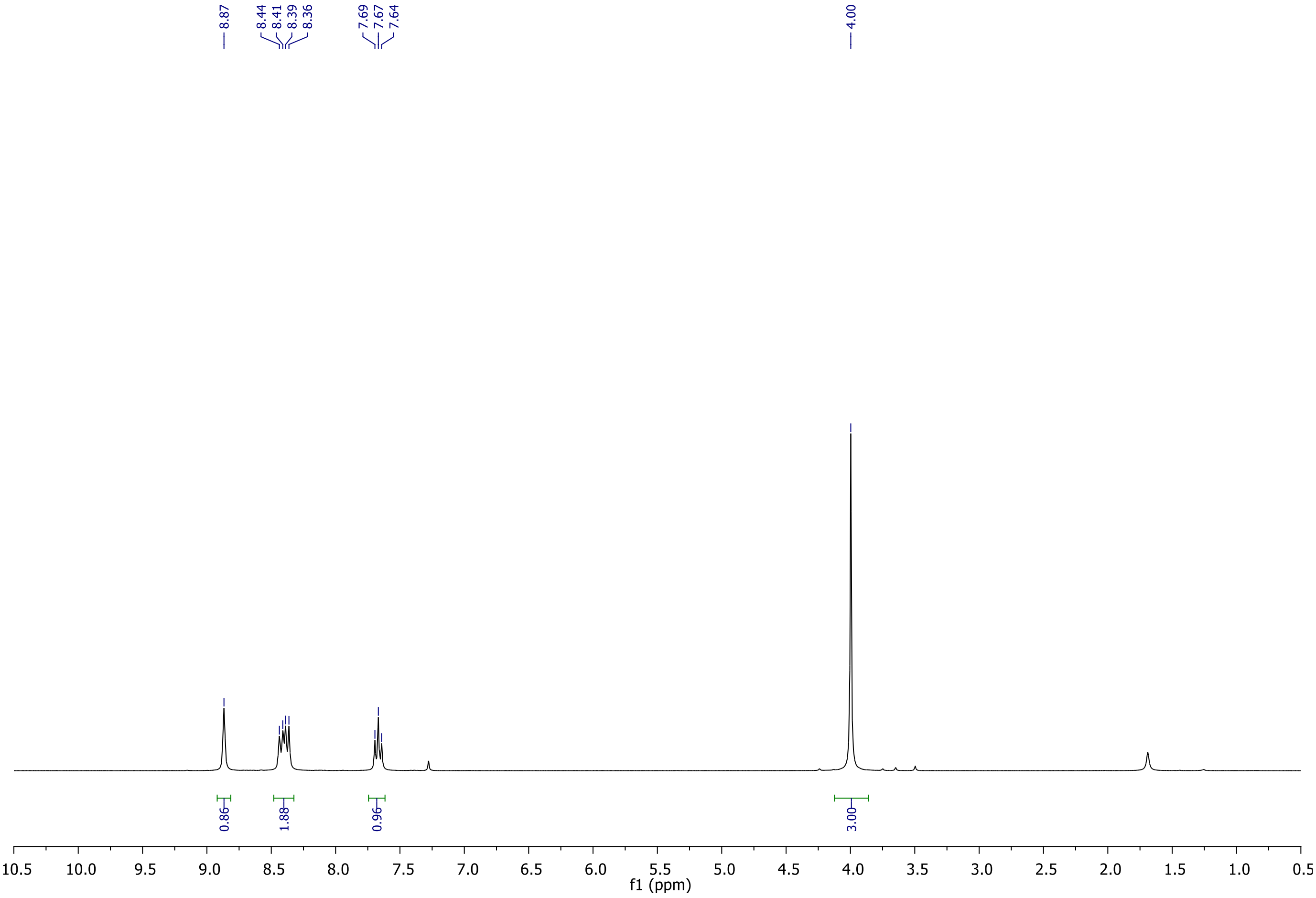

#### $^{13}C$ NMR spectra of Compound-1 in $CDCl_3$

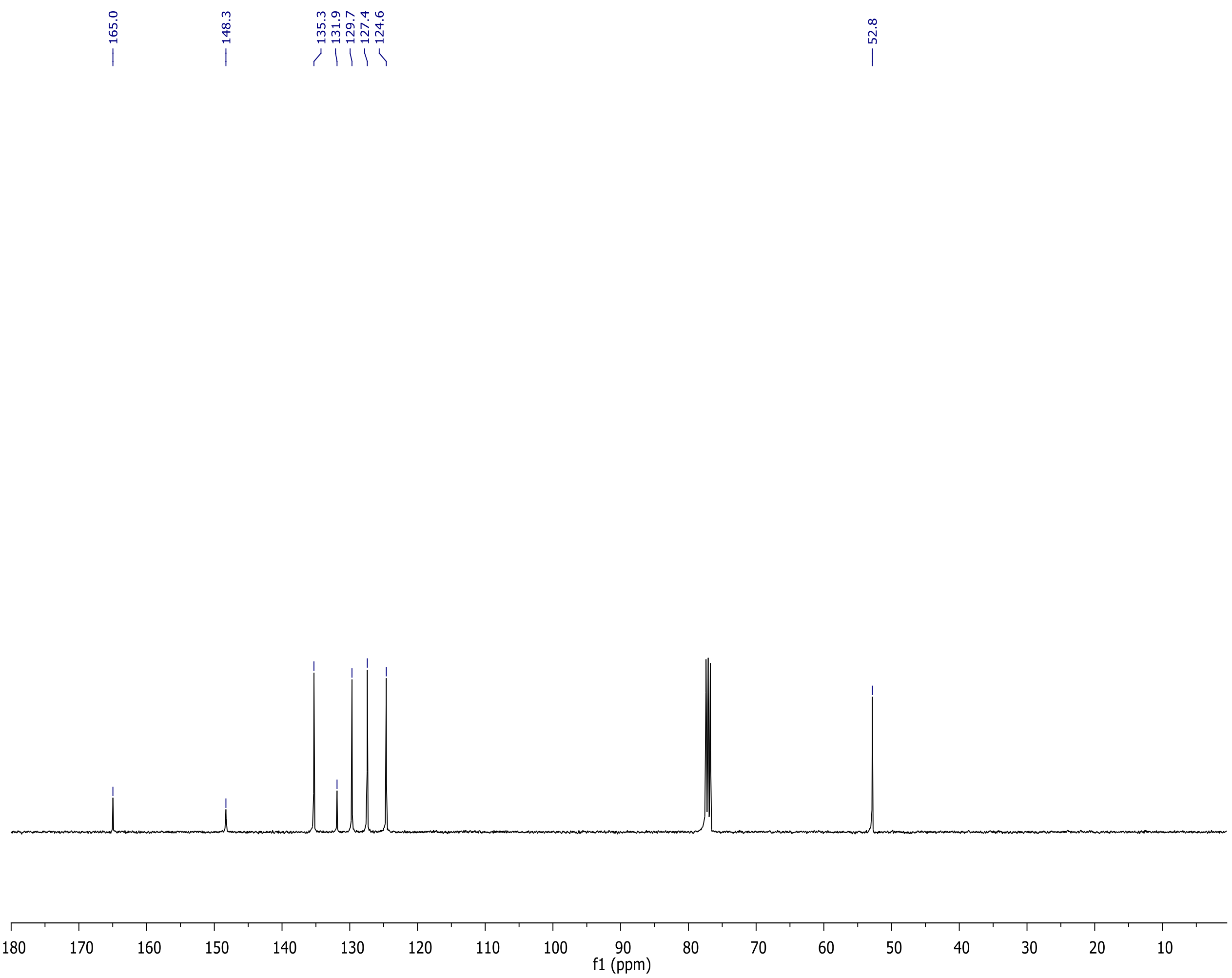

#### $^1$H NMR spectra of Compound-3 in $CDCl_3$

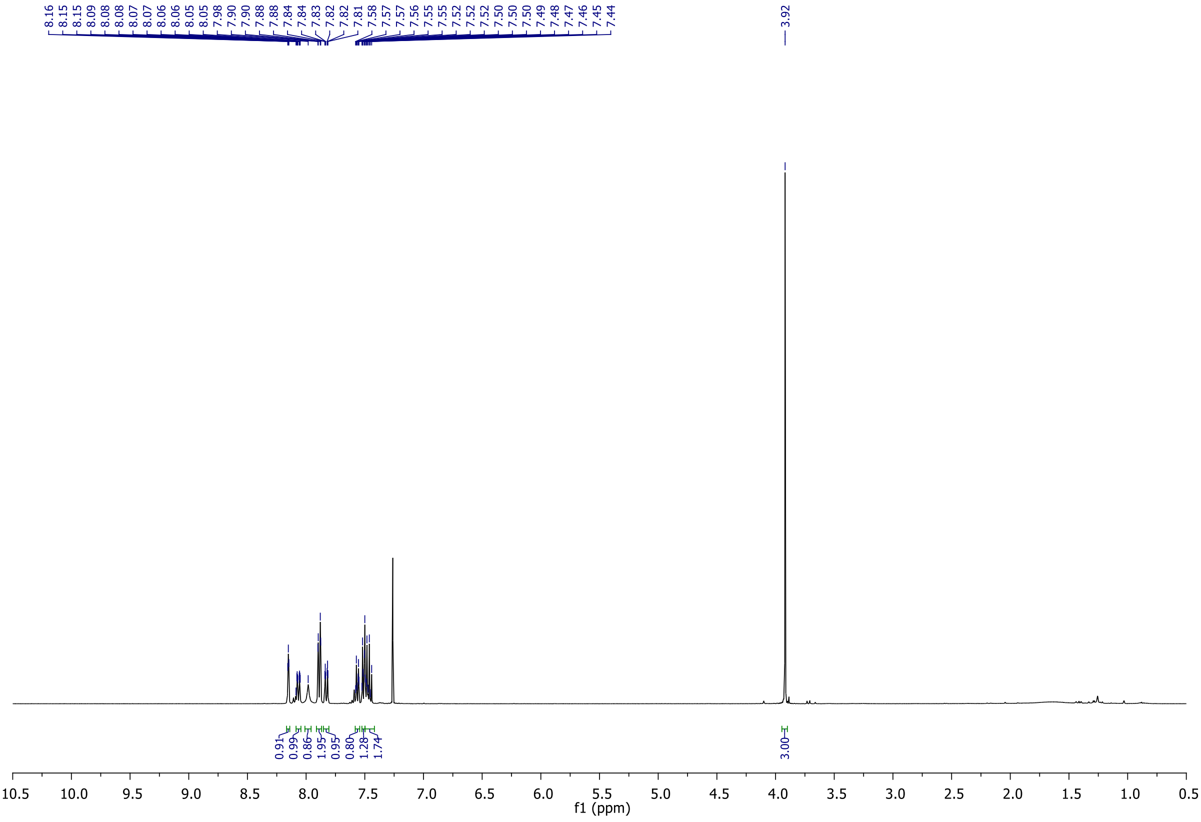

#### $^{13}C$ NMR spectra of Compound-3 in $CDCl_3$

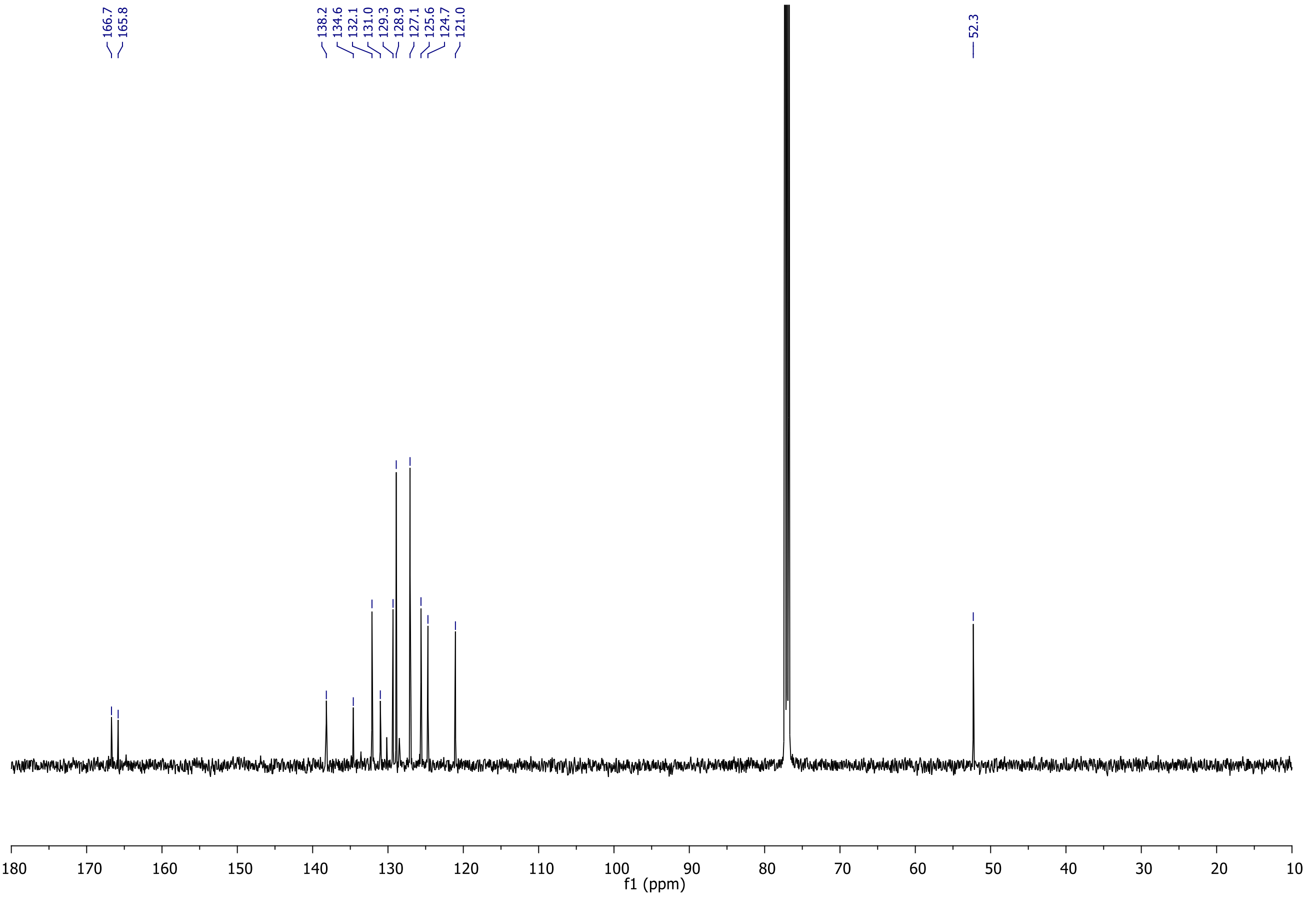

#### HRMS spectrum of Compound-3

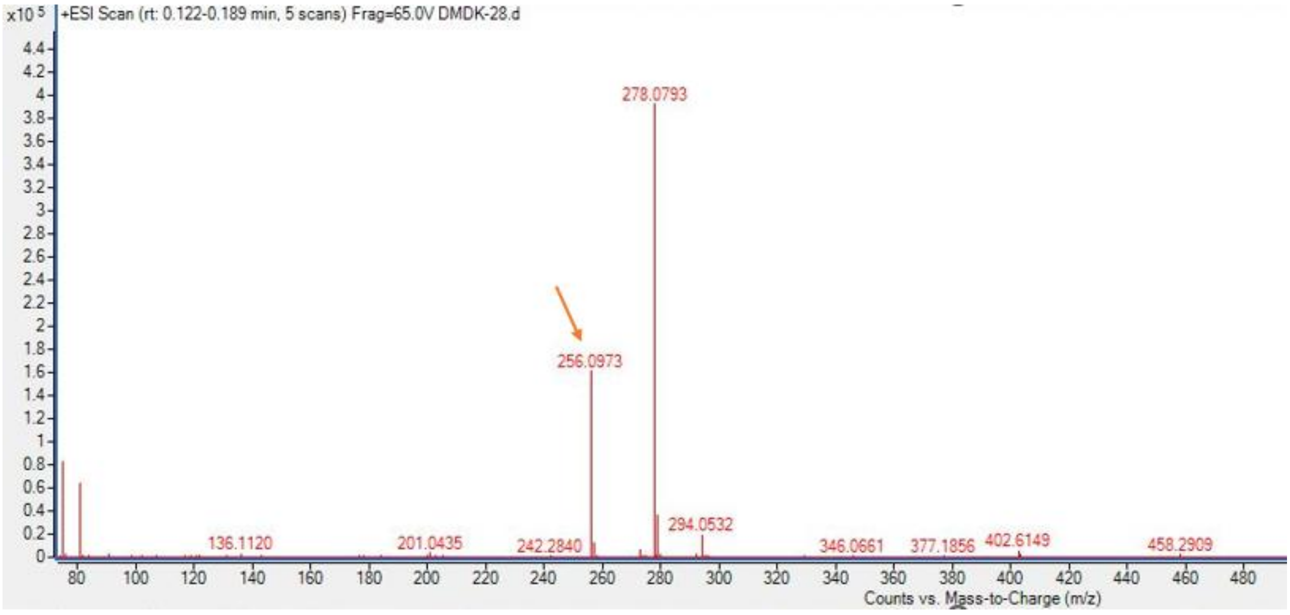

Calculated for $C_{15}H_{14}NO_3$ $(M+H)^+$: 256.0974, found 256.0973.

#### $^1$H NMR spectra of DM-01 in DMSO-$d_6$

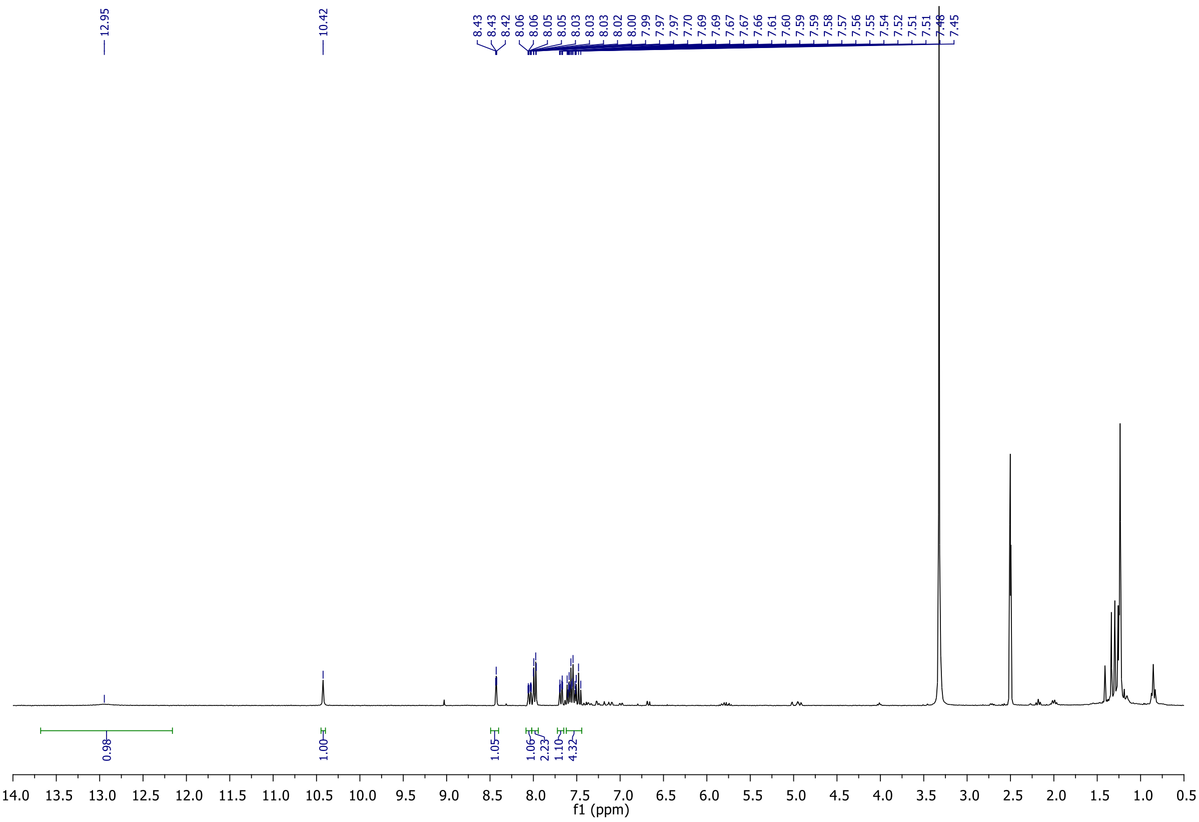

**$^{13}$C NMR spectra of DM-01 in DMSO-$d_6$**

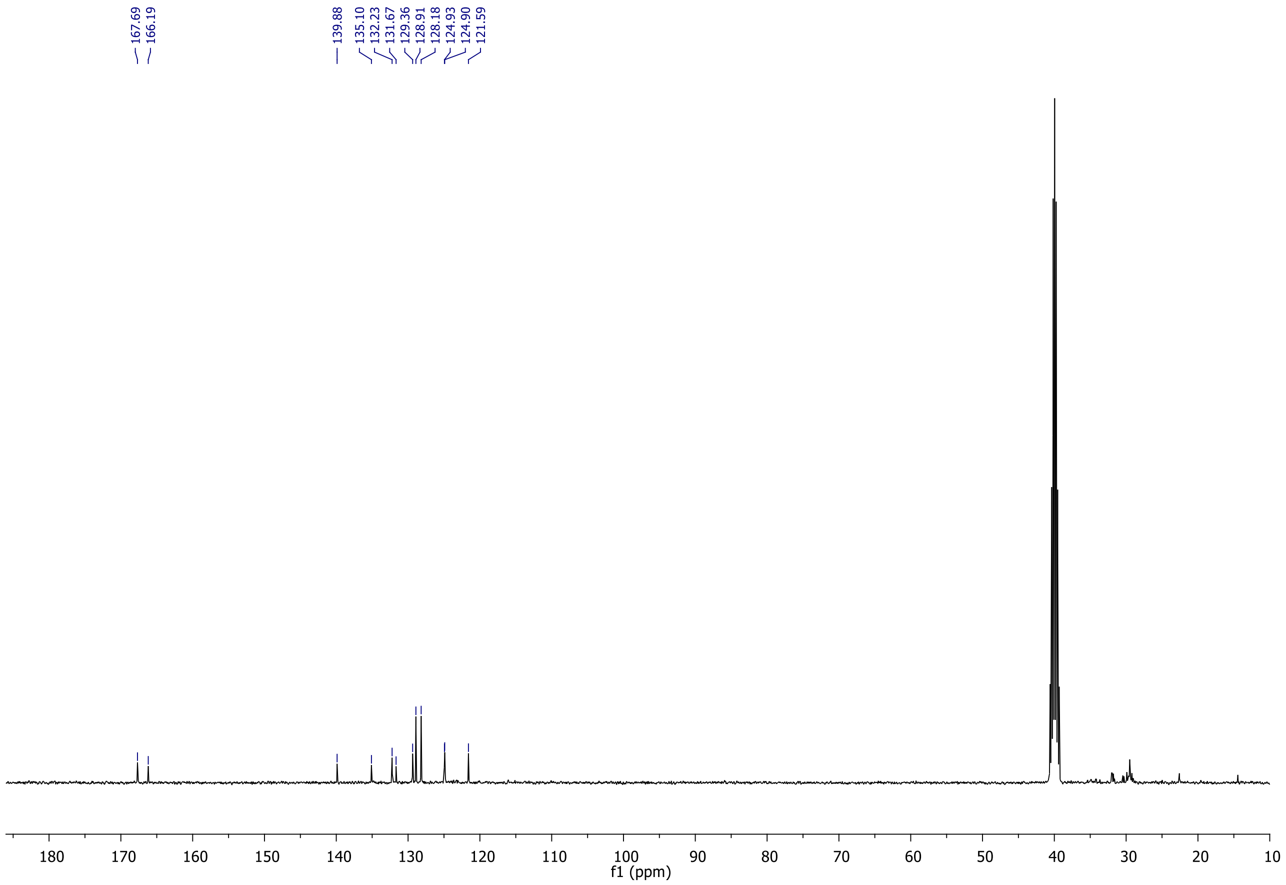

**HRMS spectrum of DM-01**

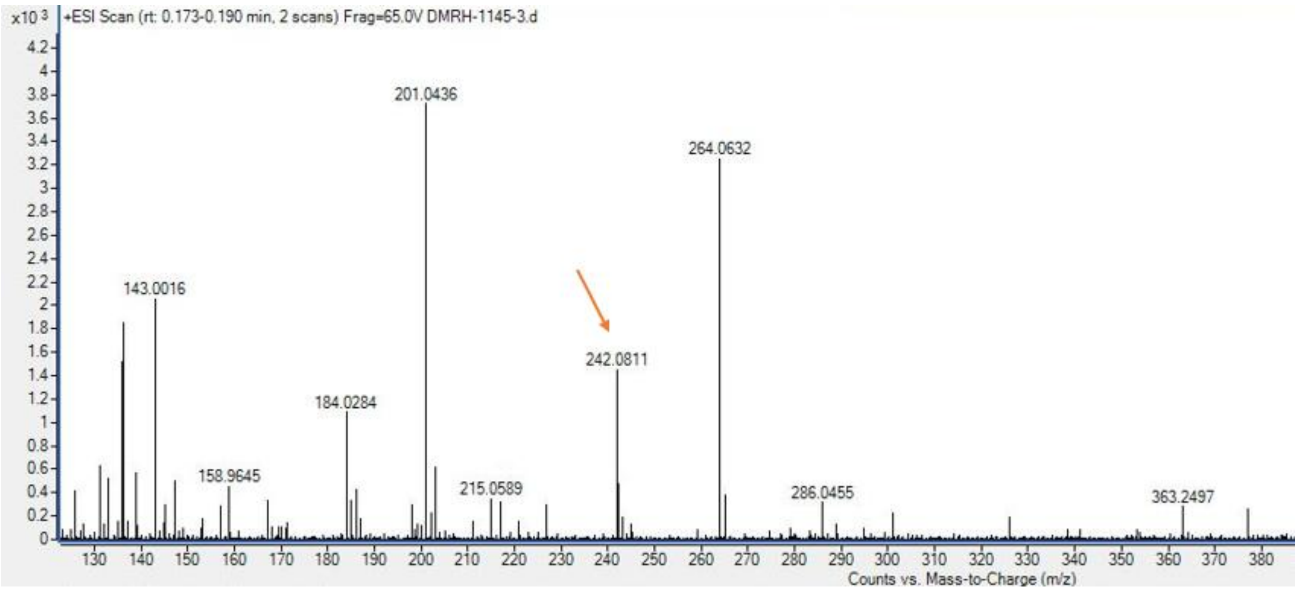

Calculated for $C_{14}H_{12}NO_3$ $(M+H)^+$: 242.0817, found 242.0811.

#### $^1$H NMR spectra of Compound-4 in $CDCl_3$

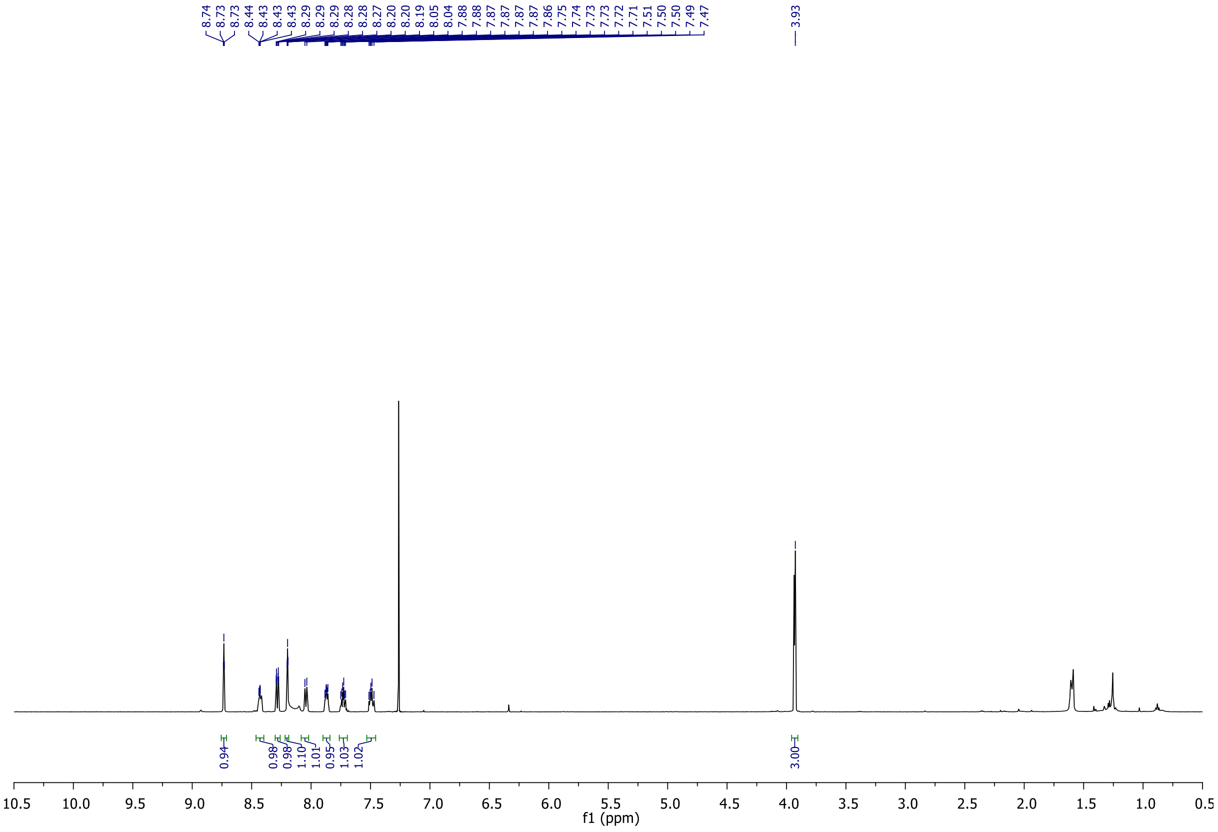

**$^{13}C$ NMR spectra of Compound-4 in $CDCl_3$**

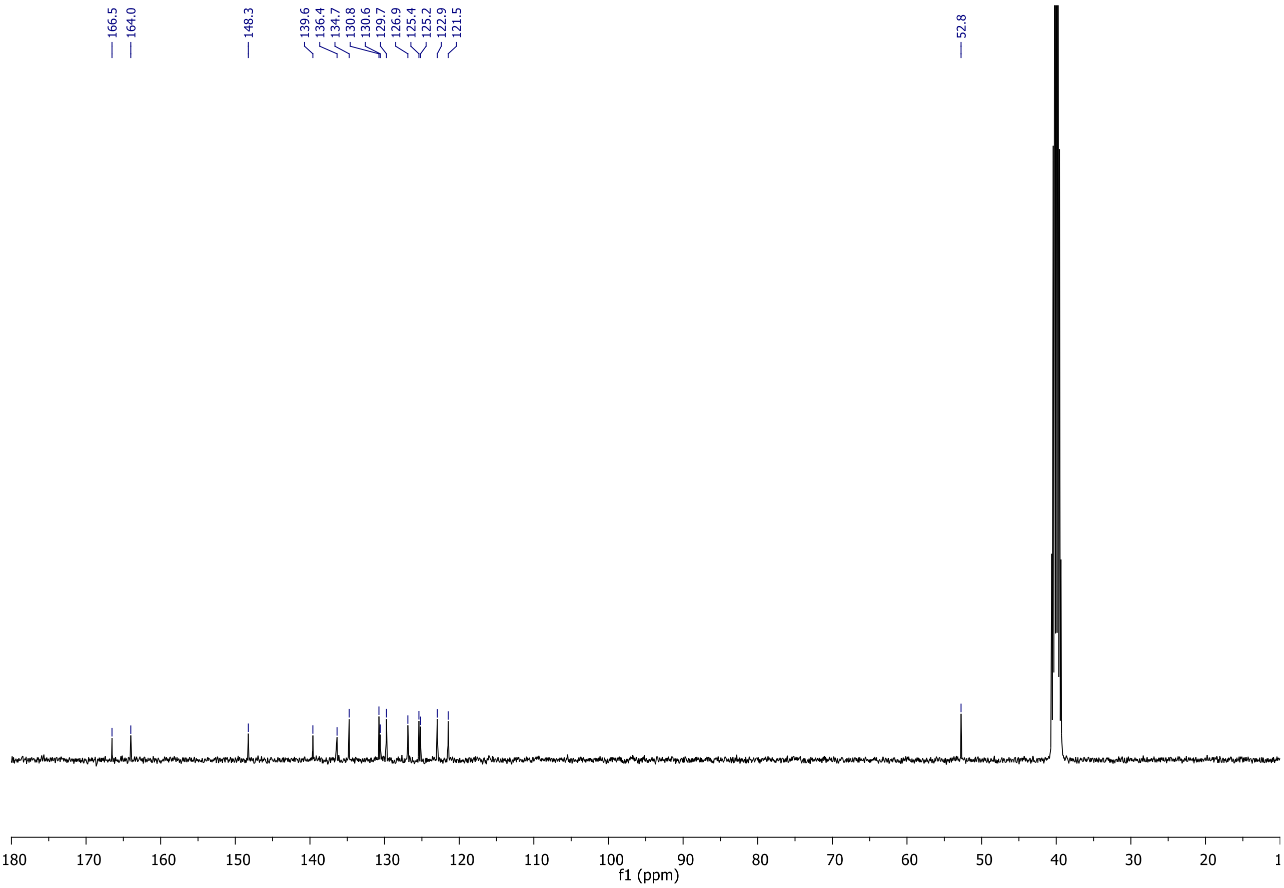

**HRMS spectrum of Compound-4**

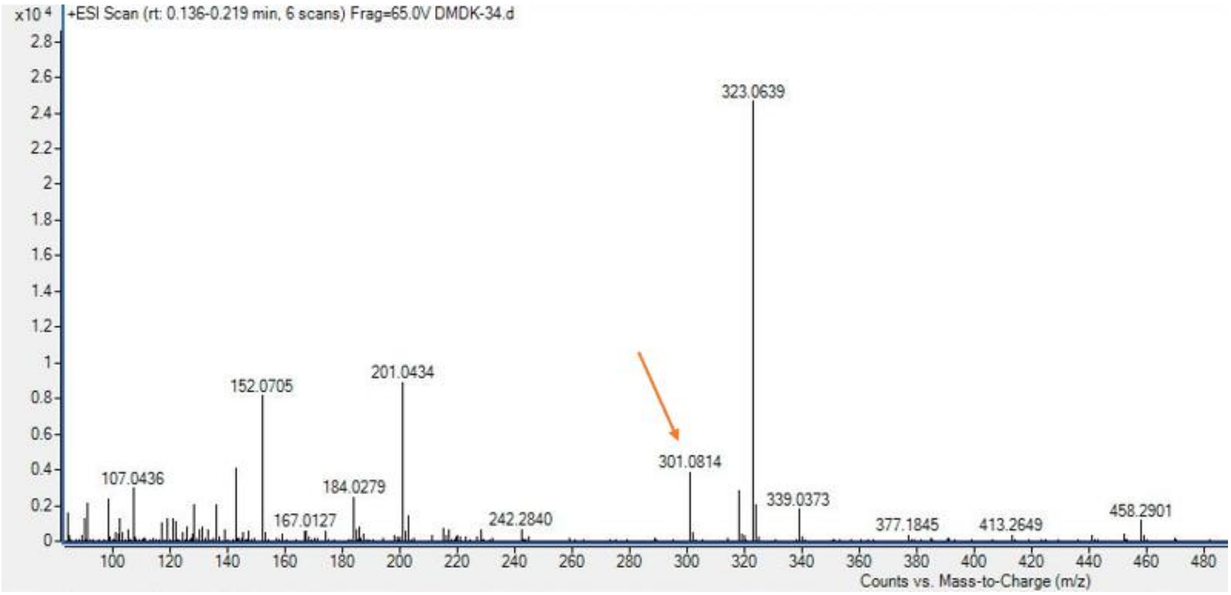

Calculated for $C_{15}H_{13}N_2O_5$ $(M+H)^+$: 301.0824, found 301.0814.

#### $^{1}H$ NMR spectra of DM-02 in DMSO-$d_6$

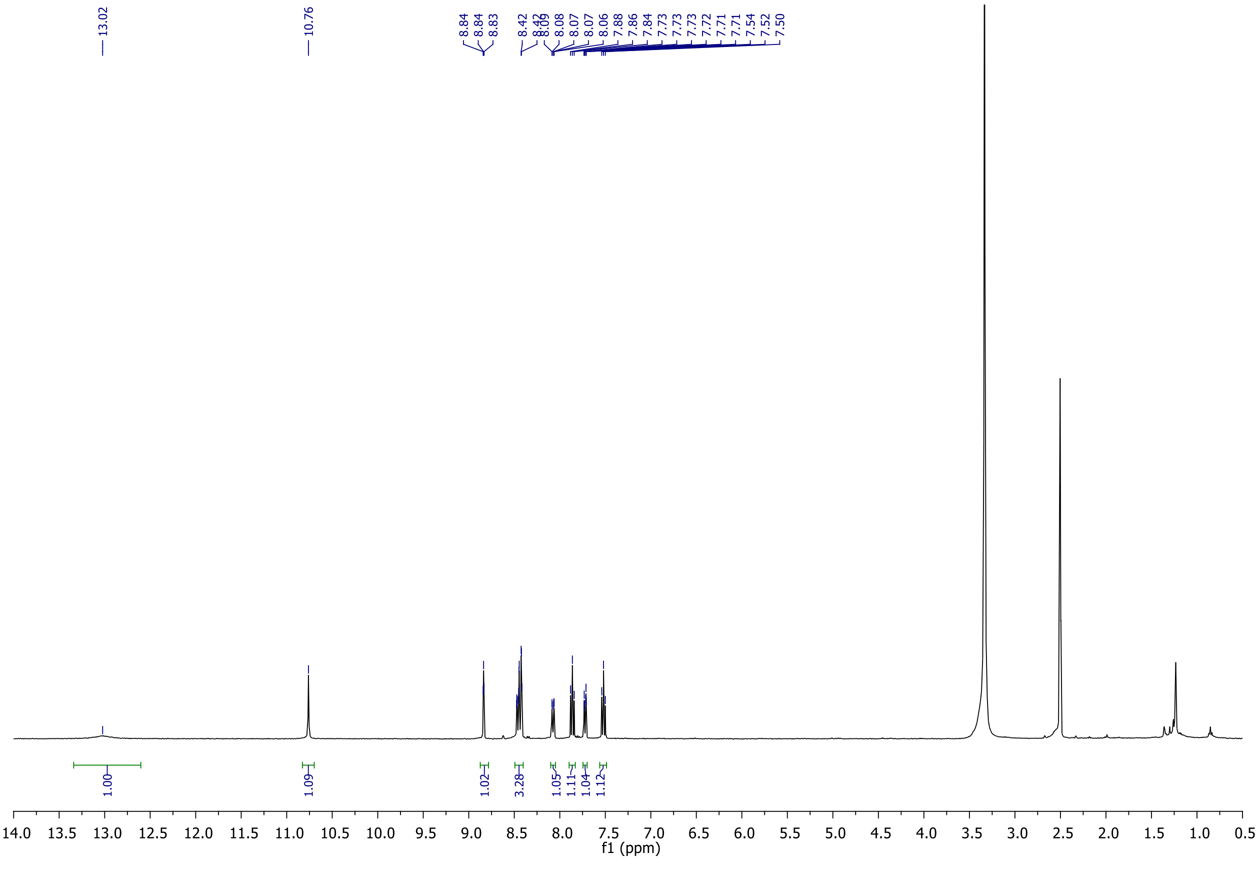

**$^{13}C$ NMR spectra of DM-02 in DMSO-$d_6$**

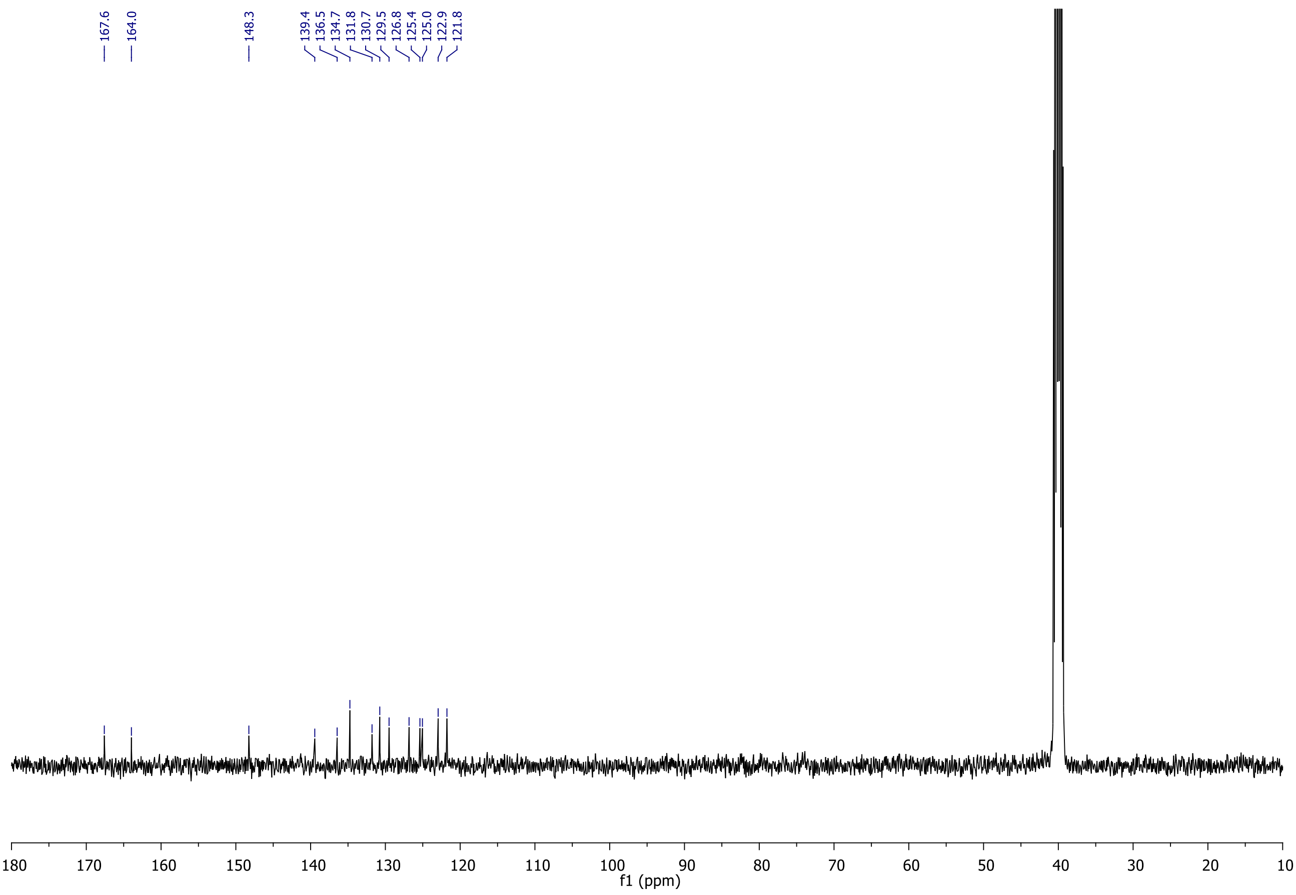

**HRMS spectrum of DM-02**

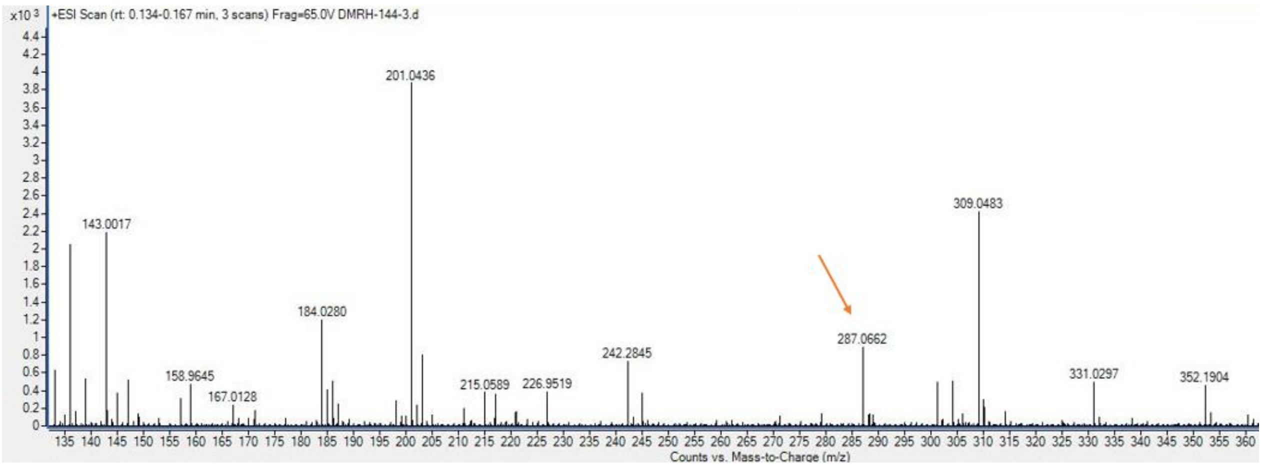

Calculated for $C_{14}H_{11}N_2O_5$ $(M+H)^+$: 287.0668, found 287.0662.

**$^{1}$H NMR spectra of DM-03 in DMSO-$d_6$**

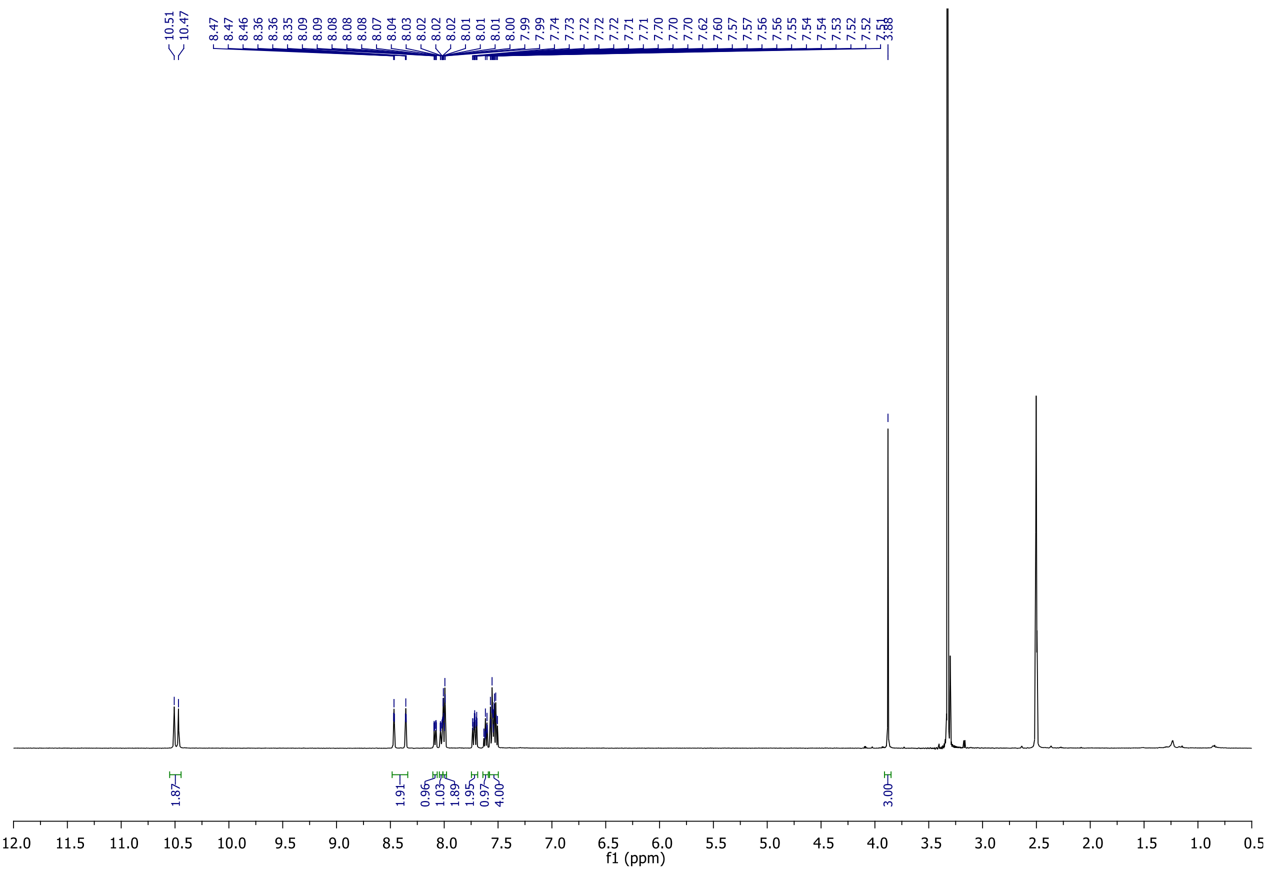

**$^{13}C$ NMR spectra of DM-03 in DMSO-$d_6$**

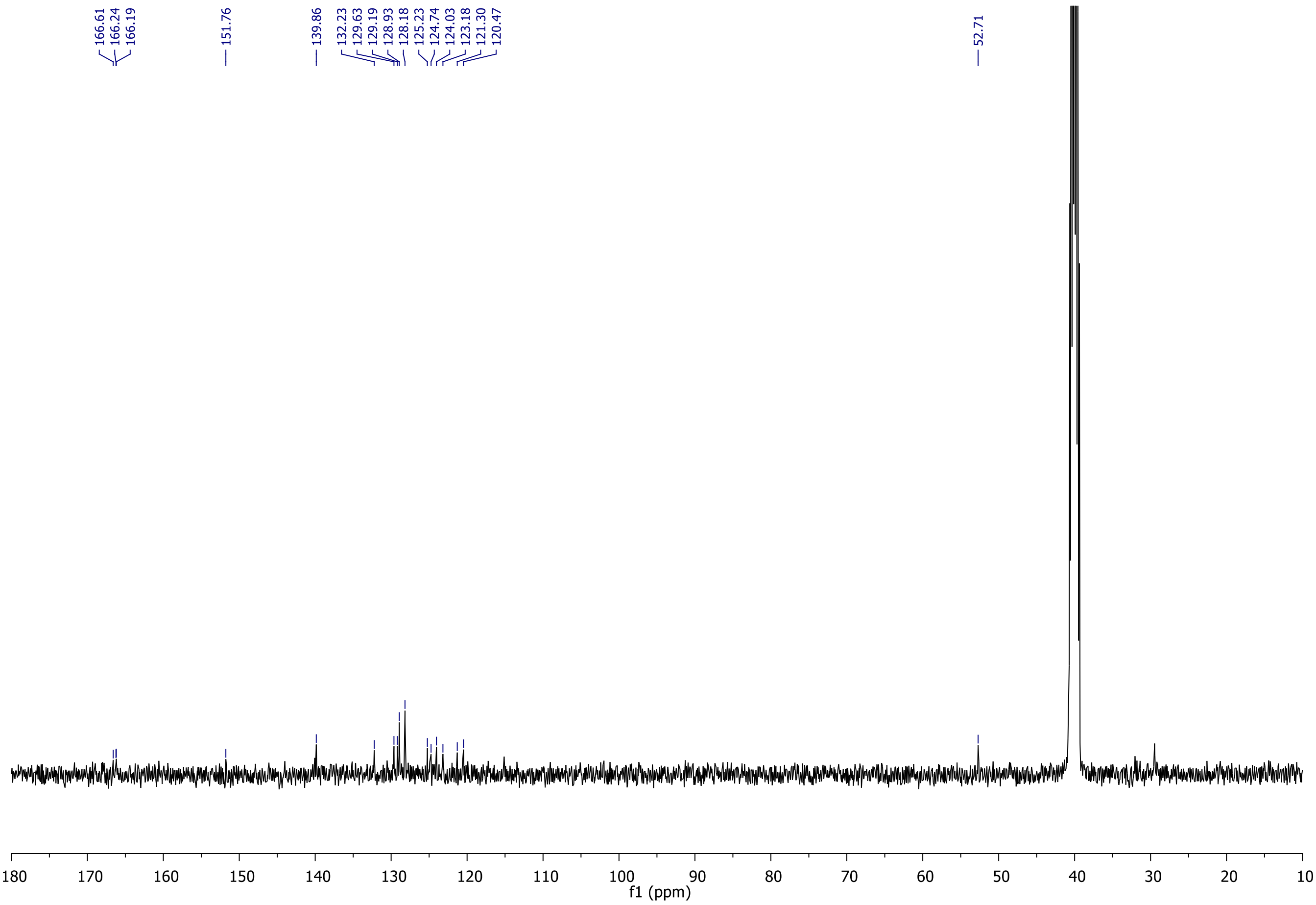

**HRMS spectrum of DM-03**

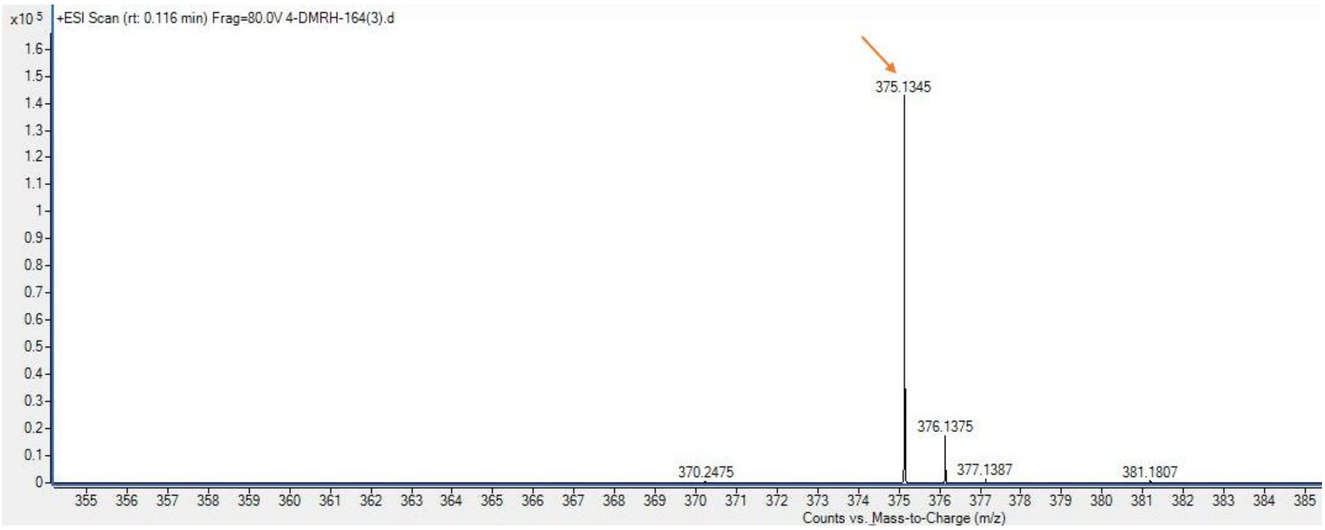

Calculated for $C_{22}H_{19}N_2O_4$ $(M+H)^+$: 375.1345, found 375.1345.

#### $^1$H NMR spectra of DM-04 in DMSO-$d_6$

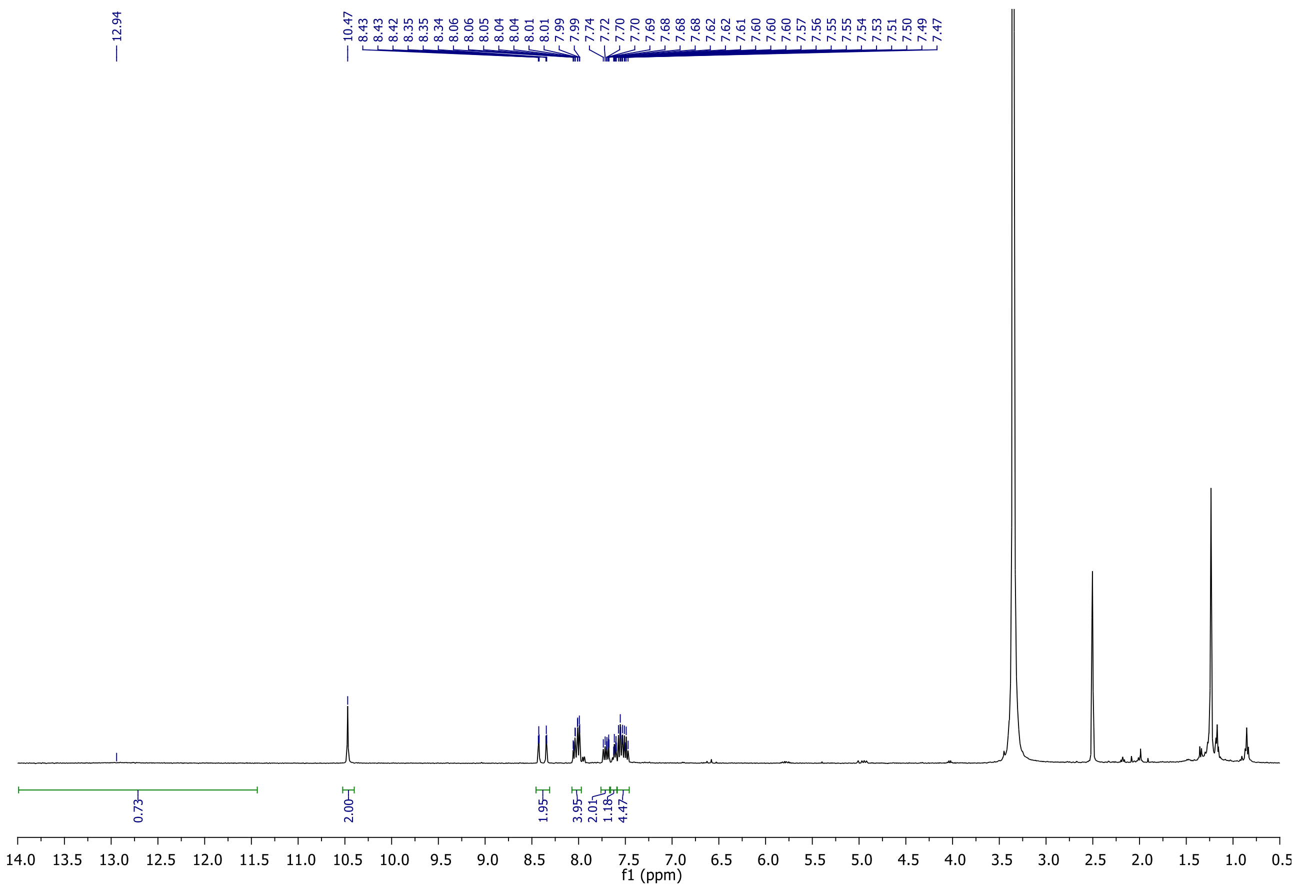

#### $^{13}C$ NMR spectra of DM-04 in DMSO-$d_6$

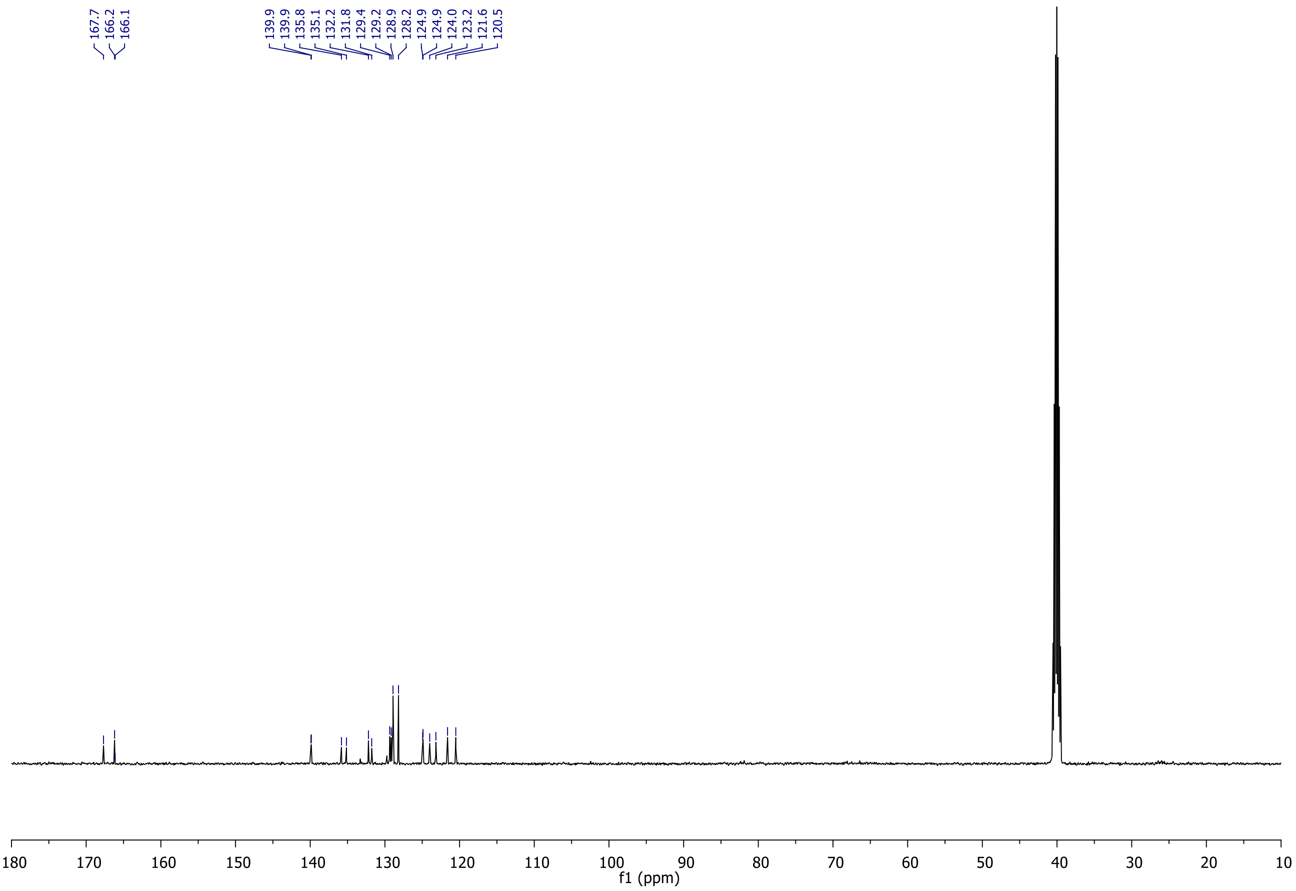

#### HRMS spectrum of DM-04

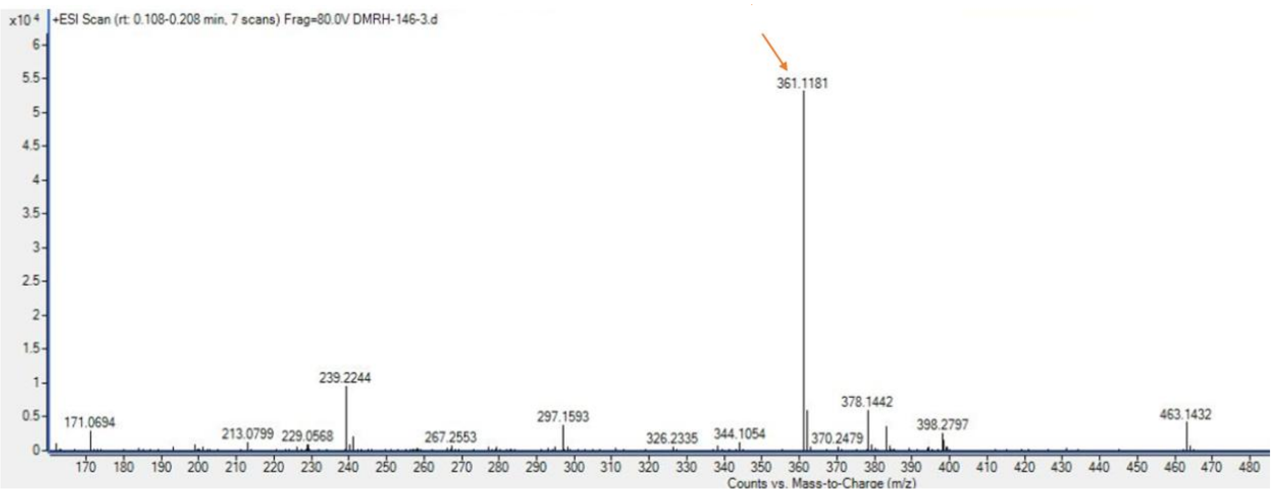

Calculated for $C_{21}H_{17}N_2O_4$ $(M+H)^+$: 361.1188, found 361.1181.

### Supplementary Data 6

#### Maity Lab CSIR-IICT

##### <Sample Information>

| | | | |
|---|---|---|---|
| Sample Name | : DMRH-04 | | |
| Sample ID | : DMRH-04 | | |
| Data Filename | : DMRH-04-Purity.lcd | | |
| Method Filename | : Purity analysis10-90h.lcm | | |
| Batch Filename | : DMRH-04.lcb | | |
| Vial # | : 1-4 | Sample Type | : Unknown |
| Injection Volume | : 3 uL | | |
| Date Acquired | : 11/12/2025 4:17:23 PM | Acquired by | : System Administrator |
| Date Processed | : 11/12/2025 4:44:51 PM | Processed by | : System Administrator |

##### <Chromatogram>

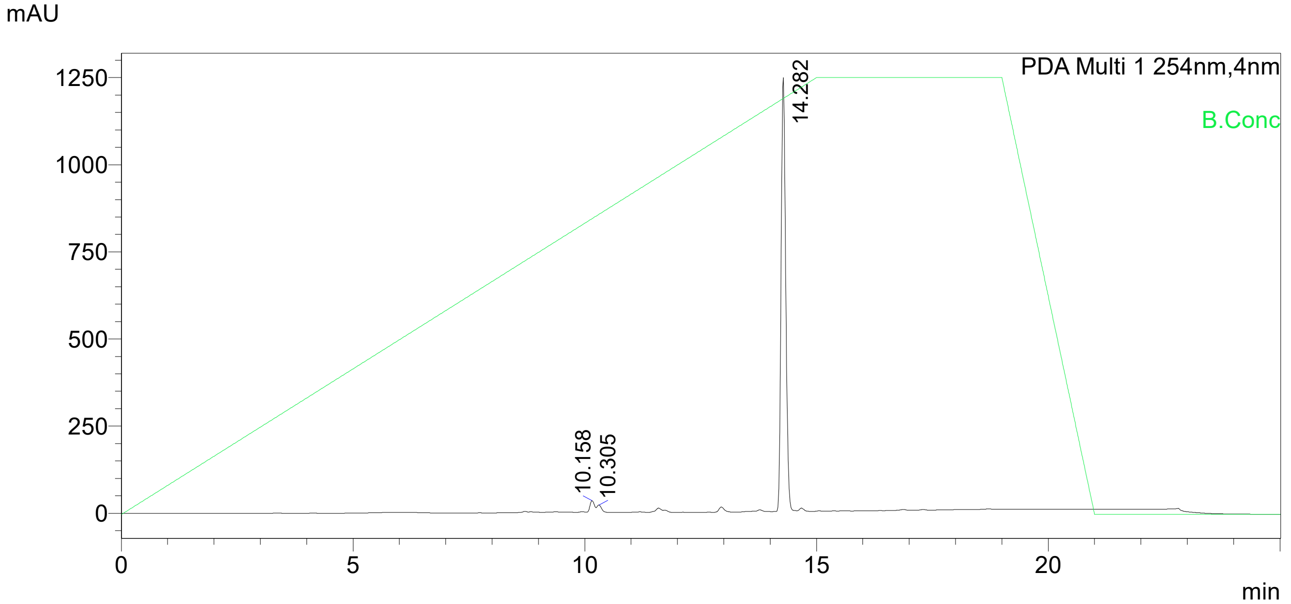

UV Spectrum

Peak# : 3
Retention Time : 14.282 min
Compound Name :
Spectrum Operation : None

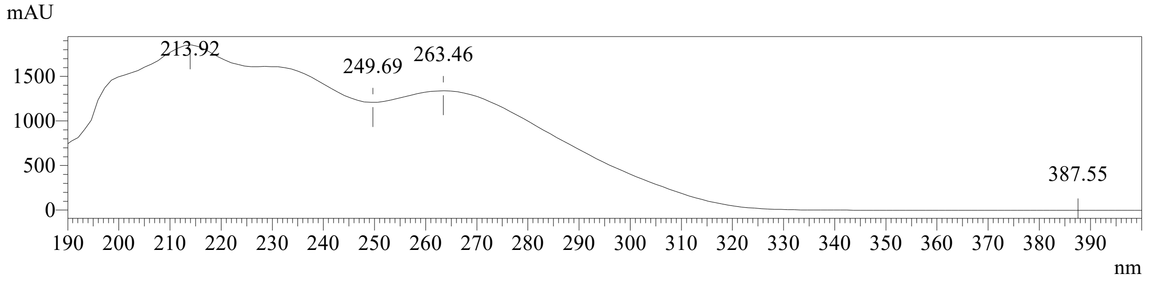

Peak Table

PDA Ch1 254nm

| Peak# | Ret. Time | Area | Height | Area% |
|---|---|---|---|---|
| 1 | 10.158 | 145158 | 25689 | 1.662 |
| 2 | 10.305 | 58965 | 11704 | 0.675 |
| 3 | 14.282 | 8528631 | 1243661 | 97.663 |
| Total | | 8732753 | 1281053 | 100.000 |
